# Insect-habitat-plant interaction networks provide guidelines to mitigate the risk of transmission of *Xylella fastidiosa* to grapevine in Southern France

**DOI:** 10.1101/2024.06.23.600284

**Authors:** Xavier Mesmin, Marguerite Chartois, Pauline Farigoule, Christian Burban, Jean-Claude Streito, Jean-Marc Thuillier, Éric Pierre, Maxime Lambert, Yannick Mellerin, Olivier Bonnard, Inge van Halder, Guillaume Fried, Jean-Yves Rasplus, Astrid Cruaud, Jean-Pierre Rossi

## Abstract

1. *Xylella fastidiosa* (*Xf*) is a xylem-limited bacterium that has been recorded in several European countries since its detection in 2013 in Apulia (Italy). Given the prominence of the wine industry in many southern European countries, a big threat is the development of Pierce’s disease in grapevines. Yet, the insect-habitat and insect-plant interaction networks in which xylem feeders, possible vectors of *Xf,* are involved around European vineyards are largely unknown.
2. Here we describe these networks in three important wine-growing regions of southern France (Provence-Alpes-Côte d’Azur, Occitanie, Nouvelle-Aquitaine) to identify the main source habitats of xylem feeders, and to gather information about their specialization degree at the habitat, plant family, and plant species levels. A total of 92 landscapes (and 700 sites) were studied over three sampling sessions in the fall 2020, the spring 2021, and the fall 2021.
3. Among the habitats sampled, meadows hosted the largest xylem feeder communities, followed by alfalfa fields. Vineyard headlands and inter-rows hosted slightly smaller xylem feeder communities, indicating that potential *Xf* vectors were thriving in the close vicinity of vulnerable plants. Finally woody habitats in general and grapevines in particular hosted very few xylem feeders, showing that transfer on vulnerable plants are rare, but not inexistent. In terms of specialization degrees, *Philaenus spumarius*, *Aphrophora alni, Lepyronia coleoptrata*, and *Cicadella viridis* were all similarly generalists at the habitat, plant family or plant species level. The only specialist was *Aphrophora* grp. *salicina*, which was restricted to riparian forests, and more specifically to Salicaceae. *Neophilaenus* spp. were extremely specialist at the plant family level (Poaceae), but rather generalist at the habitat and plant species levels. All 1017 insects screened for the presence of *Xf* tested negative, showing that *Xf* is not widespread in the studied regions.
4. Our study provides new basic ecological information on potential vectors of *Xf*, especially on their specialization and feeding preferences, as well as practical information that may be relevant for the design of epidemiological surveillance plans.

## Introduction

*Xylella fastidiosa* (*Xf*) (Xanthomonadaceae) is a xylem-limited bacterium transmitted by hemipteran insect vectors (mainly sharpshooters and spittlebugs) (Chatterjee et al., 2008; Krugner et al., 2019) that causes various diseases in a broad range of cultivated and uncultivated plants (Almeida et al., 2019; EFSA et al., 2023). *Xf* originates from the Americas but has been introduced to different regions of the world during the last decades (https://gd.eppo.int/taxon/XYLEFA/distribution).

Among the three described subspecies of *Xf*, *X. fastidiosa* subsp. *fastidiosa* is the causal agent of Pierce’s disease in grapevines, which causes millions of losses to the US grape industry every year (Tumber et al., 2014). In France, the wine and spirits sector is prominent, with annual export profits estimated to exceed 10 billion euros (Durand, 2016). The epidemic risk posed by *Xf* is therefore a legitimate concern for the french wine industry, especially since the subspecies *fastidiosa* has recently been detected in southern Italy (EPPO, 2024; Teatro Naturale, 2024)

When an insect feeds on an infected plant and acquires the bacterium, it can transmit it to the next plant it feeds on, without a latency period (Purcell & Finlay, 1979). The probability of contaminating a healthy crop is thus linked to the connectance of the trophic network composed of the crop, the vectors, and the contaminated plants surrounding the crop. This connectance somewhat reflects vector diet breath in different types of habitat. Therefore, to better assess the risk of contamination by *Xf*, it is essential to decipher habitat-plant-vector interactions at the landscape level around crops.

We still have little knowledge on these networks of interaction. In the US, riparian forests were proven favorable to *Graphocephala atropunctata*, which is responsible for most of the primary transmission of *Xf* to neighboring vineyards (Purcell, 1975). As a consequence, greater damage due to Pierce’s disease was observed near rivers (Purcell, 2013). In Belgium, Casarin et al. (2021) suggested that Salicaceae, abundant on river banks, hosted populations of vectors of *Xf* (especially *Aphrophora* grp. *salicina*). Salicaceae could therefore be a key component of networks through which *Xf* could spread. In France, reports from Corsica also indicated large populations of *Aphrophora* grp. *salicina* and *Cicadella viridis* in riparian forests (Chauvel et al., 2015). *Philaenus spumarius*, which is considered as the main *Xf* vector in Europe (Cornara et al., 2018), is particularly abundant in typical scrubland of the Mediterranean area. It is often found in strong association with *Cistus monspeliensis* (Chartois et al., 2023). Large populations of *P. spumarius* were also reported in grasslands (Cappellari et al., 2022) and alfalfa fields (Weaver & King, 1954). Alfalfa and red clover are indeed the only crops for which *P. spumarius* is considered a pest through direct feeding on plant sap (e.g. Parman & Curtis, 1982). Species in the genus *Neophilaenus* are known to specialize in Poaceae (Bodino, Cavalieri, Dongiovanni, Saladini, et al., 2020; Villa et al., 2020) and are very abundant in herbaceous habitats, especially in grasslands (Cappellari et al., 2022) but also in the herbaceous cover and headlands of groves (Bodino, Cavalieri, Dongiovanni, Plazio, et al., 2019; Mesmin et al., 2022). So far, almost nothing is known about the habitat-plant-vector networks in temperate wine-growing areas of Europe, which prevents any investigation on which levers we could act to prevent or mitigate transmission risk of *Xf* to grapevine.

In mainland France, *Xf* has been detected by official surveillance of plants in two wine-producing regions in southern France: Provence-Alpes-Côte d’Azur (PACA) and Occitanie (OCC). The subspecies *multiplex* is by far the most commonly detected. Only one focus of the subspecies *pauca* has been identified so far, near Menton (MASA, 2023) and is now eradicated (CROPSAV, 2024). *Xf* subsp. *fastiodiosa* has never been detected. *Xylella* (unidentified subspecies) has been fortuitously found once in vine bark in the Lussac Saint-Émilion wine-growing region (Martins et al., 2013). Although this result needs to be confirmed, it suggests that the bacterium might be present at very low prevalence levels in a third wine-producing region of France, Nouvelle-Aquitaine (NAQ), for several years.

In the present study, we characterized insect-habitat-plant interaction networks in the three above mentioned wine-growing regions of France. We addressed the following questions: 1/ How specialized are xylem feeders at the habitat and host plant levels? and 2/ What habitats are the biggest reservoirs of xylem feeders? Based on the literature, we expected that 1/ *P. spumarius* would be the most generalist at the habitat and host plant levels whereas other xylem feeders would be mostly specialists such as *Neophilaenus* spp. in meadows, or *Cicadella viridis* and *Aphrophora* spp. in riparian habitats (Casarin et al., 2021; Morente et al., 2018; Thompson et al., 2023); 2/ the biggest reservoirs would be mainly semi-natural herbaceous strata (meadow, alfalfa, riparian vegetation) (Casarin et al., 2021; Weaver & King, 1954).

In addition, we assessed whether or not *Xf* was present in the studied environment by relying on a sentinel insect approach. It relies on the idea that as vectors feed on various plants, their infectious status somewhat summarizes the infection status of the environment where they have been sampled (Cruaud et al., 2018; Farigoule et al., 2022; Yaseen et al., 2017). This strategy is an interesting addition to surveillance based on symptomatic plants, because it can reveal overlooked contamination (Cruaud et al., 2018; Farigoule et al., 2022). Indeed, while in certain species the proliferation of *Xf* can lead to plant death, in many species, the bacterium is commensal, and plants do not develop symptoms (Roper et al., 2019; Sicard et al., 2018). In this case, asymptomatic habitats can act as disease reservoir posing a threat to nearby susceptible crops. Given the results obtained by official surveillance we would expect that *Xf* could spread within insect-habitat-plant interaction networks in the Mediterranean region. However, if present, the prevalence of *Xf* in vector populations of the NAQ region would be low.

## Methods

### Study sites and sampling design

Our sampling strategy focused on a set of vineyards in which we obtained authorization to sample entomofauna from the growers in three wine-producing regions in France: Nouvelle-Aquitaine (NAQ), Occitanie (OCC), Provence-Alpes-Côte d’Azur (PACA) (Fig. 1).

**Figure 1.**
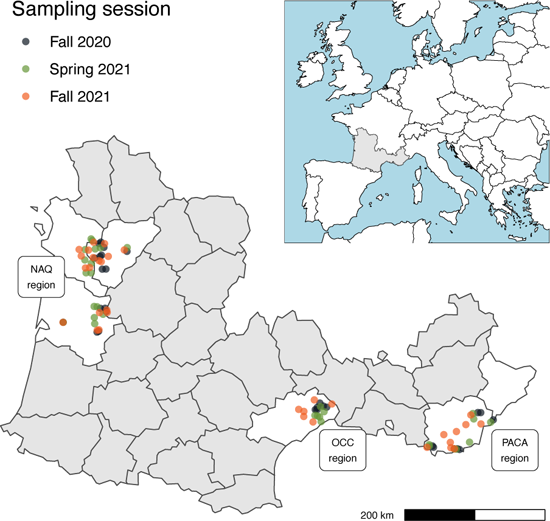
Map of the studied areas. The grey area on the inset shows the sampled area in France, and the zoomed-in main figure highlights in white the areas where buffers (colored dots) were placed.

We used a clustered sampling strategy (Table 1, Appendix 1) based on the dispersal ability of the main vectors of *Xf*, which is some hundreds of meters in their whole life, and exceptionally more than 1 km (Bodino, Cavalieri, Dongiovanni, Simonetto, et al., 2020; Casarin et al., 2022; Lago et al., 2021). We defined 1 km-radius buffers regularly distributed among the three studied areas (Fig. 1). These buffers included target vineyards and at least two types of land cover that could be reservoirs of *Xf* vectors according to the literature and our personal observations: olive groves, alfalfa fields, forests, meadows, riparian zones, and the borders of vineyards and olive groves. Inside each buffer, we sampled a median number of 7.5 sites. In each site, and when available, two subsites were sampled, the lower (below 1m, mostly composed of herbaceous plants) and the upper stratum (i.e. from 1 to 2 m above ground, mostly composed of woody plants). “Habitat” was here defined as a specific stratum in a specific land cover (e.g. “lower stratum of vineyards” or “upper stratum of vineyards”).

**Table 1.** Sample size and full description of the clustered sampling strategy used in the survey. Buffers comprise four to thirteen sites belonging to the land cover types analyzed. Each site includes two subsites (lower and upper strata) except when no vegetation exceeded 1m (no upper stratum). The sampling unit (i.e. on which the sampling protocol was applied) is the subsite. “x ⊃ y” stands for “x includes y” and denotes the clustered sampling strategy with several sites per buffer and two subsites (upper and lower stratum if available) per site. The two last columns show the number of xylem feeders sampled as well as the number used for *Xf* screening.

| Session | Region | Number of buffers | Number of sites | Number of subsites | Number of xylem feeders | Number of insects analysed for the presence of *Xf* |
| --- | --- | --- | --- | --- | --- | --- |
| Fall 2020 | NAQ | 15 | 111 | 169 | 1426 | 464 |
|  | OCC | 6 | 49 | 78 | 83 | - |
|  | PACA | 8 | 64 | 97 | 796 | 93 |
| Spring 2021 | NAQ | 16 | 108 | 161 | 1047 | - |
|  | OCC | 6 | 43 | 72 | 79 | - |
|  | PACA | 8 | 58 | 93 | 619 | - |
| Fall 2021 | NAQ | 18 | 127 | 191 | 3159 | 401 |
|  | OCC | 6 | 55 | 86 | 286 | - |
|  | PACA | 9 | 85 | 132 | 1277 | 59 |
| Total |  | 92 | 700 | 1079 | 8772 | 1017 |

Three sampling sessions were performed: two targeting the adults of xylem feeders, in the fall of 2020 (14^th^ Sept.-12^th^ Oct.) and 2021 (6^th^-28^th^ Oct.), and one targeting the nymphs in the spring of 2021 (12^th^ Apr.-9^th^ May). This timing is based on a preliminary phenological survey (unpublished data) conducted in the South and in the West of France and matches the peak densities of adults (in the autumn) and nymphs (in the spring). New buffers were selected for each sampling session to avoid pseudoreplication (Colegrave & Ruxton, 2018).

### Insect and plant sampling

For the adults, interaction networks were evaluated at the habitat level only. Adults were collected by passing a sweep net through the vegetation for a total of four minutes. The four minutes were divided into eight periods of 30-second of sweep netting followed by insect collection in the net using a mouth aspirator (the time needed for insect collection was not included in the 4’). At the end of the four minutes, insects in the mouth aspirator were killed using ethyl acetate, stored in 75° ethanol, and brought to the laboratory for identification under a binocular microscope using the Biedermann and Niedringhaus (2009) identification key. They were then stored in 96° ethanol at 4°C. Each subsite in each sampling site was sampled only once.

For the nymphs, interaction networks were evaluated at the habitat and the host plant levels (family and species). Indeed, with the exception of *C. viridis* that was therefore not sampled in spring, nymphs are concealed in a spittle lying on the vegetation. This makes the identification of host feeding plants possible. Nymphs were sampled using a quadrat procedure in each subsite.

To account for differences in vegetation density (dense or sparse) among lower and upper strata, we adapted the size of quadrats to sample a similar amount of vegetation in all strata. This was done by fixing the lower strata quadrats to 1 # × 0.25 # (Bodino, Cavalieri, Dongiovanni, Plazio, et al., 2019) and choosing upper quadrat sizes so that the sampling effort in terms of time spent was the same. We found that a good compromise was to increase the size of upper strata quadrats to 2 # × 0.5 #, and the upper strata of vineyards to 5 # × 0.5 # as vegetation cover was usually very low in April-May (Appendix 2). Three quadrats were positioned randomly in each subsite. Datasets were summed at the subsite level for data analysis.

For each spittle detected in the quadrat, the nymph(s) was/were collected with a paintbrush and placed in vials filled with 75° ethanol. They were later identified in the lab following Stöckmann et al. (2013) and Mesmin et al. (2022). *Neophilaenus campestris* and *N. lineatus* were found morphologically indistinguishable at the nymph stage. Therefore, nymphs of this genus were pooled in the *Neophilaenus* sp. group. Similarly, no morphological features are currently reliable to distinguish nymphs of *A. salicina* from those of *A. pectoralis* (Streito et al., 2019). All nymphs were thus labelled *Aphrophora* grp. *salicina*. The plant on which the spittle was found was collected and placed in a ziploc bag together with the insect tube, and identified in the lab following Tison & de Foucault (2014), Tison et al. (2014) and Eggenberg & Möhl (2020). The nomenclature follows the GBIF Backbone Taxonomy (GBIF Secretariat, 2023), checked using the R package ‘traitdataform’ (Schneider, 2018) to detect synonyms.

### *Xf* prevalence in insects

Screening of *Xf* was performed in 1017 adults sampled in the two most contrasted regions regarding *Xf* infection status: PACA, where *Xf* has been found as early as 2015, and NAQ where *Xf* has never been detected by official surveillance so far (Table S1.1, Appendix 1). For financial reasons insects were analysed only when they were collected in vineyards and vineyard borders belonging to the network of growers of our funder or in buffers belonging to INRAE. Areas in which *Xf* screening was performed are shown in Appendix 1 (Fig. S1.4 & S1.5). We covered most of the area around Cognac (NAQ), included samples from semi-natural habitats around Bordeaux (NAQ) and a buffer 25 km away from official detections in PACA (Martinetti & Soubeyrand, 2018). For molecular analyses, screened populations were defined as groups of 30 insects from the same species, sampled in the same habitat of the same buffer. Adults in all populations of xylem feeders sampled in 2020 were screened for *Xf*. All specimens tested negative for *Xf.* Therefore, in 2021, we chose to screen only populations of *P. spumarius* and *N. campestris* which are known vectors of *Xf* (Cavalieri et al., 2019), as well as *N. lineatus*, whose vector status has not been assessed yet but which is closely related to *N. campestris*.

*Xf* screening was performed using the protocol described in detail in Farigoule et al. (2022). Briefly, DNA was extracted from individual specimens following Cruaud et al. (2018) to minimize the impact of PCR inhibitors. *LeuA*, one of the housekeeping genes of *Xf* (Yuan et al., 2010) was amplified using the two-step PCR approach described in Farigoule et al. (2022). Four replicates were performed for each specimen and a unique combination of indexes was used for tagging all PCR products obtained from the same specimen. Sequencing was performed on a MiSeq system (2 × 250)*). The complete pipeline to analyze raw sequence data is available from Farigoule et al. (2022) [https://github.com/acruaud/prevalenceXfinsectclimate_2022].

### Ecological data analysis

Data analyses were performed using the programming language “R” (R core team, 2022). Networks and associated “species-level” (Dormann, 2011) metrics were computed using the package ‘bipartite’ (Dormann et al., 2008).

#### Network computation to assess insect specialization

Comparing species specialization levels is notoriously challenging (Dorado et al., 2011), especially when small networks are analysed (Fründ et al., 2016). Three precautions were thus taken, based on the literature, to avoid misinterpretation of specialization levels. Firstly, we computed networks at the grain of the region (NAQ, OCC, PACA) to have at least 20 individuals per insect species in most cases (Fründ et al., 2016), and removed singletons and doubletons (i.e. only one or two individuals of a taxon in the network, Lami et al., 2021). Secondly, we used different specialization metrics. We computed the Resource Range (RR) and the Paired Difference Index (PDI), which are known to efficiently segregate specialists from generalists including in low connectance networks (Poisot et al., 2012). We also calculated the *d’* (Blüthgen et al., 2006) and the Species Specificity Index (SSI, Julliard et al., 2006), that are less sensitive to differences in abundance among consumers (Fründ et al., 2016). Thirdly, in addition to the raw specialization metrics, we used the z-transformed metrics computed relatively to the distribution of the metric in 1000 random networks drawn from fixed marginal totals (e.g. Lami et al., 2021). This approach helps to disentangle specialization and sample sizes (Dormann et al., 2017). Because *d’* already integrates a confrontation to a null model (Dormann et al., 2017), only the RR, PDI and SSI were analyzed using z-scores. All metrics increase with specialization, either in absolute value (raw scores), or relative to the probable distribution of values given the sampling design (z-scores).

Three types of interaction networks were computed, that varied in the definition of the resource level, because we expected that insect specialization could occur at various levels depending on the species: the habitat, the plant family or the plant species. In habitat (resp. plant family or species) networks, the strength of the interaction between an insect species and a habitat (resp. a plant family or species) was the total number of individuals of that species found in this habitat (resp. on this plant family or species) for a given region and a given sampling session. Insect-plant family or plant species networks based on nymph data were computed only in the spring of 2021. For all specialization metrics involving a null model in insect-habitat networks, marginal totals were set proportional to the sampling intensity of habitats in the corresponding networks (see Appendix 3 for details) using the function ‘mgen’ of the package ‘bipartite’ (Dormann et al., 2008). For the insect-plant family and species networks, marginal totals were only realized interactions (function ‘rd2table’).

#### Statistical analyses

As specialization metrics give different information (e.g. RR only accounts for presence/absence, others use quantitative data) and have different sensibility to network properties (e.g. connectance, relative abundances), our main analysis gathered all seven specialization metrics. We performed three GLMMs (Bolker et al., 2009) – one for insect-habitat, one for insect-plant families and one for insect-plant species networks – with the following formula:

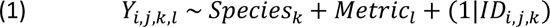

where *Y_i,j,k,l_* is the insect specialization metric *l* computed on region *i* on session *j* for species *k*. The random effect set on the interaction between network ID and species was added to avoid inflated statistical power due to pseudoreplicated data (i.e. multiple assessments of specialization for the same species on the same network). For all GLMMs performed in this study, we followed the modeling procedure detailed in Appendix 4. Briefly, models were fitted using the simplest distribution and link yielding a correct model regarding basic modeling assumptions tested with the package ‘DHARMa’ (Hartig, 2020). Once fitted, they were analyzed using an analysis of deviance tables (package ‘car’, Fox & Weisberg, 2019). Insignificant fixed terms were removed and the modeling procedure was applied again until all fixed terms were significant. At the end of each modeling procedure, pairwise comparisons among the modalities of significant fixed terms were performed using the packages ‘emmeans’ (Lenth, 2020) and ‘performance’ (Lüdecke et al., 2021) and marginal R^2^ (i.e. part of variance explained by fixed effects) were computed (Nakagawa & Schielzeth, 2013).

Additionally to network metrics, raw abundances of *P. spumarius*, *Neophilaenus* sp. and *C. viridis* were analyzed with GLMMs to assess their habitat preferences using the following formula:

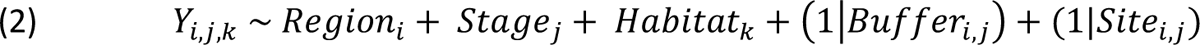

where *Y_i,j,k_* is the abundance of an insect species metric on region *i* on session *j* for habitat *k*. Random effects were added to consider that metrics computed on the same buffer and the same site (upper and lower strata) were not independent.

To address our second hypothesis (the biggest reservoirs would be mainly semi-natural herbaceous strata), we modeled xylem feeder abundance (sum of all species) and diversity (effective number of species, i.e. exponential of Shannon index; Jost, 2006; set to NA for null abundances) in each subsite using GLMMs, with formula (2).

## Results

Overall, we sampled 8772 xylem feeders belonging to, in decreasing abundance: *P. spumarius*, *N. campestris*, *N. lineatus*, *C. viridis*, *A.* grp. *salicina*, *L. coleoptrata* and *A. alni* (Fig. 2, Appendix 1). Except for 1 individual collected in the PACA region, *N. lineatus* was only encountered in the NAQ region (in similar proportion with *N. campestris*; 669 *N. campestris* / 680 *N. lineatus*, Fig. 2). *Cicadella viridis* and *L. coleoptrata* were absent from samplings conducted in the OCC region. At the nymph stage, *L. coleoptrata* was only collected in the NAQ region (Fig. 2). For the statistics described below, all details on χ^2^, degrees of freedom and p-values are available in the tables of Appendix 4.

**Figure 2.**
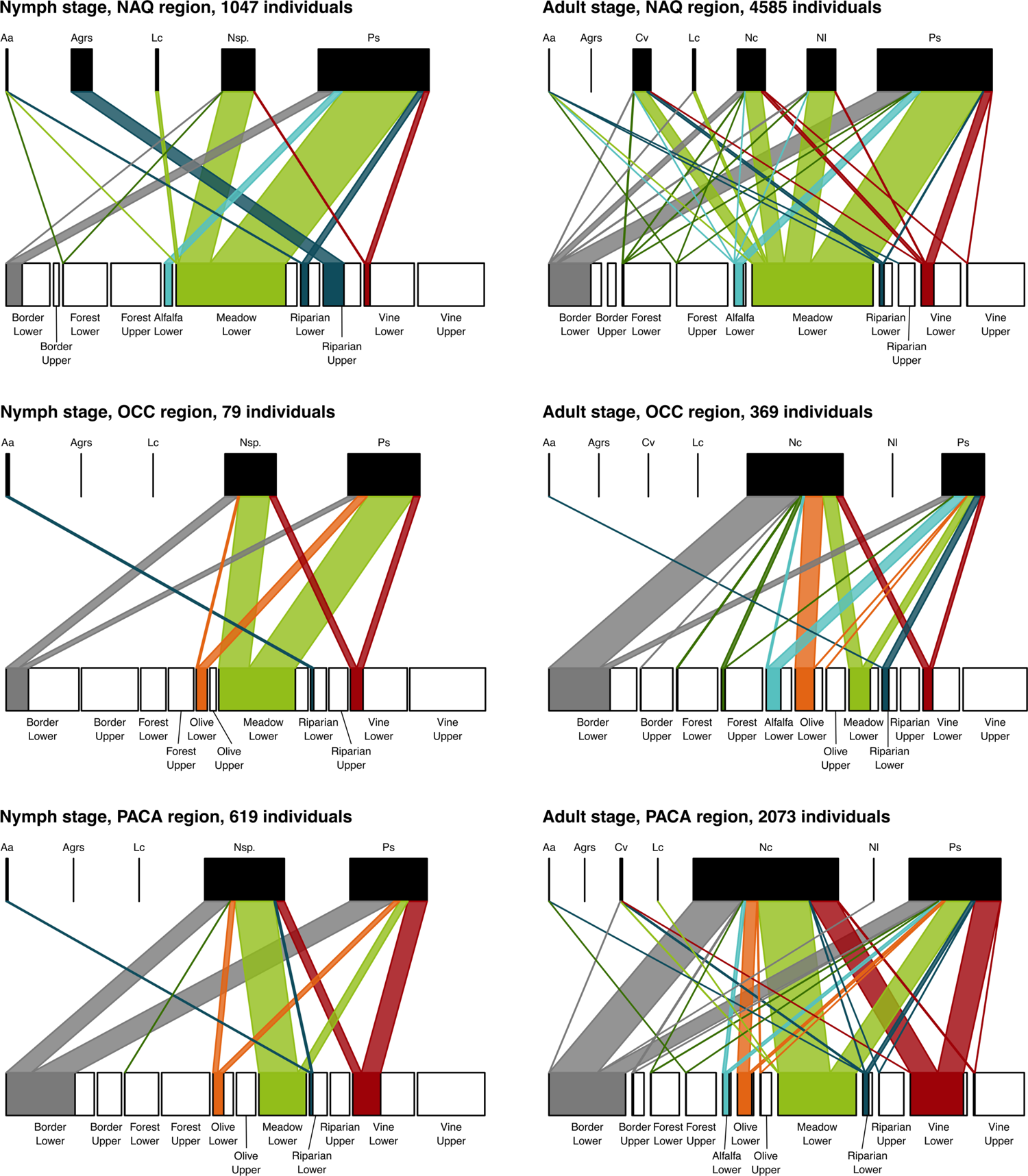
Insect-habitat networks for nymph (left) and adult (right) stages in the NAQ, OCC and PACA regions. At the resource (bottom) level of each network, the width of each habitat box (filled plus empty portion) is proportional to its sample size (i.e. number of subsites among all subsites surveyed) in the network displayed. For each habitat, the empty portion of the box shows the proportion of subsites on which no insect was found. Insect species are abbreviated as follows Aa: *Aphrophora alni*, Agrs: *Aphrophora* grp. *salicina*, Cv: *Cicadella viridis*, Lc: *Lepyronia coleoptrata*, Nc: *Neophilaenus campestris*, Nl: *Neophilaenus lineatus*, Nsp.: *Neophilaenus* sp. And Ps: *Philaenus spumarius*. See Appendix 3 for details on the computation involved.

### Xylem feeders specialization on habitats and host plants

At the level of habitats, xylem feeders had similar specialization degrees (Fig. 2 & 3A): most species were ubiquitous except for *A.* grp. *salicina* that was restricted to riparian habitats (Fig. 2). Among all specialization metrics, this latter species was more specialized than *N. campestris*. Additionally, *N. lineatus* was mostly restricted to meadows, and globally more specialized than its congeneric *N. campestris*, and also more specialized than *P. spumarius*. At the habitat level, *Neophilaenus* sp. nymphs were distributed similarly to adults of *N. campestris* or *N. lineatus* (Fig. 3).

**Figure 3.**
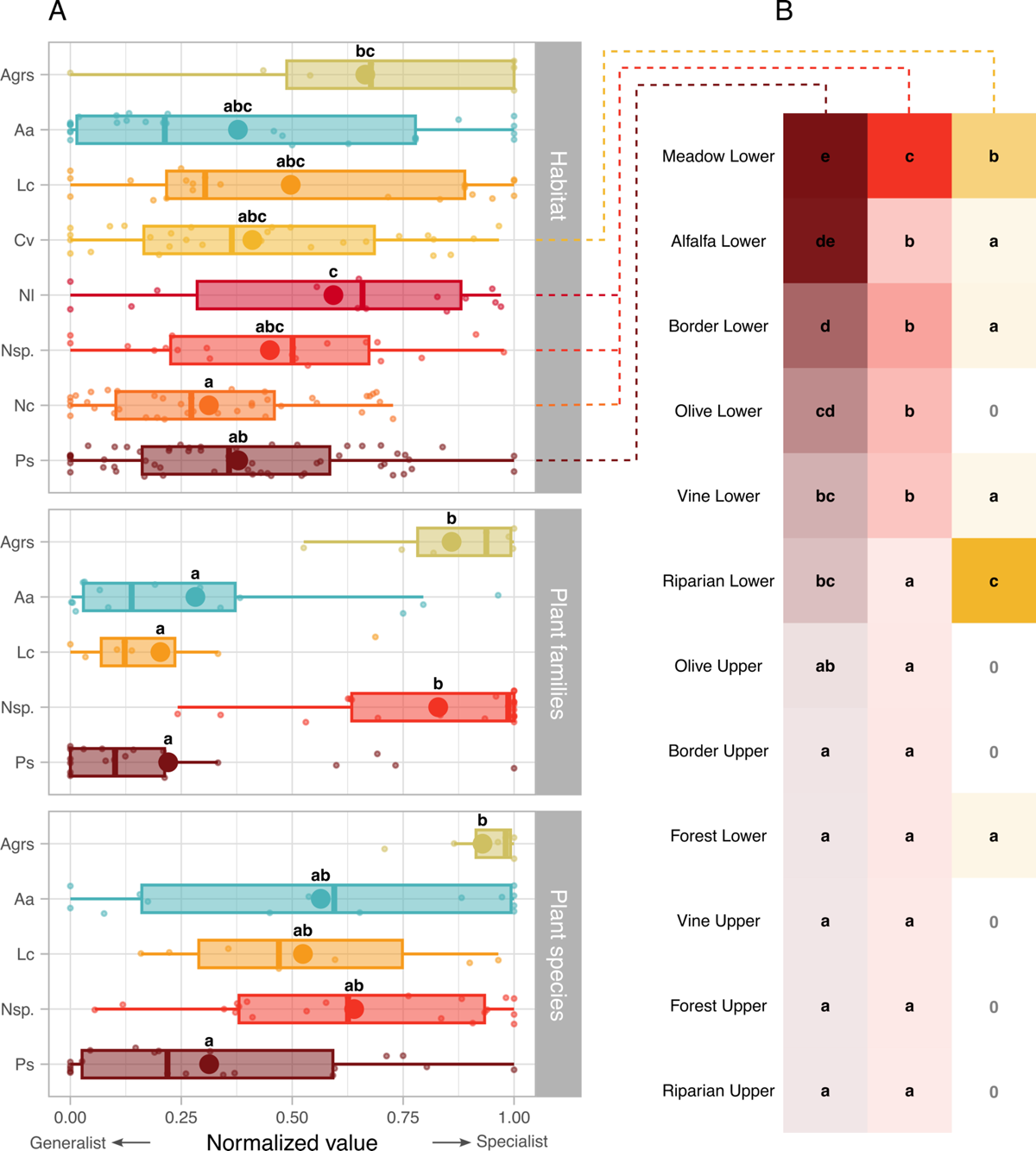
A – Distribution of specialization values for insect-habitat (top panel), insect-plant families (middle panel) and insect-plant species (bottom panel) networks. Small points in the background depict the data: one point is one specialization metric for a given species in each network (e.g. one point in the top panel depicts the resource range of *P. spumarius* in the insect-habitat network assessed in the NAQ region in fall 2020). The big points depict means. The boxplots depict quartiles and extend up to 1.5 times the inter-quartile range. The letters report statistics performed on the distributions: in each panel independently, species that share a letter have specialization degrees that are not significantly different. Note that in the top panel, habitat preference for *Cicadella viridis*, *Neophilaenus lineatus* and *Neophilaenus campestris* were only assessed at the adult stage and that of *Neophilaenus* sp. was only assessed at the nymphal stage. **B –** Habitat preferences of the three most abundant xylem feeders. Color ramps are scaled independently for the three species and depict log-transformed estimated marginal means of GLMMs. For each species independently, habitats sharing a letter do not differ significantly. Zeros indicate habitats where the species was never collected. Insect species are abbreviated as follows Aa: *Aphrophora alni*, Agrs: *Aphrophora* grp. *Salicina*, Cv: *Cicadella viridis*, Lc: *Lepyronia coleoptrata*, Nc: *Neophilaenus campestris*, Nl: *Neophilaenus lineatus*, Nsp.: *Neophilaenus* sp. And Ps: *Philaenus spumarius*. See Appendix 4 and Appendix 5 for more details.

Although ubiquitous – i.e. able to thrive in various habitats – the generalist xylem feeders had habitat preferences, and this was particularly clear for *P. spumarius*, *Neophilaenus* sp. and *C. viridis* (Fig. 3B, Table S4.2). *Philaenus spumarius* was found in higher abundances on lower strata of meadows and alfalfa fields (Fig. 3B & S6.1). It was found gradually less on the lower strata of field borders, olive groves, grapevines and riparian vegetation (Fig. S6.1) and only rarely on the upper strata of field borders (i.e. hedges), grapevines, riparian zones, forests, olive groves, as well as on the lower stratum of forests. It was globally less abundant in the OCC than in the PACA or NAQ regions (Fig. S6.1). *Neophilaenus* species (including here *N. campestris* and *N. lineatus* at both nymph and adult stages) were most abundant in meadows, and then, to a lesser extent, on the lower strata of the borders of vineyards/olive groves, of olive groves, of vineyards and of alfalfa (Fig. 3B & S6.1). Similarly to *P. spumarius*, *Neophilaenus* species were scarce on the upper strata of borders of vineyards/olive groves (i.e. hedges), of vineyards, of riparian zones, of forests, of olive groves, as well as on the lower stratum of forests and riparian areas. *Neophilaenus* species were more abundant in the PACA region than in the OCC or NAQ regions (Fig. S6.1). Finally, *C. viridis* was found mostly on the lower stratum of riparian zones, then on meadows, and was rare on lower strata of alfalfa, of borders of vineyards/olive groves, vineyards and forests (Fig. 3B). It was not found in the other habitats. *Cicadella viridis* was on average 6.5 times more abundant in the NAQ than in the PACA region (Table S4.2) and it was not found in the OCC region.

*Philaenus spumarius* was found on 18 plant families, followed by *A. alni* (8), *L. coleoptrata* (6), *Neophilaenus* sp. (4), and *A.* grp. *salicina* (2, Table S8.1). Two groups of species could be differentiated: *P. spumarius*, *L. coleoptrata* and *A. alni* were the most generalists, while *Neophilaenus* sp. And *A.* grp. *salicina* were the most specialists (Fig. 3A & 4). *Neophilaenus* sp. was almost restricted to Poaceae and *A.* grp. *salicina* was mostly restricted to Salicaceae (found only once on *Fraxinus excelsior*, an Oleaceae, Fig. 4).

**Figure 4.**
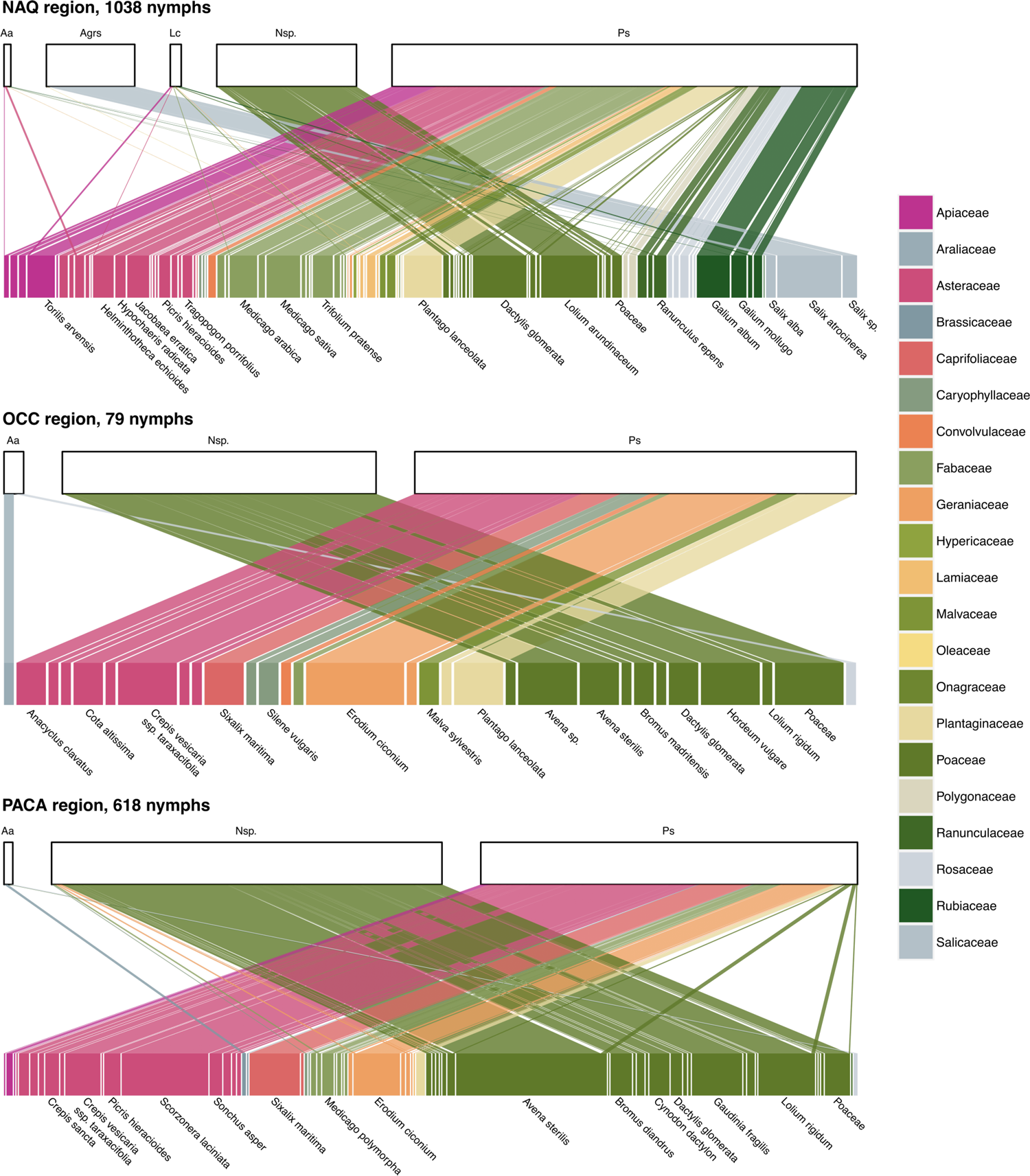
Insect-plant networks obtained in the spring 2021 for nymphs of xylem feeders. Only the names of the most abundant plant species are displayed here for the sake of readability. See Appendix 7 and Appendix 8 for full details on these three networks. Note that differences with Fig. 2 in the number of nymphs are due to 20 nymphs found on plants that could not be identified even at the family level. Insect species are abbreviated as follows Aa: *Aphrophora alni*, Agrs: *Aphrophora* grp. *salicina*, Cv: *Cicadella viridis*, Lc: *Lepyronia coleoptrata*, Nc: *Neophilaenus campestris*, Nl: *Neophilaenus lineatus*, Nsp.: *Neophilaenus* sp. and Ps: *Philaenus spumarius*.

At the plant species level, it was again *P. spumarius* that was found on the highest number of plants (87 species), followed by *Neophilaenus* sp. (31), *A. alni* (10), *L. coleoptrata* (7), and *A.* grp. *salicina* (3, Table S8.1). Consistently, all specialization metrics combined, *P. spumarius* was more generalist than *A.* grp. *salicina*, and the three other taxa – *Neophilaenus* sp., *A. alni* and *L. coleoptrata* – had intermediate specialization levels (Fig. 3A).

### Habitat exploitation by xylem feeders

The total abundance of xylem feeders per subsite was significantly correlated with the habitat type (Fig. 5, Table S4.3), meaning that habitats were not equally exploited by insects (Fig. 2). The least exploited habitats, for both xylem feeder abundance and diversity, were the upper strata of forests, vines, riparian areas, borders (i.e. hedges), olive groves, as well as the lower stratum of forests (Fig. 5, and see empty boxes in Fig. 2). Due to lack of variation of insect diversity in these habitats (always only one xylem feeder species found), xylem feeder diversity could not be statistically compared to other habitats. Xylem feeder abundance was intermediate in lower strata of riparian areas, vines, olive groves, borders and alfalfa. Finally, the habitat where xylem feeders were the most abundant was the lower stratum of meadows (Fig. 5, see also the proportion of full boxes in Fig. 2). In terms of diversity, although a significant effect of habitat was detected (Table S4.3), most habitats shared similar diversities (Fig. 5). Overall, xylem feeder abundance per subsite was higher in the NAQ and PACA regions than in the OCC region, and diversity was higher in the PACA region than in the two others (Fig. 5, Table S4.3).

**Figure 5.**
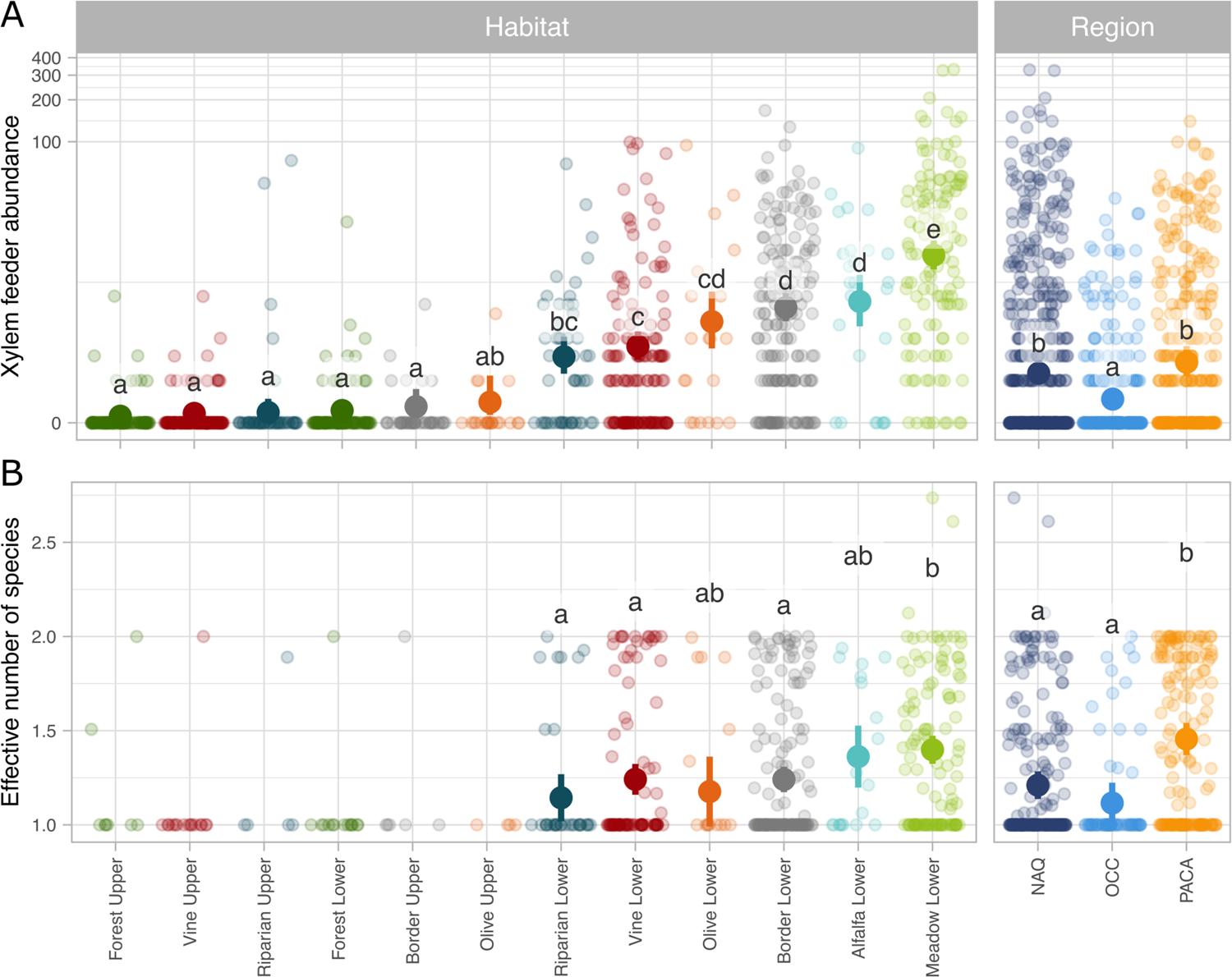
Results of the GLMM performed on the abundance of xylem feeders (A) and xylem feeder species diversity (B, effective number of species, exponential of Shannon index) for each habitat and each region. Letters depict the significance of the effect of habitats and regions. For each panel taken independently, modalities sharing a letter do not differ significantly.

### *Xf* prevalence in xylem feeders

Overall 714 *P. spumarius*, 156 *N. campestris*, 96 *N. lineatus*, 31 *C. viridis*, 17 *L. coleoptrata* and 3 *A. alni* sampled in the NAQ (n=865) or PACA (n=152) regions were screened for *Xf* (Table S1.1). All insects tested negative.

## Discussion

In the present study, we identified insect-habitat-plant interaction networks in vineyards and their surrounding landscapes. We also assessed the prevalence of *Xf* in the environment based on a sentinel insect approach.

### Most xylem feeders are generalists

Our first hypothesis was that *P. spumarius* would be extremely generalist and that other xylem feeders would be mostly specialists at the habitat and host plant levels. Using interaction networks that have previously proven effective in assessing habitat specialization in insects (Lami et al., 2021; Marini et al., 2019), we found that this hypothesis was only partially supported. At the habitat level, the degree of specialization of *P. spumarius* was very similar to that of *A. alni*, *C. viridis*, *L. coleoptrata* and *N. campestris*. This suggests that the widely reported ubiquity of *P. spumarius* (Cornara et al., 2018, 2021) is shared with most spittlebugs and sharpshooters in the studied landscapes of southern France. Despite this ubiquity, xylem feeders did not distribute evenly among all studied habitats. As expected from the literature, *Neophilaenus* spp. preferred meadows over other habitats. *Neophilaenus lineatus* was indeed almost restricted to meadows. *Philaenus spumarius* also showed a marked preference for meadows. It is noteworthy that we confirm for the first time in Europe its association with alfalfa. This association was only documented in Ohio and Michigan so far (Weaver & King, 1954; Wiegert, 1964). Finally, *A.* grp. *salicina* was almost restricted to riparian habitats.

The plant family level (assessed at the nymph stage only) was more relevant to identify and differentiate specialization level among spittlebugs. *Philaenus spumarius*, *A. alni* and *L. coleoptrata* were generalists. On the opposite, *Neophilaenus* sp. and *A.* grp. *salicina* were specialists, each thriving on almost only one plant family, confirming the biological informations available for these species (Casarin et al., 2021; Morente et al., 2018).

Finally, although *P. spumarius* had the highest generality score at the plant species level, it was not significantly more generalist than *A. alni*, *L. coleoptrata* or *Neophilaenus* sp. The 18 plant families on which it was found in the present study were also reported by Thompson et al. (2023). However, we observed *P. spumarius* on 6 plant genera not yet reported in the literature – namely *Anacyclus*, *Carthamus*, *Cota*, *Petrorhagia*, *Thrincia* and *Thyrimnus* – and 30 new plant species records (see Table S10.1 & S10.2 in Appendix 10). When we group all association records for *P. spumarius* and following the GBIF Backbone Taxonomy (GBIF Secretariat, 2023), we reach a total of 670 plant genera and 1279 plant species (Appendix 10).

Interestingly, species in *Neophilaenus* were extremely specialized at the plant family level: they were found only occasionally on dicots. They were relatively generalists at the plant species level, being able to feed on a wide variety of graminaceous species. As reported in previous studies, *Neophilaenus* spp. were frequently found on *Dactylis glomerata* and *Avena sterilis* (Antonatos et al., 2021; Dongiovanni, Cavalieri, et al., 2019; Morente et al., 2018) but we also highlight high frequencies on *Lolium arundinaceum*, *Hordeum vulgare*, *Lolium rigidum* or *Gaudinia fragilis*. Our study provides a list of 31 species of Poaceae on which nymphs of *Neophilaenus* spp. can feed, 26 of which are new when we compare our list with the 21 host species reported by Antonatos et al. (2021).

Finally, we found several nymphs of *A. alni* on *Pteridium aquilinum* (opportunistic extra samplings, Appendix 9, Fig. S9.4), one of the most ubiquitous species in the world, able to develop below forest canopies and in open areas (Dumas et al., 2022). More work is needed to determine if *A. alni* only occasionally feeds on that fern, or if bracken covers represent a significant reservoir of this xylem feeder.

To summarize the answer to our first research question, we expected *P. spumarius* to stand out clearly as the most generalist (Cornara et al., 2018; Thompson et al., 2023) at all studied levels, however this was generally not the case. We hypothesize that this expectation was driven by the frequent confusion between abundance and generalism (Dorado et al., 2011; Fort et al., 2016) and the relatively poor knowledge we have on other xylem feeders. Here we tried to disentangle dietary niche from abundance by using several tools to better describe our networks (Dormann et al., 2017; Fründ et al., 2016). Our results indicate that *P. spumarius* is abundant and generalist. *Aphrophora alni*, *L. coleoptrata* – and to a lesser extent *C. viridis* – are also generalists, but not abundant. The only “true” specialist in our study, *A.* grp. *salicina*, is generally not abundant. Finally, *Neophilaenus* has a hybrid status, being extremely specialist at the plant family level, but generalist at the habitat and plant species levels. These results are in line with the general pattern described by Fort et al. (2016) in plant-pollinator and plant-seed dispersers networks: abundance implies generalism (i.e. specialists are rarely abundant), whereas the reverse is false (i.e. generalists can be rare).

### Herbaceous semi-natural habitats are the main sources of xylem feeders

Our second hypothesis was supported by the results. The habitats where xylem feeders are the most abundant are indeed mostly semi-natural habitats. Exploring a large diversity of habitats of the agricultural mosaic, we found two groups of habitats. The first group gathers habitats that are almost free of xylem feeders. It includes the upper strata of forests, of riparian vegetation, of vineyards, of olive groves, hedges, and the lower stratum of forests. This is a reassuring result for forests because some tree species, *Quercus* species in particular, are vulnerable to *Xf* (Desprez-Loustau et al., 2021). The second group comprises habitats substantially exposed to xylem feeders, with an increasing gradient from the lower stratum of riparian vegetation up to meadows (very exposed).

For the two crops of significant economic value included in our sampling scheme, grapevine and olive, xylem feeders were found very rarely on the foliage. Two habitats that host large populations of xylem feeders are nevertheless near grapevine or olive foliage: the lower strata of vineyard/olive grove and their herbaceous borders, especially in PACA vineyards. This is in line with several studies showing that most potential vectors of *Xf* reach high abundances in herbaceous habitats – see e.g. Rodrigues et al. (2023) for a recent study in Portuguese vineyards – especially at the nymphal stage (Dongiovanni, Cavalieri, et al., 2019). Xylem feeders have been reported to migrate to crop foliage when the herbaceous vegetation gets too dry in the context of Italian olive groves (Bodino et al., 2021; Cornara et al., 2017), but also in Californian vineyards (Beal et al., 2021). Besides inter-rows and borders, meadows and alfalfa fields, many of which were within the dispersal ability of *Xf* vectors to vineyards studied here, hosted the largest xylem feeder communities. Similarly, in almond orchards and associated landscapes in California, xylem feeders are extremely abundant in open herbaceous habitats that contain many host species for *Xf* (Krugner et al., 2012; Shapland et al., 2006) and insect transfer from these habitats to almond nurseries were suggested as a potential infection pathway to almond trees (Krugner et al., 2012). In our context, xylem feeders are also likely to move from alfalfa or meadows to vineyards or olive groves at the hay harvest. Indeed, migration to neighboring cereal crops following hay cutting have been reported for *P. spumarius* in Ohio (Weaver & King, 1954) or migration to wheat fields has been observed in the NAQ region (XM, personal observation, 2023). In summary, across these different contexts (Italian olive groves, Californian vineyards, Californian almond groves), there is a consistent pattern where xylem feeder communities establish themselves on herbaceous covers and move into foliage when the herbaceous habitat conditions deteriorate (i.e. drying or harvesting). Our results suggest that this pattern could lead to *Xf* spread to vineyards (and olive groves) of southern mainland France as well. However, what we currently lack is year-round monitoring of xylem feeders to detect potential migrations to crop foliage after summer droughts or harvesting operations.

In this context, we identify few options to lower the transmission risk. The best option still remains to increase surveillance to avoid introduction and spread of the subspecies fastidiosa in France. There is little room to lower risk through sowing of inter-rows and of headlands, as many potential vectors of *Xf* are highly generalist. Still we could advise favoring dicotyledons to limit *Neophilaenus* spp. populations. *Avena* spp. that were seen in some vineyards in association with *Vicia faba* do not seem advisable as they host considerable populations of *Neophilaenus* spp. Soil tillage to reduce vegetation cover could be the most efficient lever (Dongiovanni, Fumarola, et al., 2019) but this practice favors soil erosion especially in Mediterranean areas (Novara et al., 2011). At the landscape scale, a stepwise harvesting of meadows and alfalfa plots might be interesting to avoid massive migrations of xylem feeders.

Interestingly, two contrasted dissemination pathways reported i/ in Californian vineyards and *Citrus* groves or ii/ in Corsican groves are very unlikely in the context studied here. The former involves *G. atropunctata* which is considered responsible for most of the primary transmission from riparian habitats to grapevines (Hopkins & Purcell, 2002; Purcell, 2013) and then *Homalodisca vitripennis* for most of the secondary transmission as it reaches high abundances within vineyards and *Citrus* groves (Almeida & Nunney, 2015). In our context, *A.* grp. *salicina* or *C. viridis* are unlikely to play the role of *G. atropunctata* because *A.* grp. *salicina* is restricted to Salicaceae in riparian habitats and because *C. viridis* appears almost unable to transmit *Xf* (Bodino, Cavalieri, Dongiovanni, Altamura, et al., 2019; Bodino et al., 2022). Furthermore, in our context, xylem feeders do not reach high densities within the crops, unlike the situation observed with *H. vitripennis.* The Corsican example involves *P. spumarius* that shows a very strong preference for *C. monspeliensis* in olive and *Citrus* grove landscapes (Mesmin et al., 2022). A potential dissemination pathway was identified at the interface between *C. monspeliensis* scrubland and groves. Here, woody habitats were not identified as significant sources of xylem feeders. Apart from riparian habitats mentioned earlier, we did not find any xylem feeder likely to spread Xf at the forest-agroecosystem interface or within forests.

### *Xf* was not detected in the insects sampled

*Xf* was not found in the xylem feeders analyzed in this study. The absence of *Xf* in our samples from the PACA region is surprising, because *Xf* foci were officially detected ca. 25 km from some of our sampling sites in 2015 (DRAAF, 2017), and *Xf* dissemination by insects in Apulia is reported at about 10 km per year (Barjol et al., 2018), though in different landscape conditions. This result suggests that the spread of *Xf* is low in this area, possibly due to the eradication of infected plants, possibly also because infected plants were found in urban environments that have few large herbaceous areas favorable to xylem feeders. The bacterium was not present in sampled agricultural areas and the hazard of *Xf* transmission to plants is therefore very limited to date, even though xylem feeders are present. The absence of *Xf* in the NAQ region is less surprising because *Xylella* had only been found once in vine bark, a detection that has not been confirmed ever since. This result suggests that one of the main wine-growing areas in France is still uninfected by the *Xf*. The closest *Xf* focus to date is more than 100 km away (near Toulouse (DRAAF Occitanie, 2022), ssp. *multiplex*). However the recent finding of *Xf fastidiosa*, the subspecies that is harmful to vineyards, progressing quickly in southern Italy (EPPO, 2024; Teatro Naturale, 2024) indicates that it may become a threat for the European mainland. In both areas as well as in in the French Occitanie region, we recommend enhancing surveillance and supplementing it with the sentinel insect approach as a powerful tool for the early detection of infection foci, especially when plants do not show clear symptoms (Yaseen et al., 2017).

Available species distribution models indicate that the current climate conditions are suitable for *Xf fastidiosa* in different territories of Europe, including the French regions surveyed in the present study (Godefroid et al., 2019). Climate suitability is also expected to increase in the coming decades as climate change will lead to milder winters, hence lowering the cold curing effect (Godefroid et al., 2022; Purcell, 1980). These trends indicate that the risk of *Xf fastidiosa* establishment and/or expansion in Europe is and will remain high. Less is known about the impact of climate change on the range of the insect vectors, but Godefroid et al (2021) showed that *P. spumarius*, the main vector of *Xf,* is expected to remain present on most of the continent. According to these results, both *Xf fastidiosa* and its main insect vector will remain a serious risk in most of the wine-producing regions in Europe. Importantly, surveillance should also target quarantine, highly competent vectors of Xf especially *H. vitripennis* for which the climate conditions of the European continent are and will remain suitable (Rossi & Rasplus, 2022) or *G. atropunctata*.

Finally, it seems crucial to develop and test prophylactic and control strategies in the near term, particularly since *Xf fastidiosa* is currently absent. This will allow us to experiment with several strategies on xylem feeders communities without any epidemiological repercussions.

## Acknowledgements

We thank Jérémy Minguez, Malika Rouzes, Étienne Ramadier, Anna Dessaudes, Jennifer Dudit, Alice Bedani and Julien Pradel for their punctual help in the field work. We thank the UMR SAVE, especially Pauline Tolle and Adrien Rusch for giving us access to the BACCHUS winegrower network.

## Data, scripts, code, and supplementary information availability

Data and scripts are available online: https://zenodo.org/records/11181990 (Mesmin et al., 2024).

## Author contributions

XM, MC, PF, JYR, AC and JPR conceptualized the experimental design. AC and JPR supervised the project. XM coordinated the field work. XM, MC, PF, CB, JCS, JMT, EP, ML, YM, OB and IvH collected field data. XM and MC performed insect identification. XM and GF performed plant identification. XM and MC performed the statistical analyses. XM and MC led the writing of the manuscript. AC, JPR, CB and IvH contributed substantially to revisions. All authors provided final manuscript review and revision.

## Appendices

### Appendix 1: Geographical distribution of the sample size

**Figure S1.1.**
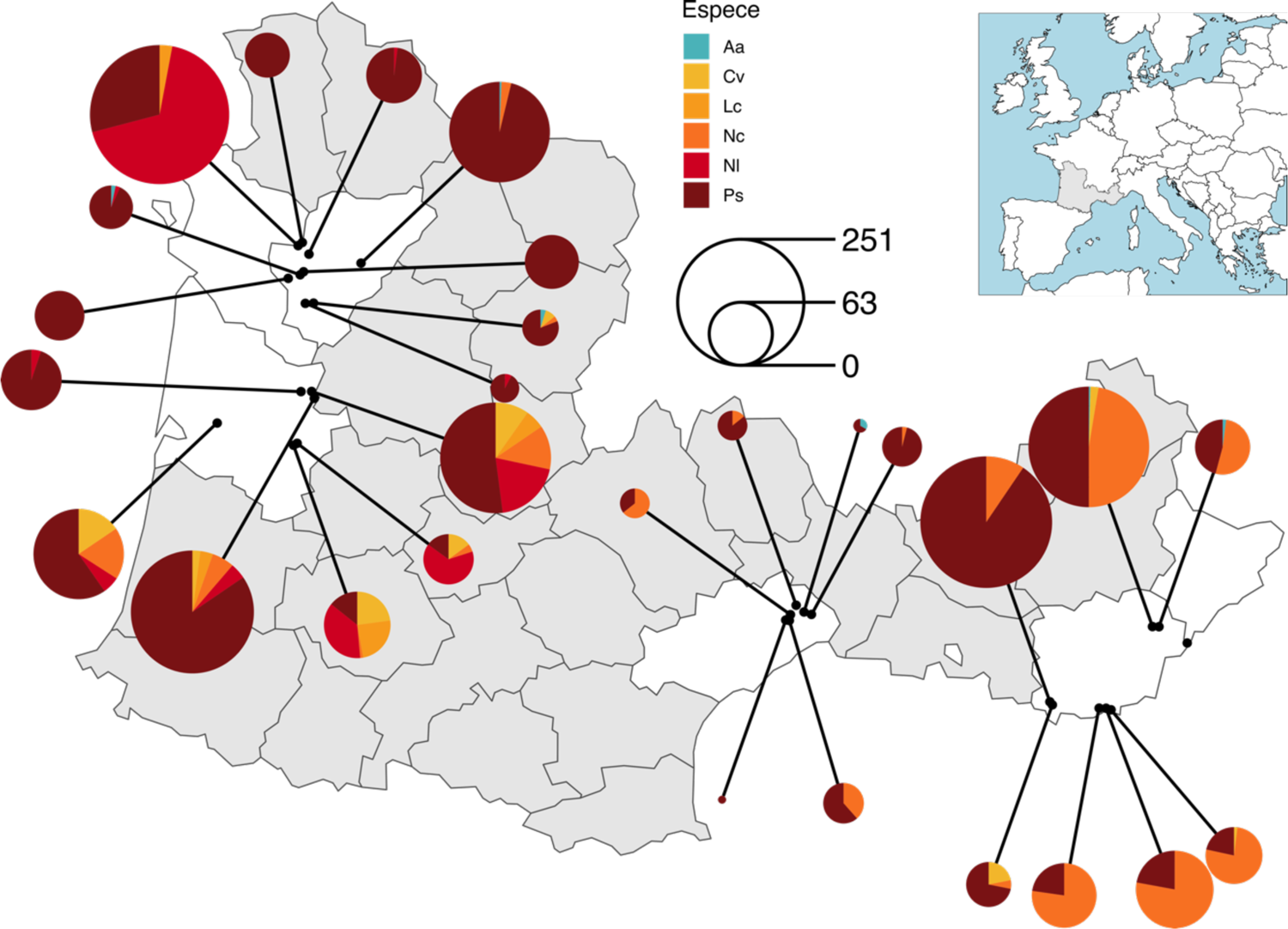
Distribution of insects sampled per buffer in the fall 2020. Insect species are abbreviated as follows Aa: *Aphrophora alni*, Agrs: *Aphrophora* grp. *salicina*, Cv: *Cicadella viridis*, Lc: *Lepyronia coleoptrata*, Nc: *Neophilaenus campestris*, Nl: *Neophilaenus lineatus*, Nsp.: *Neophilaenus* sp. and Ps: *Philaenus spumarius*.

**Figure S1.2.**
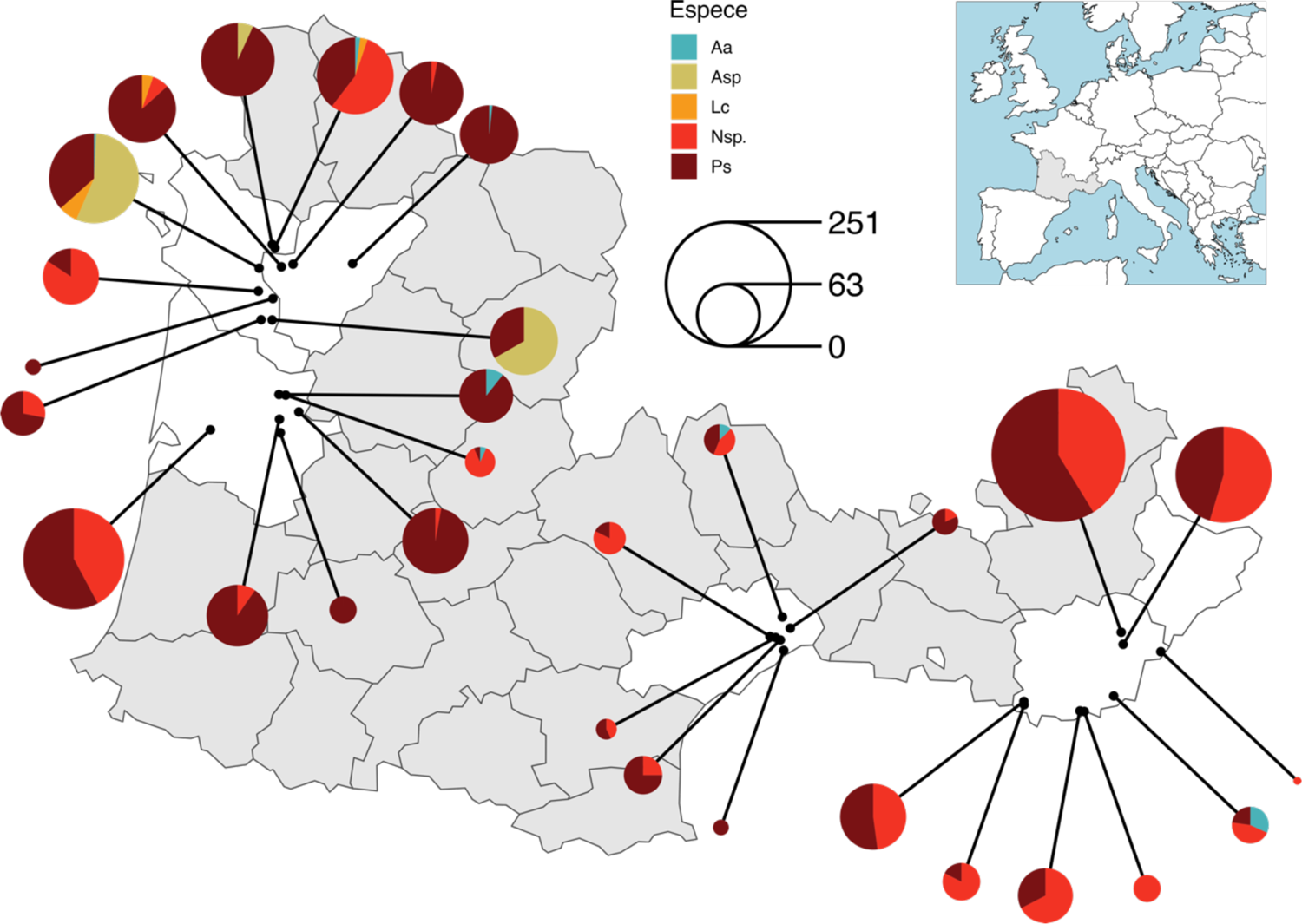
Distribution of insects sampled per buffer in the spring 2021. Insect species are abbreviated as follows Aa: *Aphrophora alni*, Agrs: *Aphrophora* grp. *salicina*, Cv: *Cicadella viridis*, Lc: *Lepyronia coleoptrata*, Nc: *Neophilaenus campestris*, Nl: *Neophilaenus lineatus*, Nsp.: *Neophilaenus* sp. and Ps: *Philaenus spumarius*.

**Figure S1.3.**
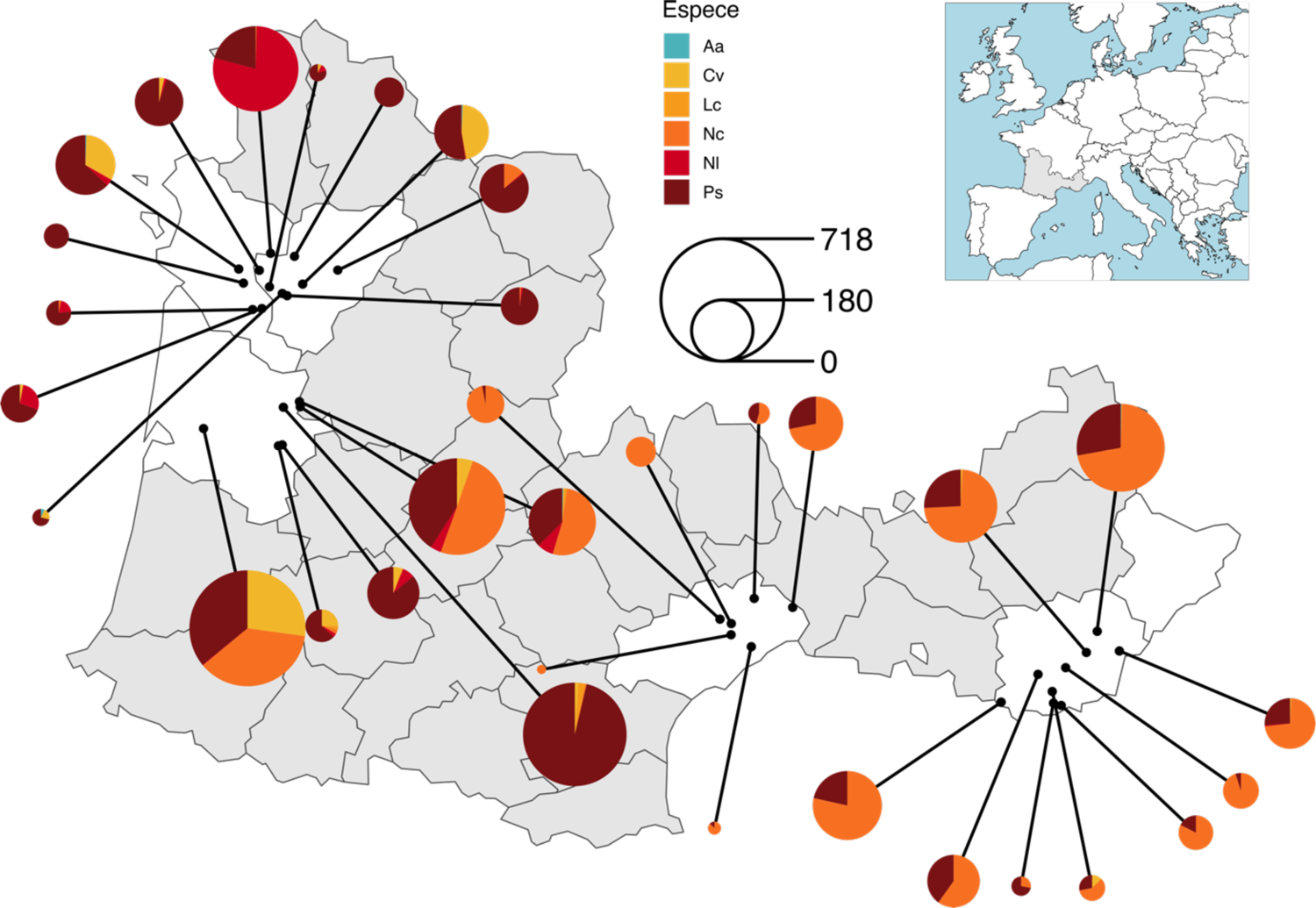
Distribution of insects sampled per buffer in the fall 2021. Insect species are abbreviated as follows Aa: *Aphrophora alni*, Agrs: *Aphrophora* grp. *salicina*, Cv: *Cicadella viridis*, Lc: *Lepyronia coleoptrata*, Nc: *Neophilaenus campestris*, Nl: *Neophilaenus lineatus*, Nsp.: *Neophilaenus* sp. and Ps: *Philaenus spumarius*.

**Figure S1.4.**
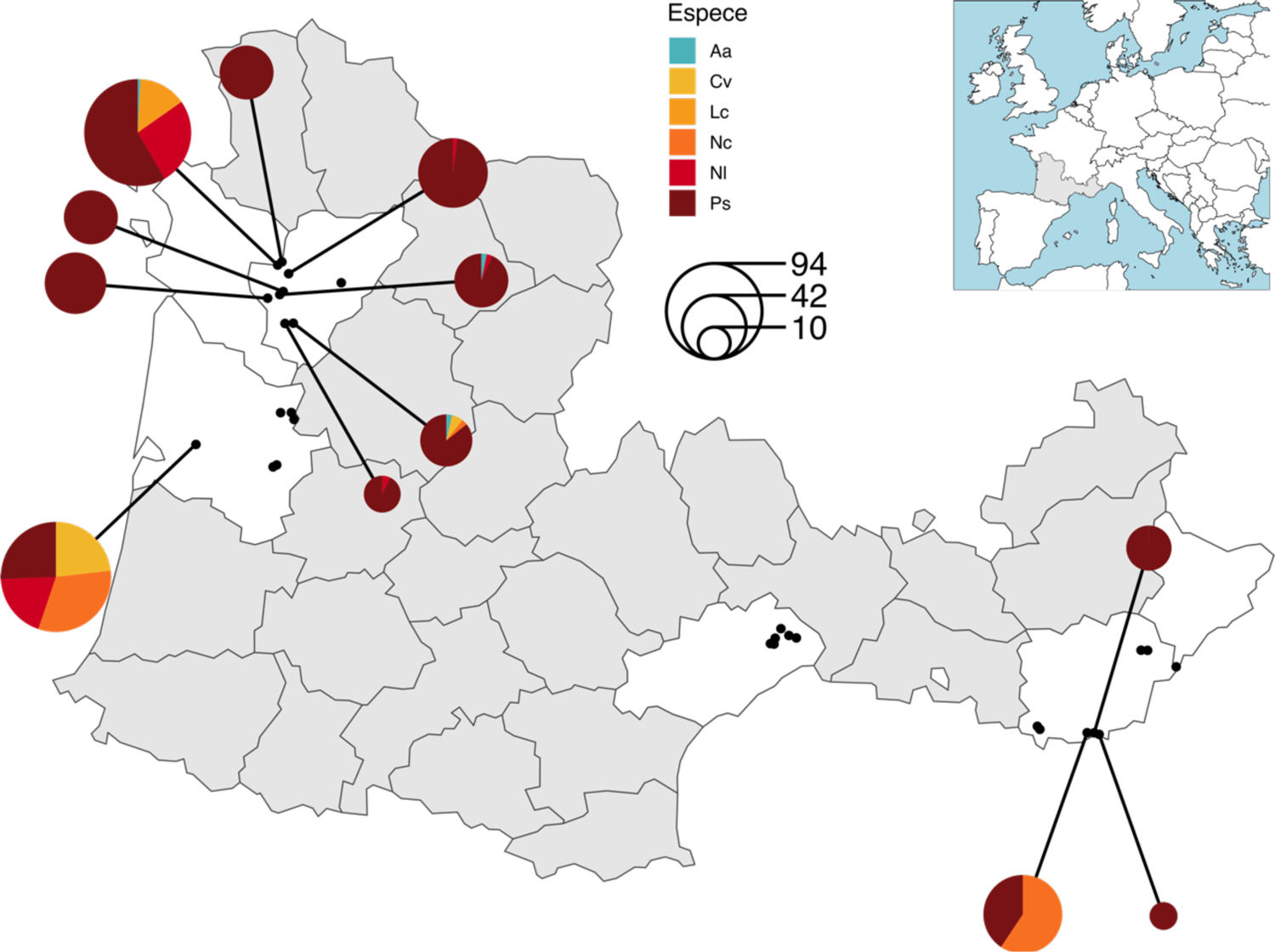
Distribution of insects sampled per buffer in the fall 2020 and screened in molecular biology for the presence of *Xf*. Insect species are abbreviated as follows Aa: *Aphrophora alni*, Agrs: *Aphrophora* grp. *salicina*, Cv: *Cicadella viridis*, Lc: *Lepyronia coleoptrata*, Nc: *Neophilaenus campestris*, Nl: *Neophilaenus lineatus*, Nsp.: *Neophilaenus* sp. and Ps: *Philaenus spumarius*.

**Figure S1.5.**
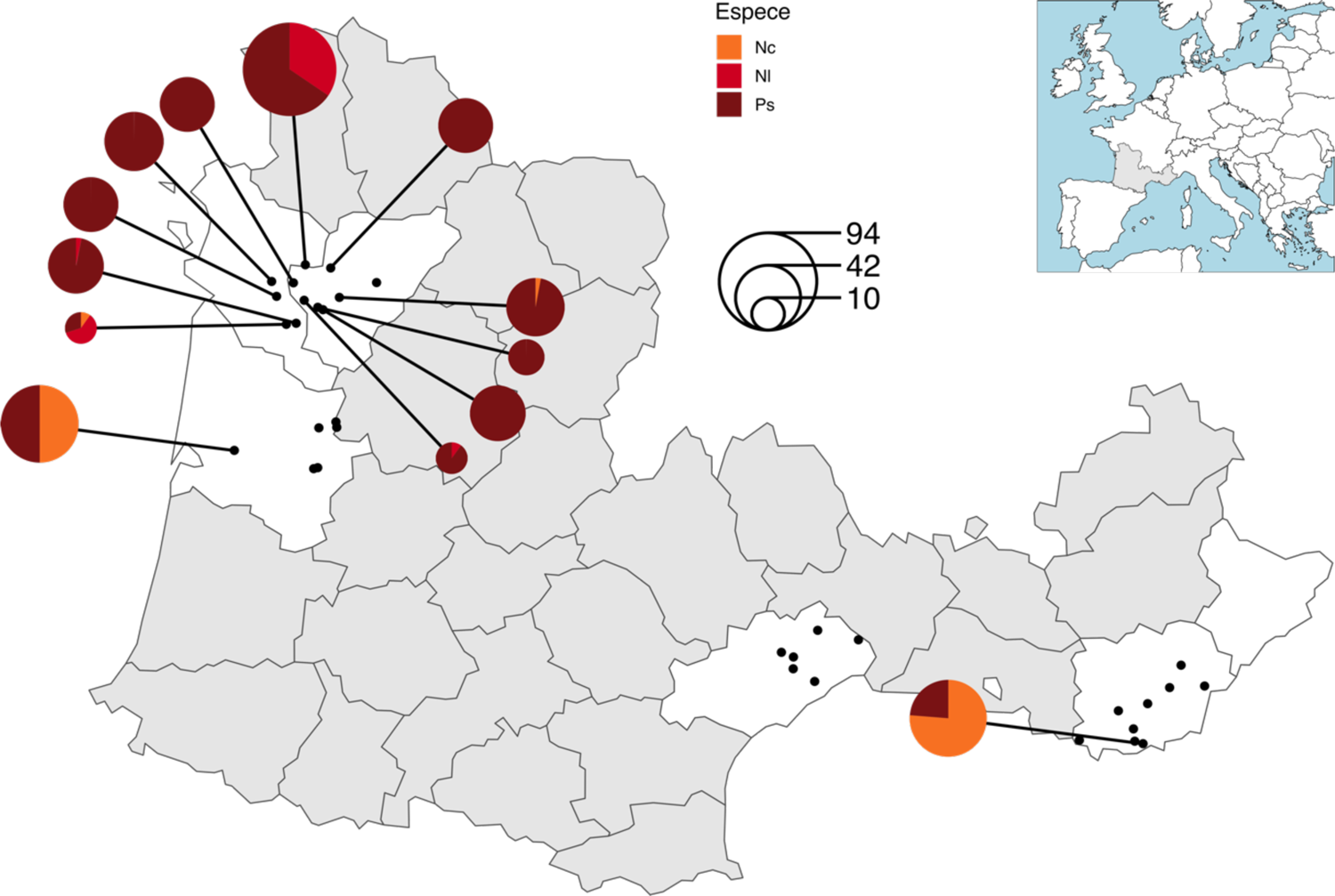
Distribution of insects sampled per buffer in the fall 2021 and screened in molecular biology for the presence of *Xf*. Insect species are abbreviated as follows Aa: *Aphrophora alni*, Agrs: *Aphrophora* grp. *salicina*, Cv: *Cicadella viridis*, Lc: *Lepyronia coleoptrata*, Nc: *Neophilaenus campestris*, Nl: *Neophilaenus lineatus*, Nsp.: *Neophilaenus* sp. and Ps: *Philaenus spumarius*.

**Table S1.1.** Distribution of the insects screened in molecular biology for the presence of *Xf* per habitat and insect species. Insect species are abbreviated as follows Aa: *Aphrophora alni*, Agrs: *Aphrophora* grp. *salicina*, Cv: *Cicadella viridis*, Lc: *Lepyronia coleoptrata*, Nc: *Neophilaenus campestris*, Nl: *Neophilaenus lineatus*, Nsp.: *Neophilaenus* sp. and Ps: *Philaenus spumarius*.

| Session | Region | Land cover type | Stratum | Aa | Cv | Lc | Nc | NI | Ps |
| --- | --- | --- | --- | --- | --- | --- | --- | --- | --- |
| Fall_2020 | NAQ | Border | Lower | 0 | 0 | 2 | 1 | 3 | 149 |
|  |  | Border | Upper | 1 | 0 | 0 | 0 | 0 | 0 |
|  |  | Forest | Lower | 0 | 13 | 0 | 8 | 6 | 0 |
|  |  | Forest | Upper | 1 | 0 | 0 | 1 | 0 | 1 |
|  |  | Alfalfa | Lower | 0 | 0 | 0 | 0 | 0 | 18 |
|  |  | Meadow | Lower | 1 | 17 | 15 | 30 | 38 | 130 |
|  |  | Vine | Lower | 0 | 0 | 0 | 0 | 0 | 15 |
|  |  | Vine | Upper | 0 | 0 | 0 | 0 | 0 | 1 |
|  |  | Bonus habitat: shrubland |  | 0 | 1 | 0 | 1 | 11 | 0 |
|  | PACA | Border | Lower | 0 | 0 | 0 | 2 | 0 | 11 |
|  |  | Forest | Lower | 0 | 0 | 0 | 0 | 0 | 1 |
|  |  | Forest | Upper | 0 | 0 | 0 | 0 | 0 | 1 |
|  |  | Alfalfa | Lower | 0 | 0 | 0 | 0 | 0 | 2 |
|  |  | Meadow | Lower | 0 | 0 | 0 | 30 | 0 | 20 |
|  |  | Riparian | Lower | 0 | 0 | 0 | 0 | 0 | 13 |
|  |  | Vine | Lower | 0 | 0 | 0 | 5 | 0 | 7 |
|  |  | Vine | Upper | 0 | 0 | 0 | 1 | 0 | 0 |
| Fall_2021 | NAQ | Border | Lower | 0 | 0 | 0 | 2 | 38 | 79 |
|  |  | Alfalfa | Lower | 0 | 0 | 0 | 0 | 0 | 90 |
|  |  | Meadow | Lower | 0 | 0 | 0 | 30 | 0 | 150 |
|  |  | Vine | Lower | 0 | 0 | 0 | 0 | 0 | 12 |
|  | PACA | Border | Lower | 0 | 0 | 0 | 44 | 0 | 13 |
|  |  | Vine | Lower | 0 | 0 | 0 | 1 | 0 | 1 |
| Total |  |  |  | 3 | 31 | 17 | 156 | 96 | 714 |

**Table S1.2.**
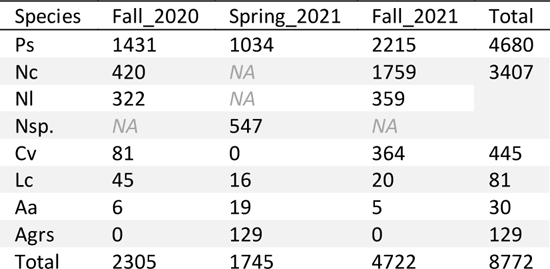
Number of xylem feeders sampled each session, all sites combined. Insect species are abbreviated as follows Aa: *Aphrophora alni*, Agrs: *Aphrophora* grp. *salicina*, Cv: *Cicadella viridis*, Lc: *Lepyronia coleoptrata*, Nc: *Neophilaenus campestris*, Nl: *Neophilaenus lineatus*, Nsp.: *Neophilaenus* sp. and Ps: *Philaenus spumarius*. At the nymph stage (Spring_2021 session) *N. campestris* and *N. lineatus* were found morphologically indistinguishable, so nymphs were analyzed at the genus level (*Neophilaenus* sp.). As *C. viridis* produces no spittle at the nymph stage, no *C. viridis* were collected in Spring.

| Species | Fall_2020 | Spring_2021 | Fall_2021 | Total |
| --- | --- | --- | --- | --- |
| Ps | 1431 | 1034 | 2215 | 4680 |
| Nc | 420 | NA | 1759 | 3407 |
| NI | 322 | NA | 359 |  |
| Nsp. | NA | 547 | NA |  |
| Cv | 81 | 0 | 364 | 445 |
| Lc | 45 | 16 | 20 | 81 |
| Aa | 6 | 19 | 5 | 30 |
| Agrs | 0 | 129 | 0 | 129 |
| Total | 2305 | 1745 | 4722 | 8772 |

### Appendix 2: Assessment of vegetation cover on quadrats

**Figure S2.1.**
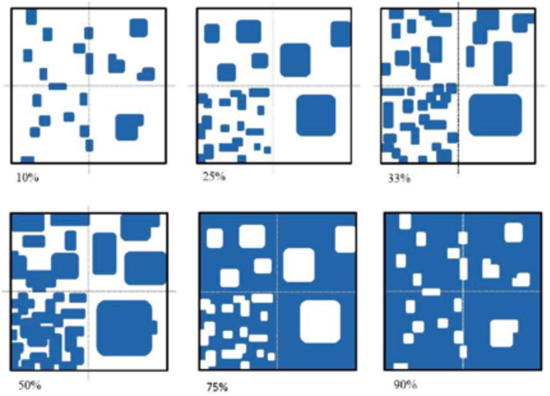
Abacus used to estimate quadrat vegetation cover

**Table S2.1.**
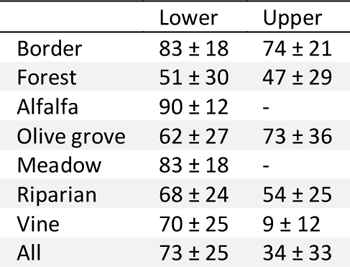
Vegetation cover per land-use type (lines) and stratum (columns). Figures stand for Means ± Standard deviation. “-” stands for “stratum always unavailable”. “All” is computed on all quadrats and is not the average of table lines.

### Appendix 3: Network layout to capture the sample size of habitats using ‘plotweb’ function, package ‘bipartite’

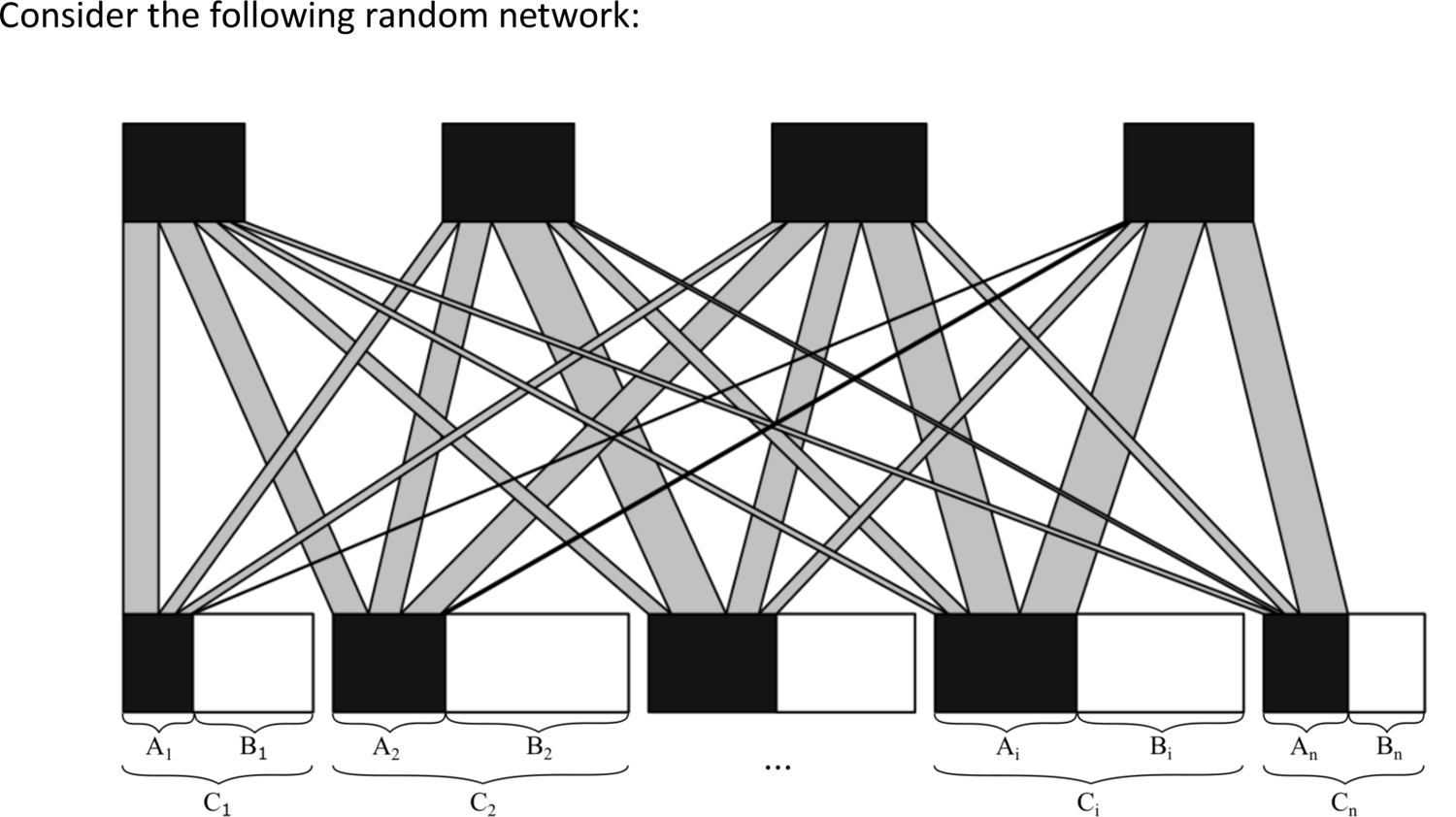

*A*_i_ is the raw number of insects sampled in habitat *i*. B_i_ is computed to include sample size information in the network based on constraints we fixed. First we define:

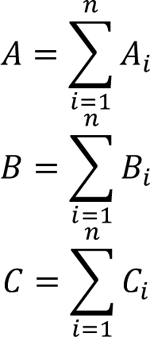

For our insect-habitat networks, we wanted the “resource-level” boxes to be scaled such as: 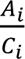 represents the proportion of non-empty (with at least one insect) samples taken in habitat i, i.e. 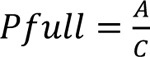. The same holds at the scale of the whole resource level, i.e. 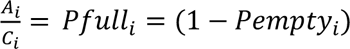 where *Pfull* is the proportion, all habitats together, of non-empty (with at least one insect) samples.

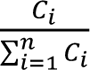 represents the proportion of subsites sampled in habitat *i* among the *n* subsites sampled, i.e. 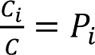

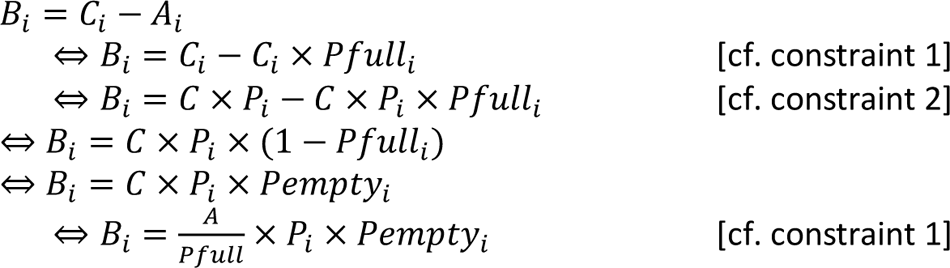

*A, Pfull, P_i_*, and *Pempty_i_* are all available from the data. So for each network we computed B_i_ and used it as the ‘low.abun’ argument in ‘plotweb’ function.

### Appendix 4: Generalized linear mixed models details

#### Supplementary methods: modeling procedure

GLMM parameters were chosen to be as parsimonious as possible, while following basic modelling guidelines, checked using package ‘DHARMa’ (Hartig, 2020). Therefore, the model architecture was complexified only if the guidelines were not followed at the earlier step. For all models we followed the steps:

- We fitted a model that followed the basic modelling guidelines with all fixed terms mentioned in the main text (see below for the parameters used depending on the type of data).
- We computed the analysis of deviance table (function ‘Anova’, package ‘car’).
- If all fixed terms were significant the modelling procedure stopped.
- If at least one fixed term was insignificant, we removed the least significant term and went back to step 1 with the simplified fixed term formula. The same parameters were kept for the new model, unless adjustments were necessary to follow modelling guidelines.

Continuous data (Normalized degree, Resource Range and PDI) were analyzed using the following sequence of parameters:

- Gaussian distribution with an identity link
- If unsatisfying, Student distribution with an identity link
- If unsatisfying, Student distribution with a log link
- If unsatisfying, Tweedie distribution with a log link

Count data (raw insect abundance) were analyzed using the following sequence of parameters:

- Poisson distribution with a log link
- If unsatisfying, Negative binomial distribution linearly parameterized (“nbinom1”) with a log link
- If unsatisfying, Negative binomial distribution linearly parameterized (“nbinom2”) with a log link
- If unsatisfying, we compared the 12 following models and used the one that best followed DHARMa tests and that produced non-NA Anova.

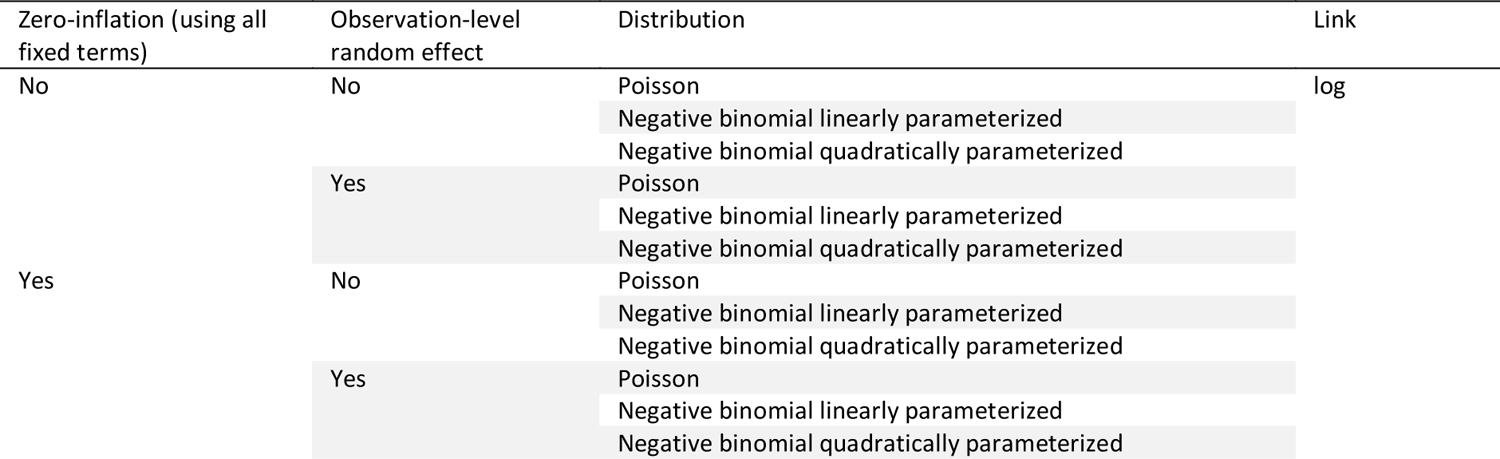

**Table S4.1.** Details of the results of the GLMM performed on habitat, plant family and plant species specialization of insects. “-“ stands for “unsignificant variable removed from the model”.

| Question | Network used | Response variable | Network metric computed on... | Family (final model) | Link (final model) | Variable | Chi2 | df | P-val | R<sup>2</sup><sub>m</sub> |
| --- | --- | --- | --- | --- | --- | --- | --- | --- | --- | --- |
| Insect specialization on habitat | Insect species - habitat network for each region and each session | All seven specialization metrics:<br>• ressource range<br>• paired difference index<br>• d'<br>• species specificity index<br>• z-score of ressource range<br>• z-score of paired difference index<br>• z-score of species specificity index | Present / absent and quantified interactions | Student | identity | Species | 27.79 | 7 | < 0.001 | 0.19 |
|  |  |  |  |  |  | Metric | 191.40 | 6 | < 0.001 |  |
| Insect specialization on plant family | Insect species - plant family network for each region and each session (nymph data only) |  |  | Student | log | Species | 91.60 | 4 | < 0.001 | 0.94 |
|  |  |  |  |  |  | Metric | 51.01 | 6 | < 0.001 |  |
| Insect specialization on plant species | Insect species - plant species network for each region and each session (nymph data only) |  |  | Gaussian | identity | Species | 16.64 | 4 | 0.002 | 0.18 |
|  |  |  |  |  |  | Metric | 27.80 | 6 | < 0.001 |  |

**Table S4.2.**
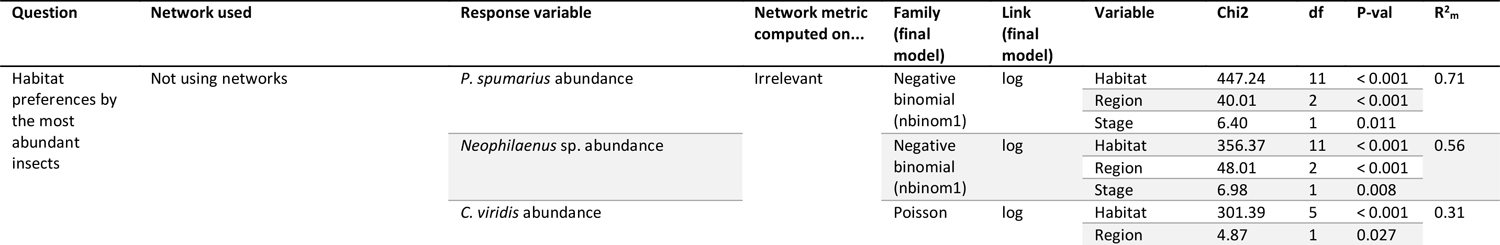
Details of the results of the GLMM performed on *P. spumarius*, *Neophilaenus* sp. and *C. viridis* abundances. “-“ stands for “unsignificant variable removed from the model”. For *Cicadella viridis* abundance model, the random effect on the site ID was not included because it was never found in the upper stratum of any site, and the fixed term “Stage” was not included as it was not found at the nymph stage (no spittle produced by this species).

**Table S4.3.** Details of the results of the GLMM performed on. “-“ stands for “unsignificant variable removed from the model”. For *Cicadella viridis* abundance model, the random effect on the site ID was not included because it was never found in the upper stratum of any site, and the fixed term “Stage” was not included as it was not found at the nymph stage (no spittle produced by this species).

| Question | Network used | Response variable | Network metric computed on... | Family (final model) | Link (final model) | Variable | Chi2 | df | P-val | R<sup>2</sup><sub>m</sub> |
| --- | --- | --- | --- | --- | --- | --- | --- | --- | --- | --- |
| Habitat exploitation by insects | Not using networks | Xylem feeder abundance in each habitat |  | Negative binomial (nbinom1) | log | Habitat | 477.20 | 11 | < 0.001 | 0.65 |
|  |  |  |  |  |  | Region | 25.56 | 2 | < 0.001 |  |
|  |  |  |  |  |  | Stage | 23.90 | 1 | < 0.001 |  |
|  |  | Effective number of species (exponential of Shannon index) |  | Student | identity | Habitat | 21.50 | 5 | < 0.001 | 0.07 |
|  |  |  |  |  |  | Region | 32.52 | 2 | < 0.001 |  |
|  |  |  |  |  |  | Stage | - | - | - |  |

**Table S4.4.**
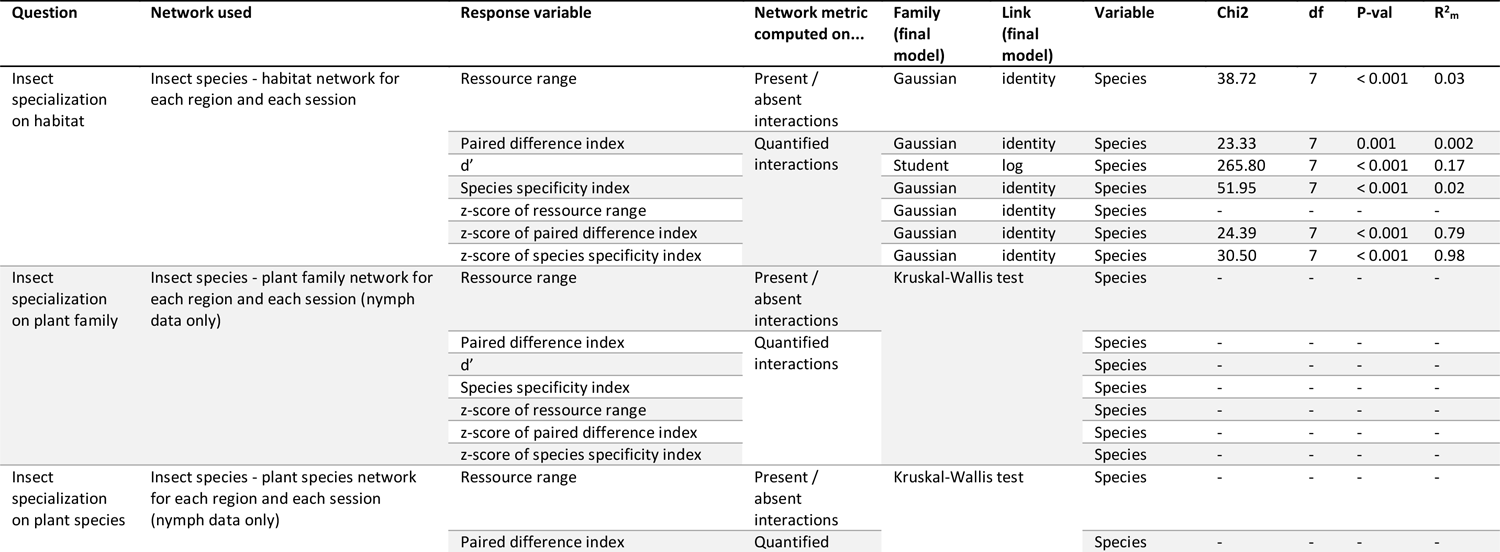
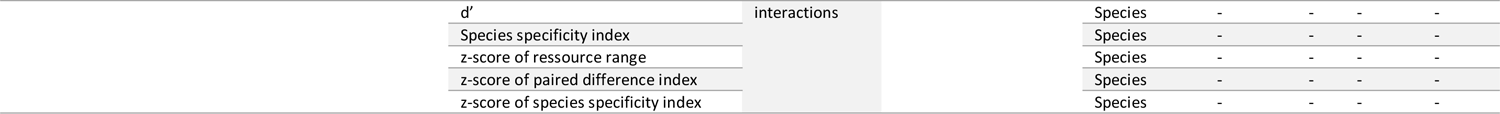
In addition to the analysis reported in Table S4.1, each specialization metric was also analyzed using GLMMs for insect-habitat networks and using Kruskal-Wallis tests (Hollander et al., 2014) for insect-plant families and insect-plant species networks (insufficient number of data to use GLMMs). The formula (1) in main text was then simplified to keep only the species-fixed effect. Details of these analyses are shown below. “-“ stands for “unsignificant variable removed from the model”.

### Appendix 5: Details on specialization analyses

**Figure S5.1.**
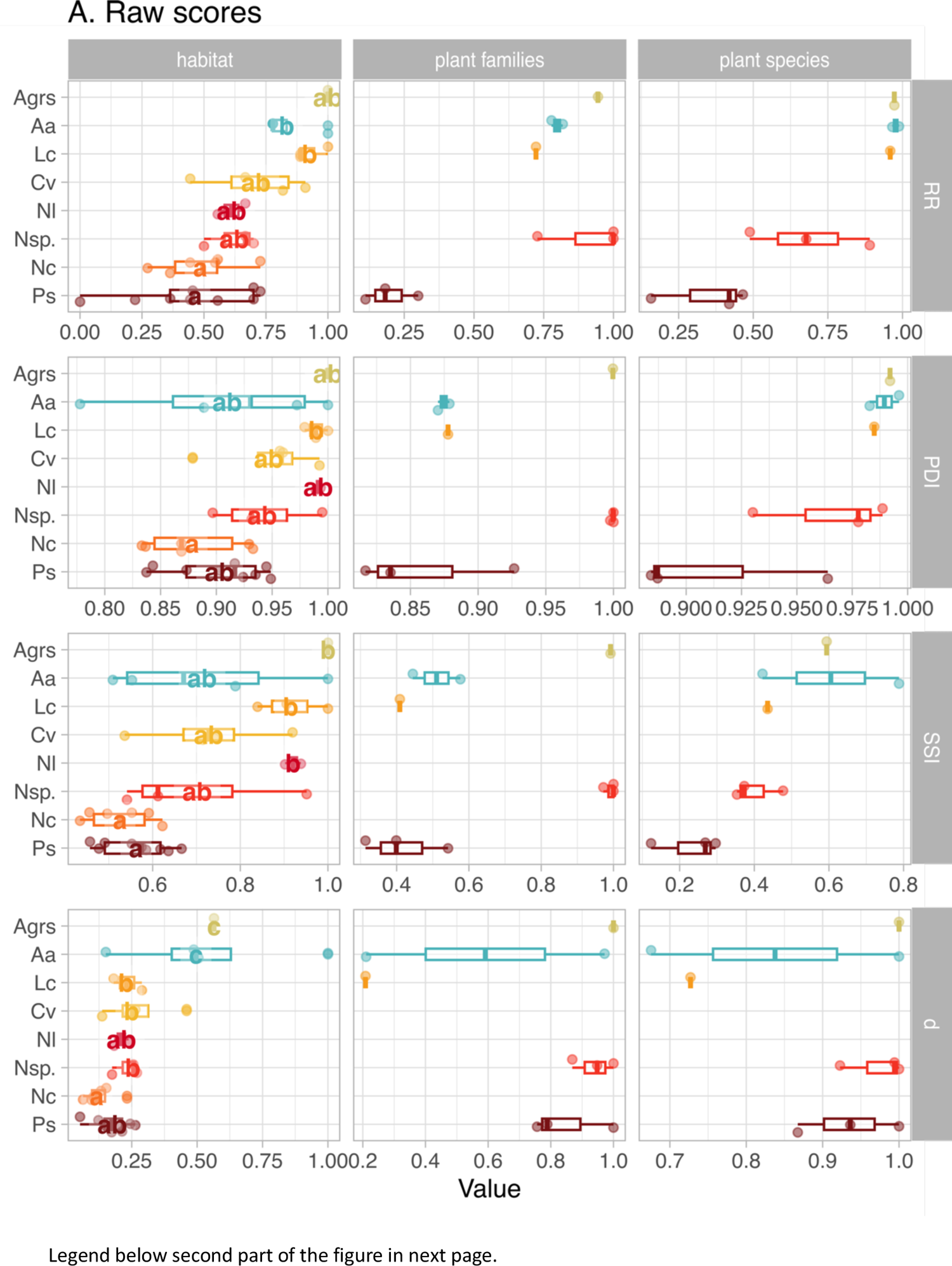
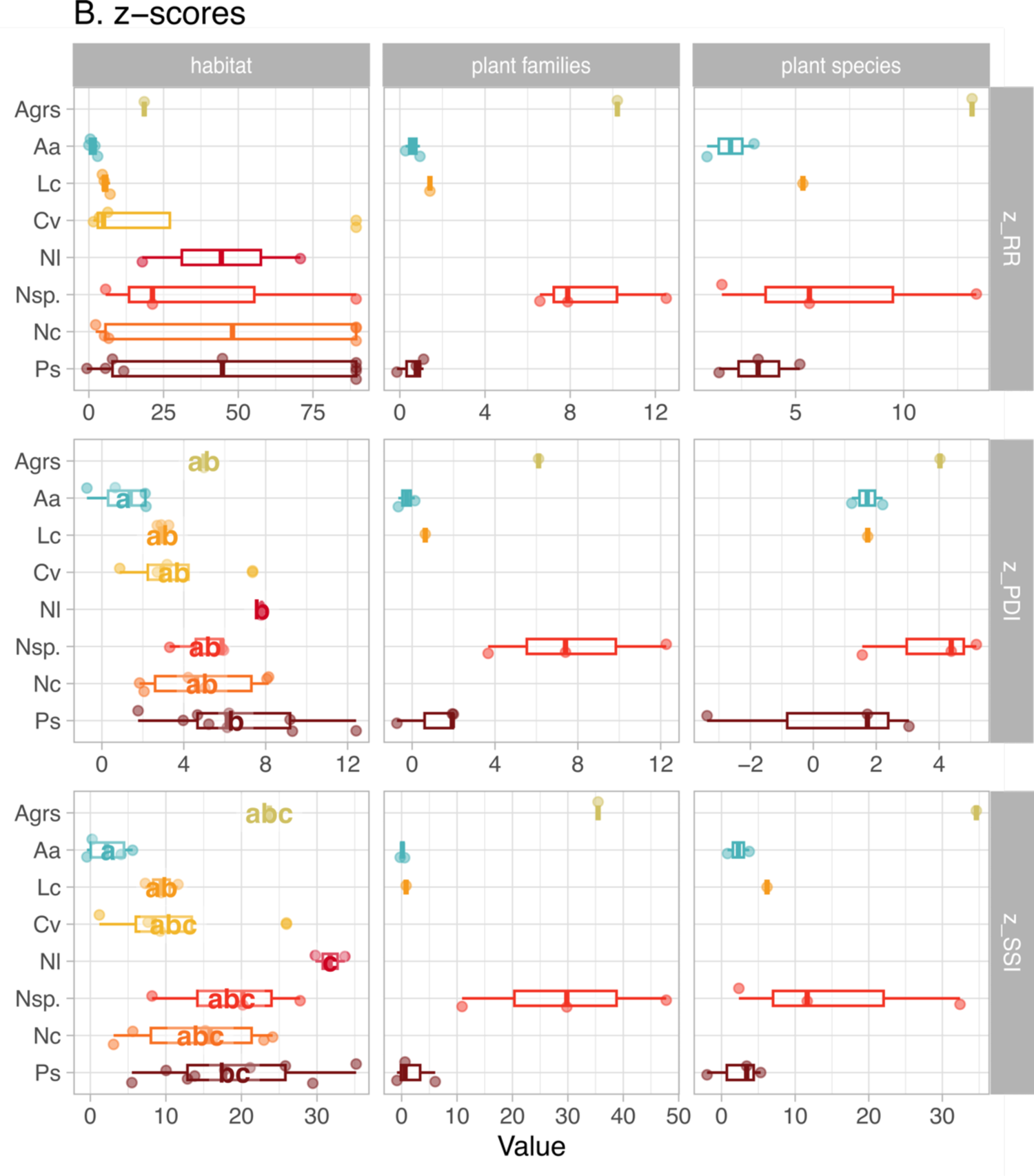
Distribution of specialization metrics for each ressource-level network, each metric, and each insect species. X scales are free, meaning that all specialization values are scaled relatively to the range of values in their panel. Fig. 3 in the main text is an overlap of all panels for each column. Panels including letters indicate networks and metrics for which the fixed effect of insect species was significant. In each panel independently, species sharing a letter do not differ significantly. See Table S4.4 for details on the statistics.

### Appendix 6: Details on habitat preferences of *P. spumarius*, *Neophilaenus* sp. and *C. viridis*

**Figure S6.1.**
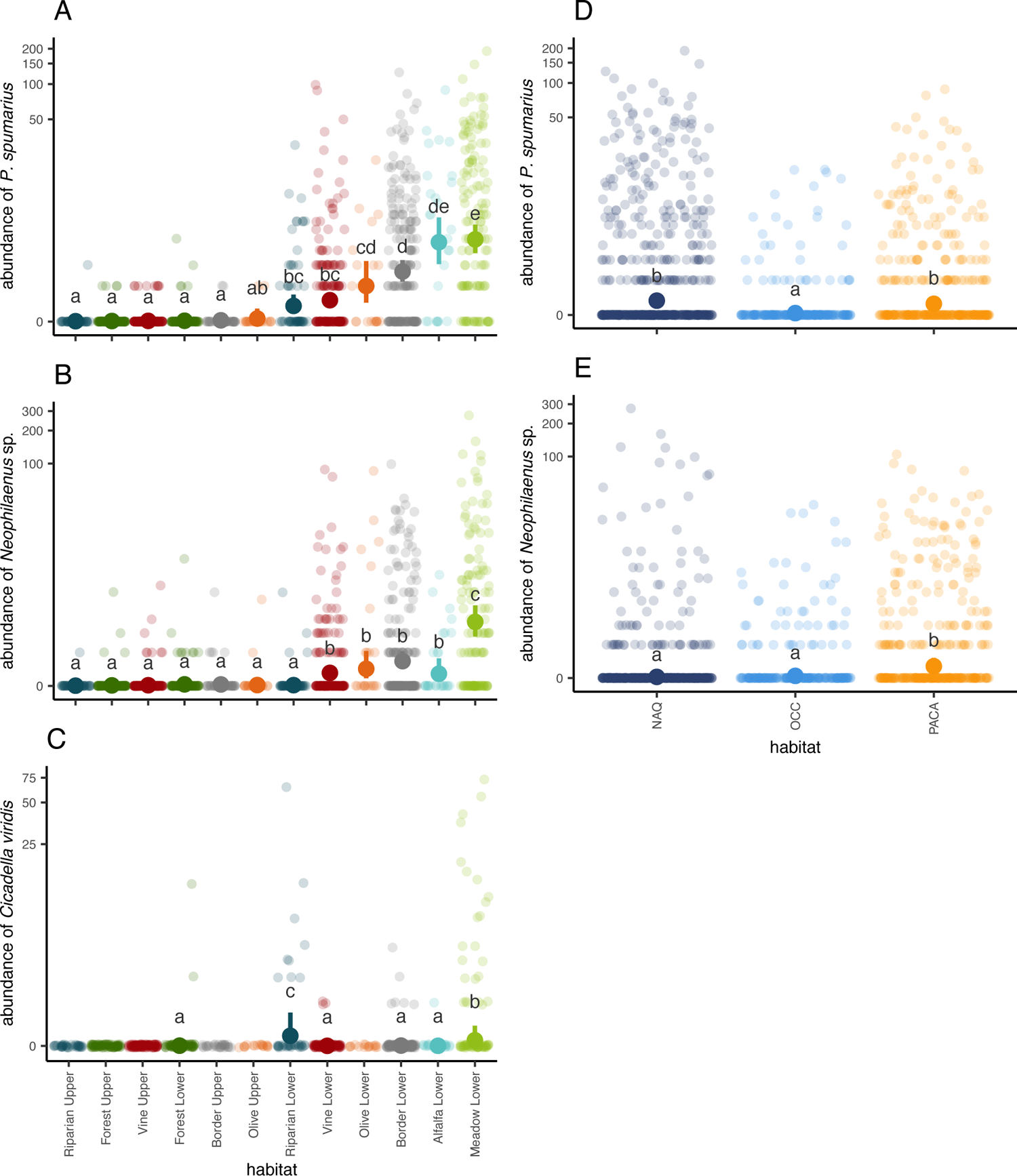
Scatterplots of raw data and estimated marginal means predicted by the model fitted on *P. spumarius* (A, D) *Neophilaenus* sp. (B, E) and *C. viridis* (C) abundance, pairwise association between habitats (A, B, C) and between regions (D, E). Abundances in habitats or regions sharing a letter do not differ significantly. Habitats that had invariably null values (for *C. viridis*) could not be compared to others in the statistical analysis (hence no letters).

### Appendix 7: Insect-habitat interaction networks colored by insect species

**Figure S7.1.**
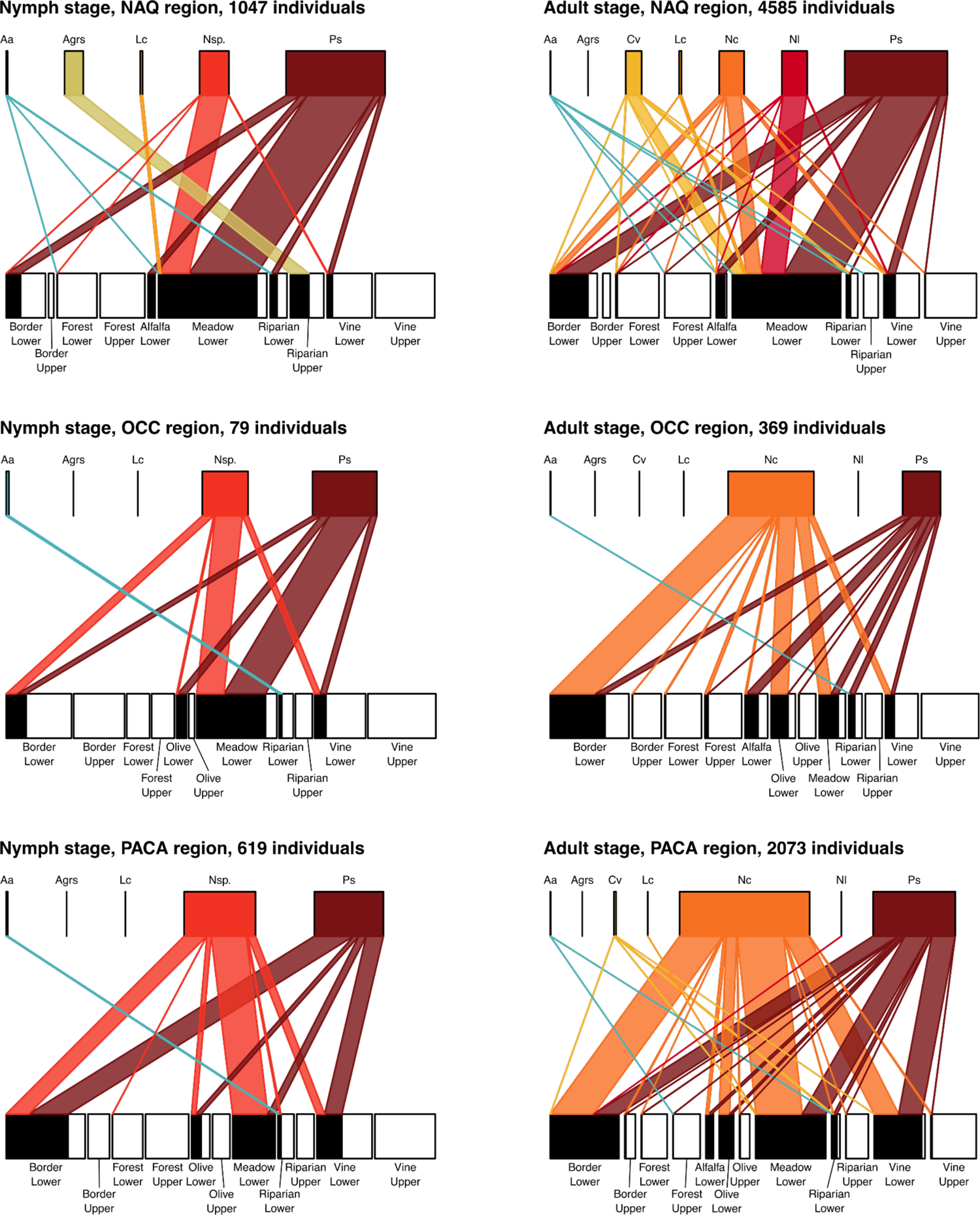
Insect habitat network with links colored by insect species. Insect species are abbreviated as follows Aa: *Aphrophora alni*, Agrs: *Aphrophora* grp. *salicina*, Cv: *Cicadella viridis*, Lc: *Lepyronia coleoptrata*, Nc: *Neophilaenus campestris*, Nl: *Neophilaenus lineatus*, Nsp.: *Neophilaenus* sp. and Ps: *Philaenus spumarius*.

### Appendix 8: Insect-plant species interaction networks with full details on plant species

**Figure S8.1.**
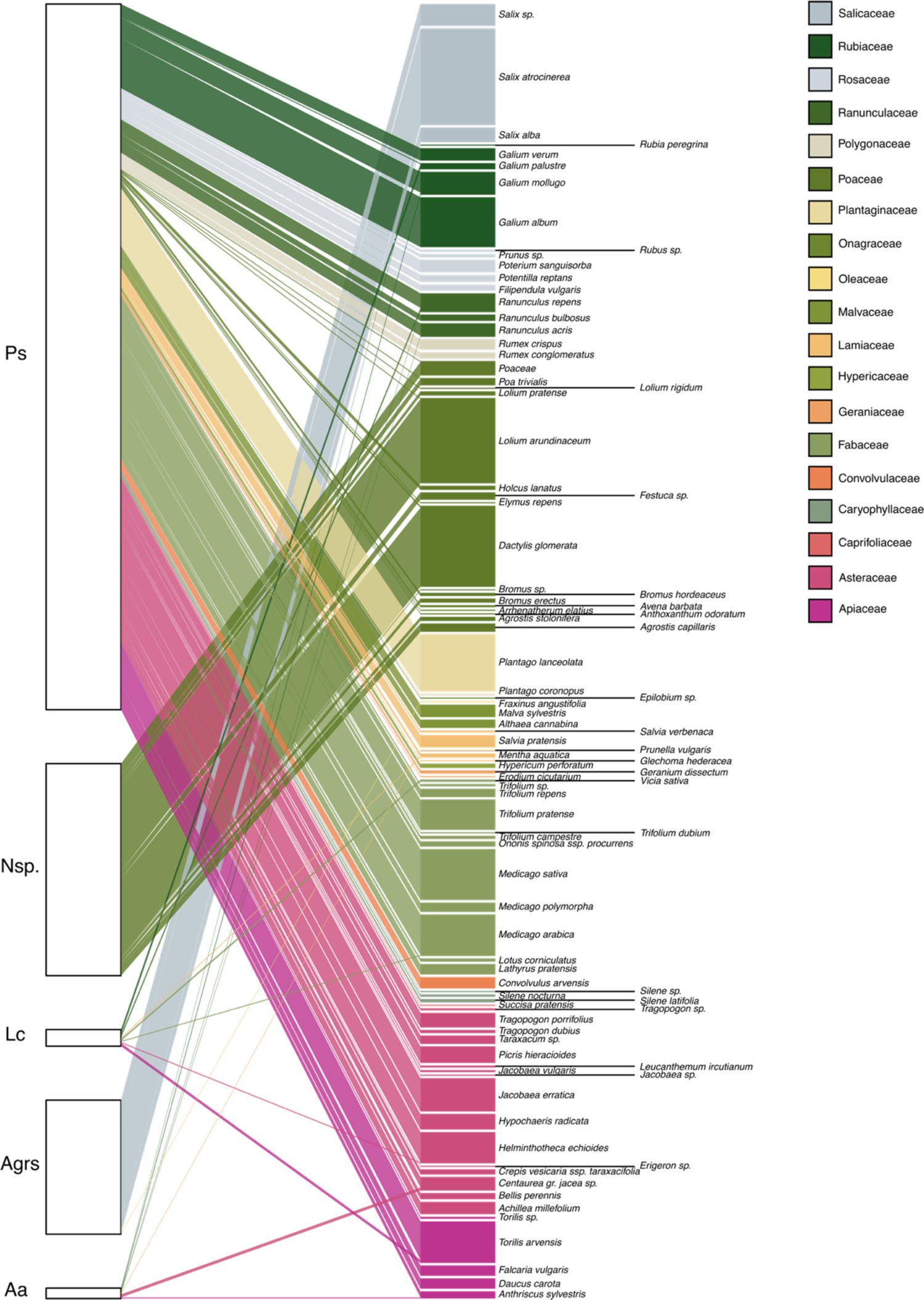
Insect plant network in NAQ region at the nymph stage (spring 2021). Insect species are abbreviated as follows Aa: *Aphrophora alni*, Agrs: *Aphrophora* grp. *salicina*, Cv: *Cicadella viridis*, Lc: *Lepyronia coleoptrata*, Nc: *Neophilaenus campestris*, Nl: *Neophilaenus lineatus*, Nsp.: *Neophilaenus* sp. and Ps: *Philaenus spumarius*.

**Figure S8.2.**
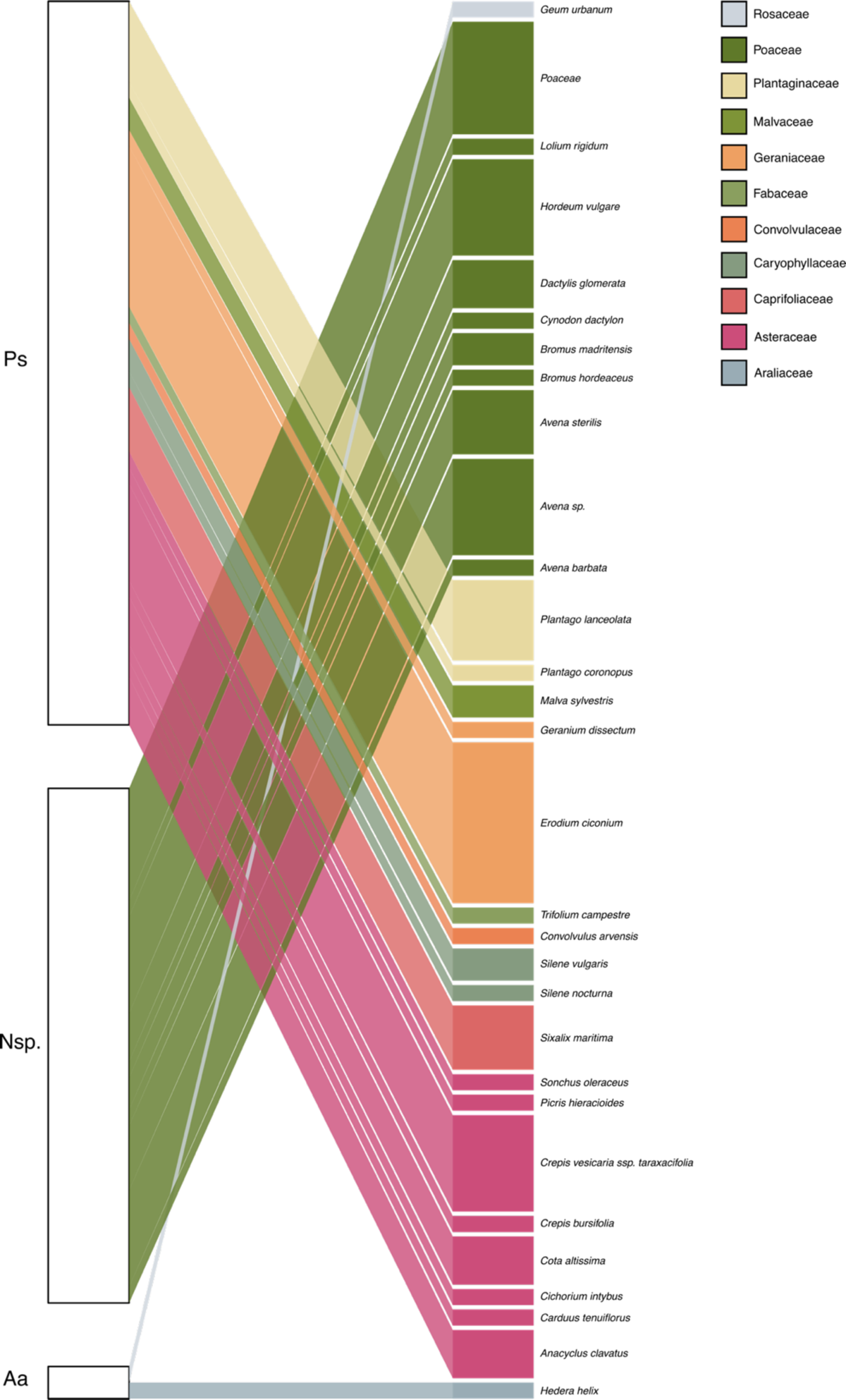
Insect plant network in OCC region at the nymph stage (spring 2021). Insect species are abbreviated as follows Aa: *Aphrophora alni*, Agrs: *Aphrophora* grp. *salicina*, Cv: *Cicadella viridis*, Lc: *Lepyronia coleoptrata*, Nc: *Neophilaenus campestris*, Nl: *Neophilaenus lineatus*, Nsp.: *Neophilaenus* sp. and Ps: *Philaenus spumarius*.

**Figure S8.3.**
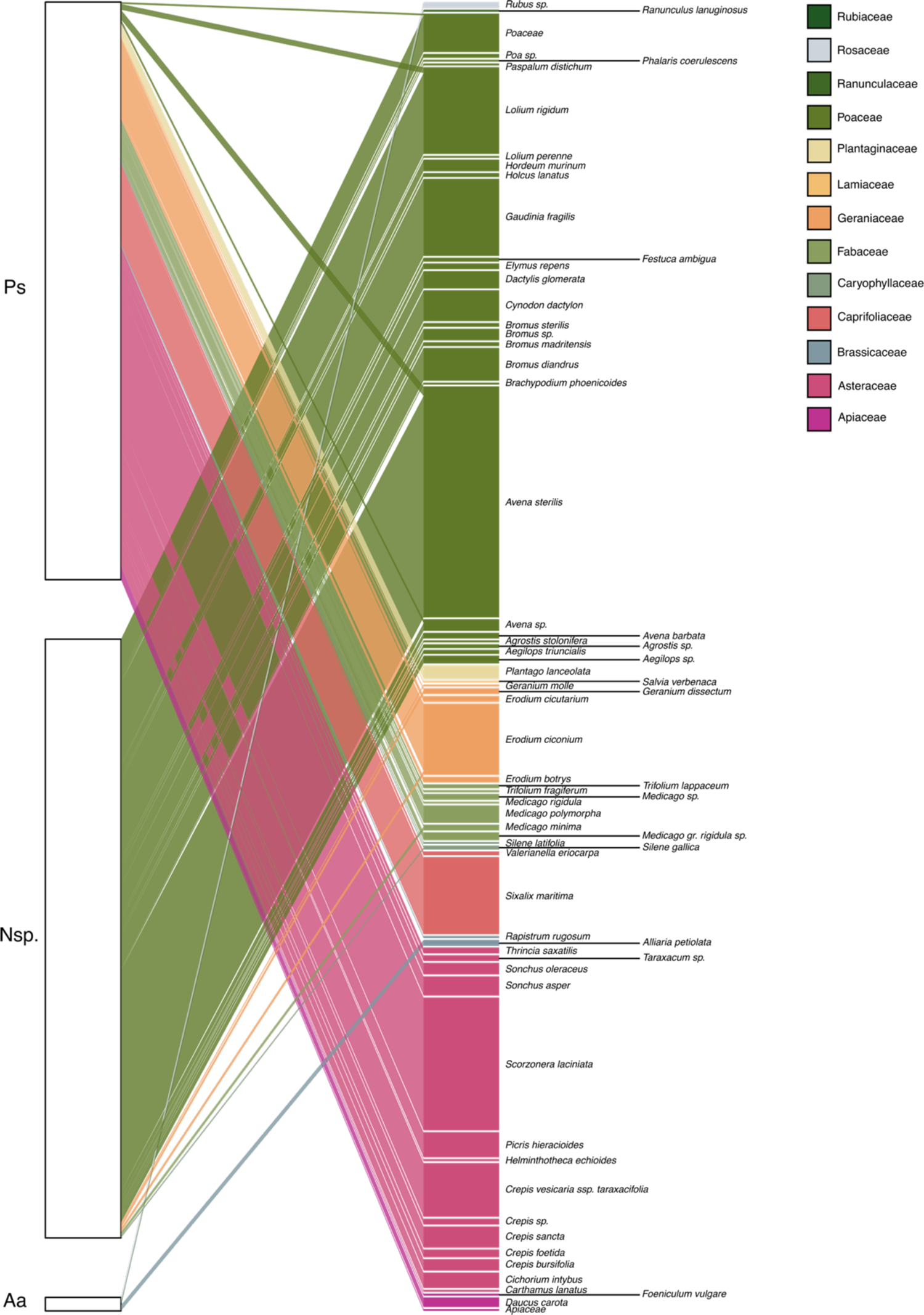
Insect plant network in PACA at the nymph stage (spring 2021). Insect species are abbreviated as follows Aa: *Aphrophora alni*, Agrs: *Aphrophora* grp. *salicina*, Cv: *Cicadella viridis*, Lc: *Lepyronia coleoptrata*, Nc: *Neophilaenus campestris*, Nl: *Neophilaenus lineatus*, Nsp.: *Neophilaenus* sp. and Ps: *Philaenus spumarius*.

**Table S8.1.** Number of plant families and plant species consumed by *P. spumarius* (Ps), *Neophilaenus* sp. (Nsp.), *A. alni* (Aa), *L. coleoptrata* (Lc) and *A.* grp. *salicina* (Agrs). The first three plant species are those on which the highest number of individuals of the species was measured, all sites combined.

| Species | #Plant family | #Plant species | Top 3 plant species |
| --- | --- | --- | --- |
| Ps | 18 | 87 | *Scorzonera laciniata*; *Plantago lanceolata*; *Medicago sativa* |
| Nsp. | 4 | 31 | *Avena sterilis*; *Dactylis glomerata*; *Lolium arundinaceum* |
| Aa | 8 | 10 | *Alliaria petiolata*; *Galium mollugo*; several at 1 individual |
| Lc | 6 | 7 | *Galium palustre*; *Ranunculus repens*; *Torilis arvensis* |
| Agrs | 2 | 3 | *Salix atrocinerea*; *Salix alba*; *Fraxinus angustifolia* |

### Appendix 9: additional data on insect-plant interaction network

Additionally to the regular sampling protocol for nymphs, other samples were made opportunistically, when we spotted spittles on infrequent plants. The networks below display all interactions observed, including those reported in Appendix 5, and additional observations.

Four new plant families appear in these networks:

- Dennstaedtiaceae, a family of ferns, represented here by *Pteridium aquilinum*;
- Fagaceae, represented by *Quercus robur* (the spittle was found on a seedling);
- Cyperaceae, represented by *Carex hirta* and
- Euphorbiaceae represented by *Euphorbia helioscopia*.

**Figure S9.1.**
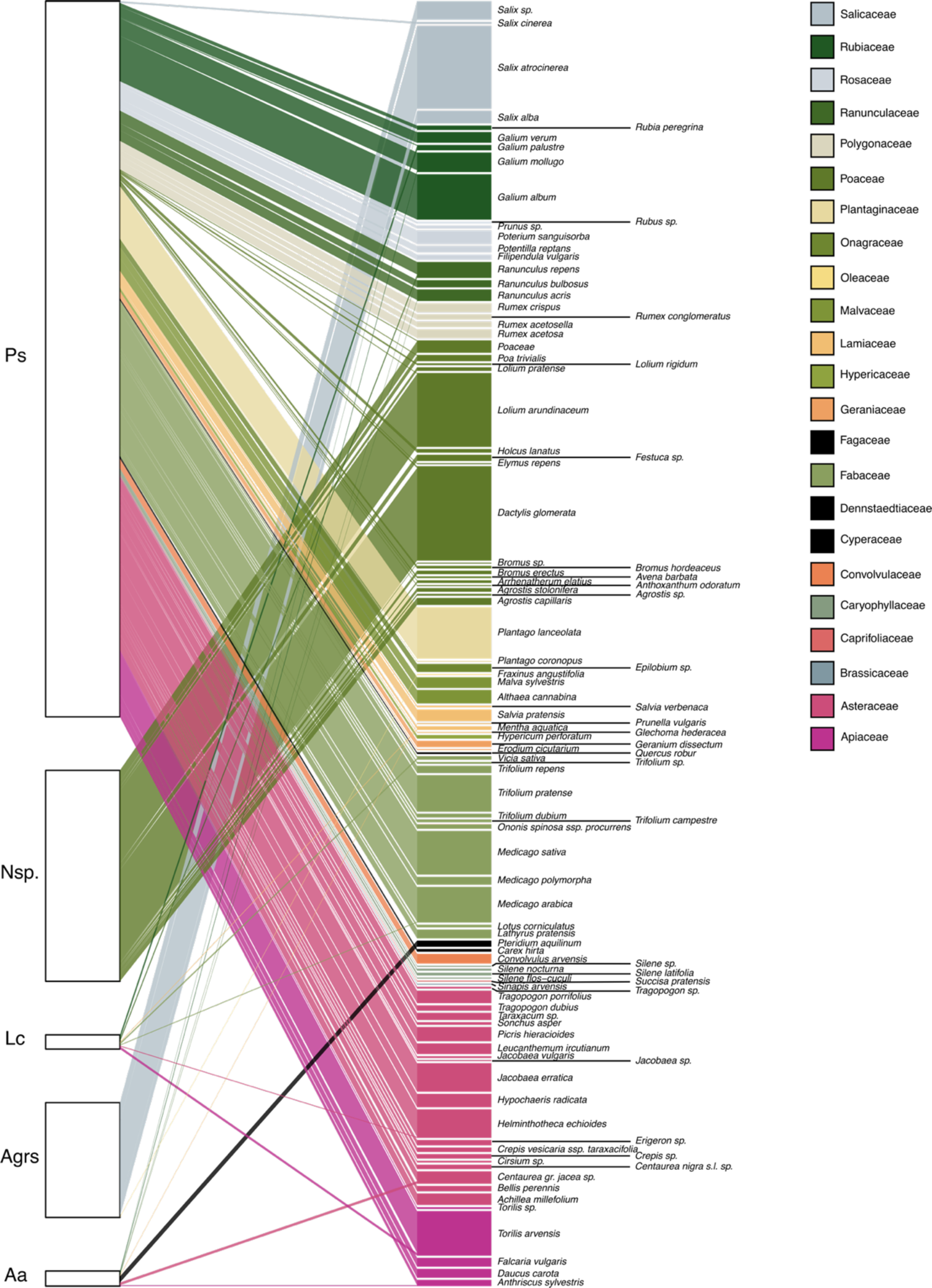
Insect plant network in NAQ at the nymph stage (spring 2021), including regular and additional opportunistic observations. The colors are the same as previously, the four new plant families are depicted in black to stand out (see above for the correspondence species-family). Insect species are abbreviated as follows Aa: *Aphrophora alni*, Agrs: *Aphrophora* grp. *salicina*, Cv: *Cicadella viridis*, Lc: *Lepyronia coleoptrata*, Nc: *Neophilaenus campestris*, Nl: *Neophilaenus lineatus*, Nsp.: *Neophilaenus* sp. and Ps: *Philaenus spumarius*.

**Figure S9.2.**
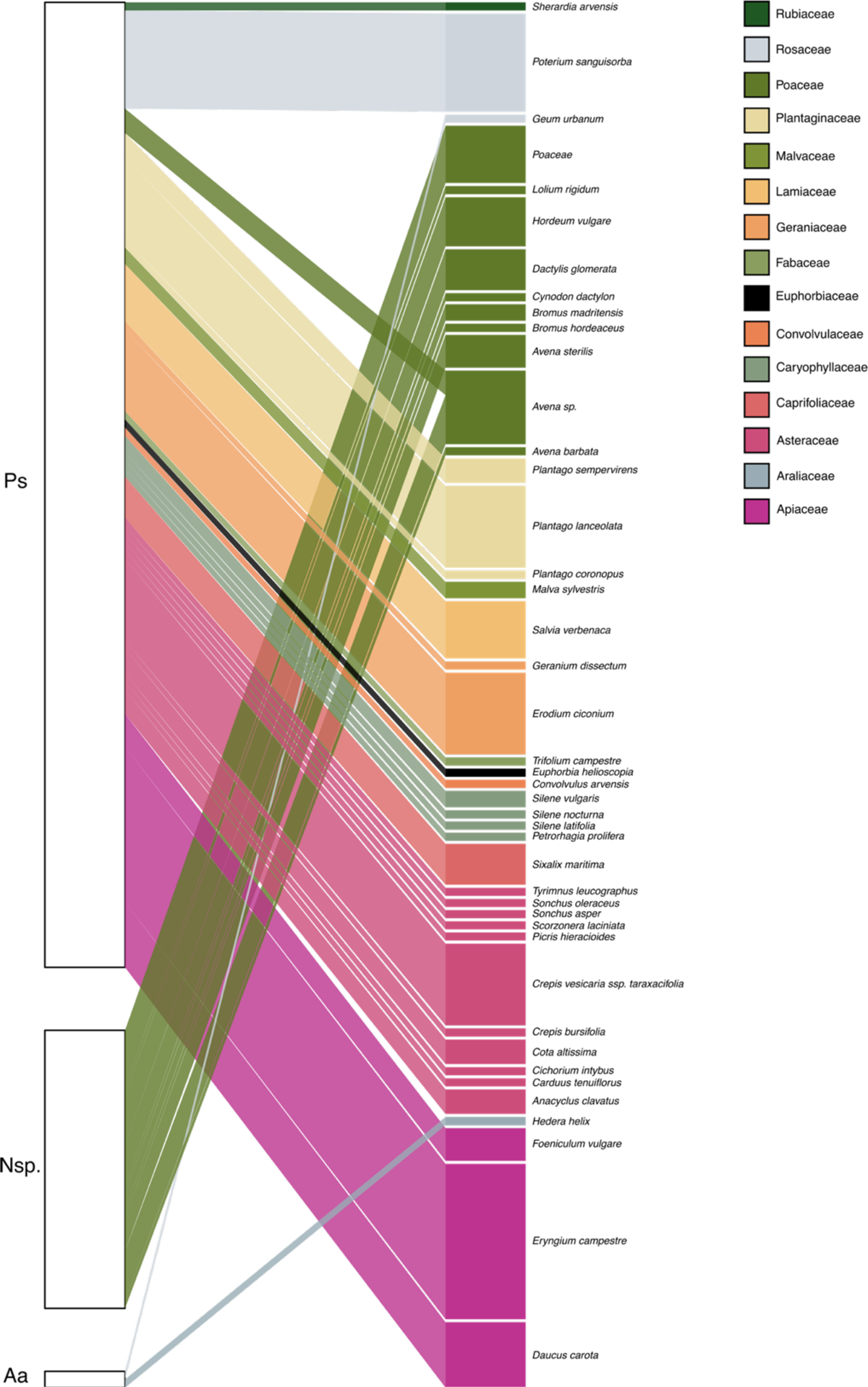
Insect plant network in OCC at the nymph stage (spring 2021), including regular and additional opportunistic observations. The colors are the same as previously, the four new plant families are depicted in black to stand out (see above for the correspondence species-family). Insect species are abbreviated as follows Aa: *Aphrophora alni*, Agrs: *Aphrophora* grp. *salicina*, Cv: *Cicadella viridis*, Lc: *Lepyronia coleoptrata*, Nc: *Neophilaenus campestris*, Nl: *Neophilaenus lineatus*, Nsp.: *Neophilaenus* sp. and Ps: *Philaenus spumarius*.

**Figure S9.3.**
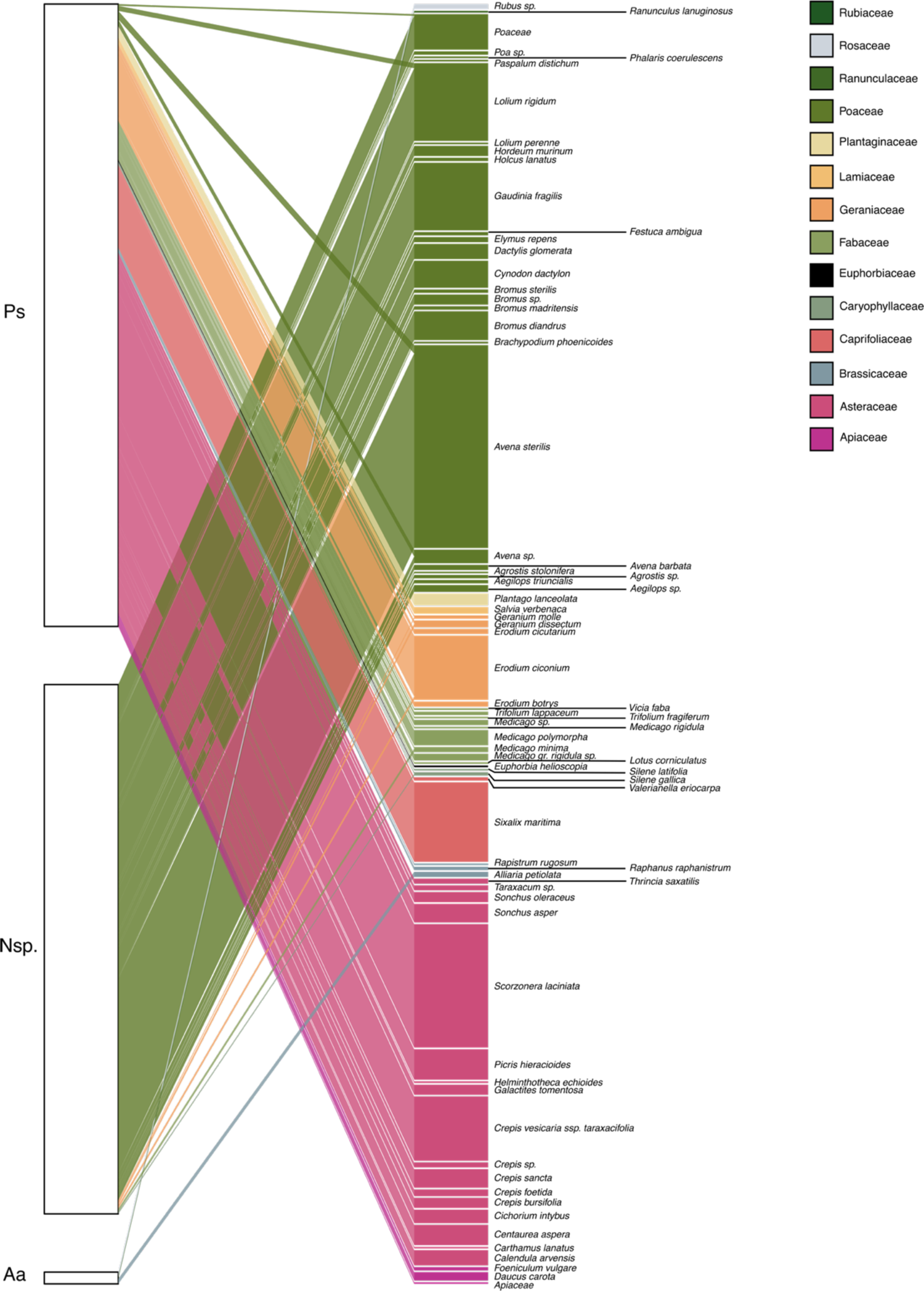
Insect plant network in PACA at the nymph stage (spring 2021), including regular and additional opportunistic observations. The colors are the same as previously, the four new plant families are depicted in black to stand out (see above for the correspondence species-family). Insect species are abbreviated as follows Aa: *Aphrophora alni*, Agrs: *Aphrophora* grp. *salicina*, Cv: *Cicadella viridis*, Lc: *Lepyronia coleoptrata*, Nc: *Neophilaenus campestris*, Nl: *Neophilaenus lineatus*, Nsp.: *Neophilaenus* sp. and Ps: *Philaenus spumarius*.

**Figure S9.4.**
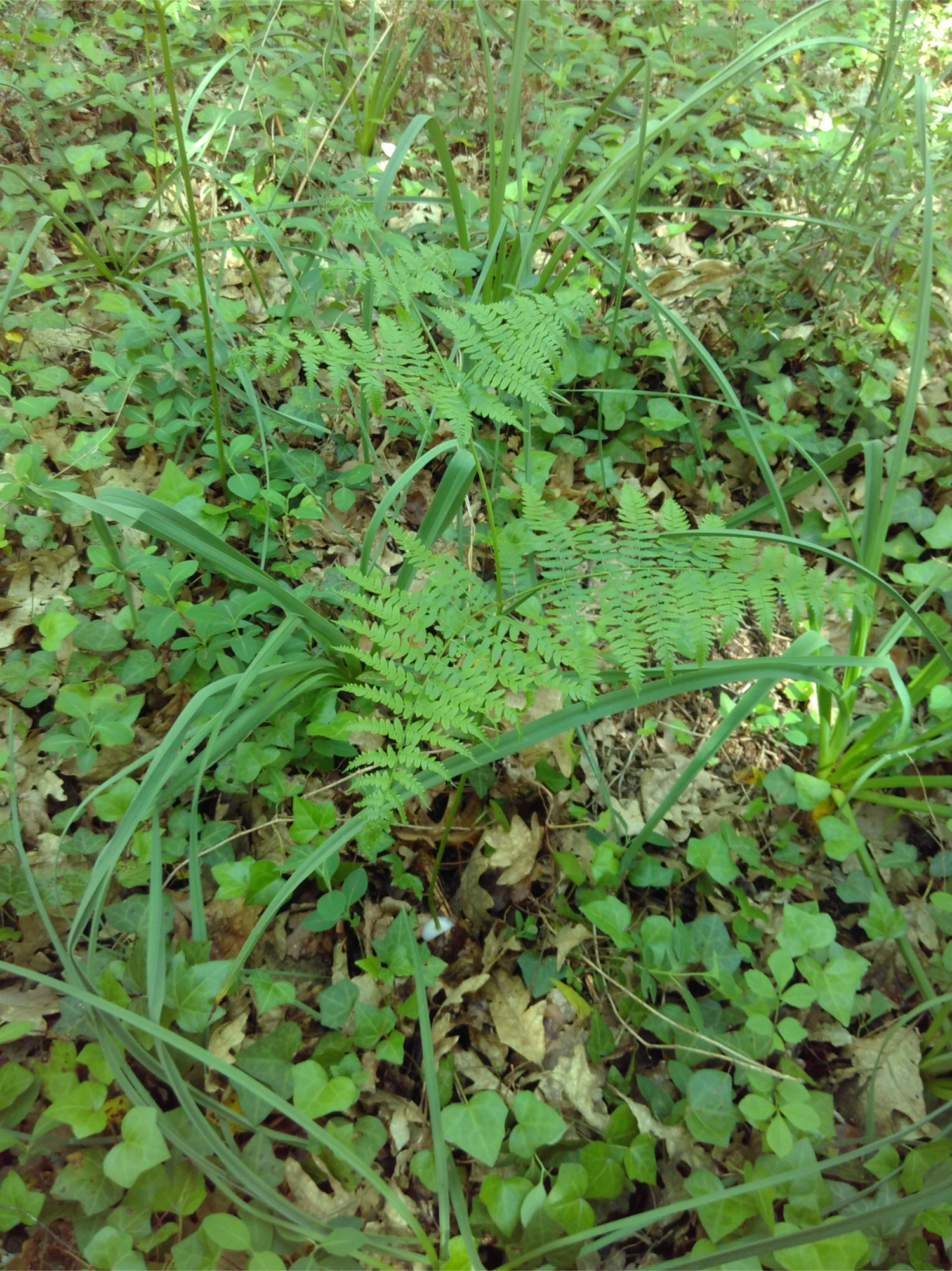
*A. alni* spittle on *P. aquilinum*

### Appendix 10: The host plants of *P. spumarius*

**Table S10.1.**
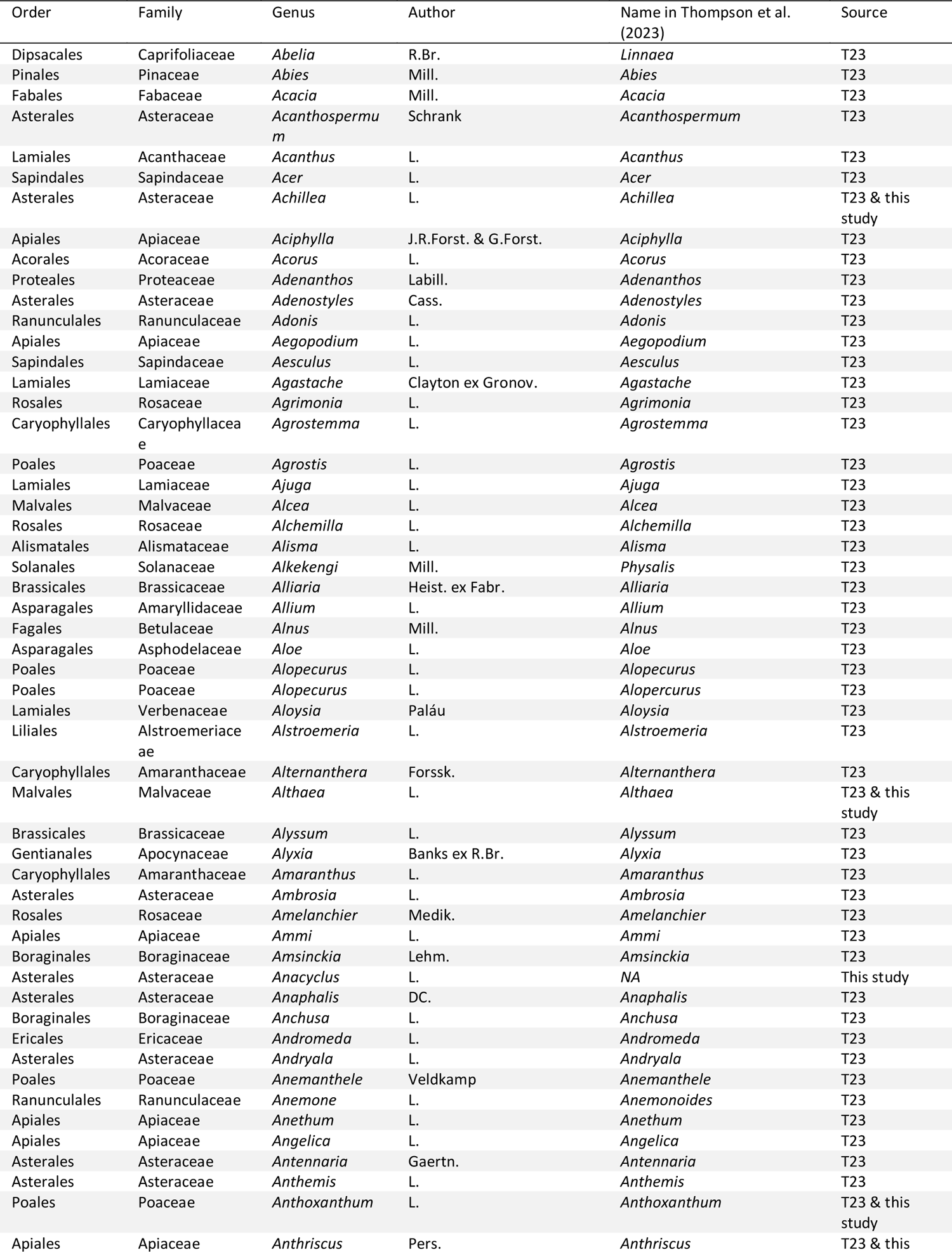
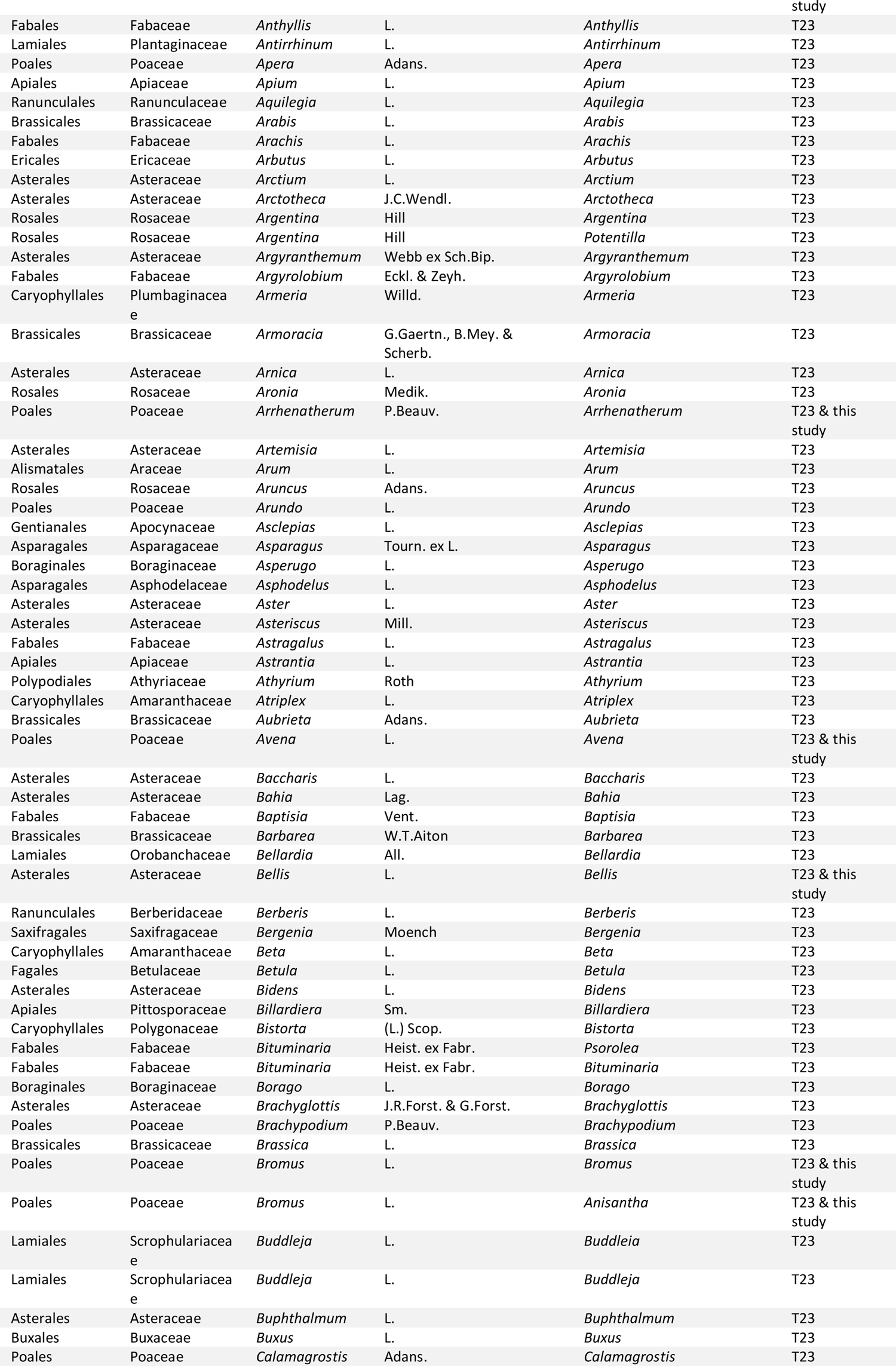
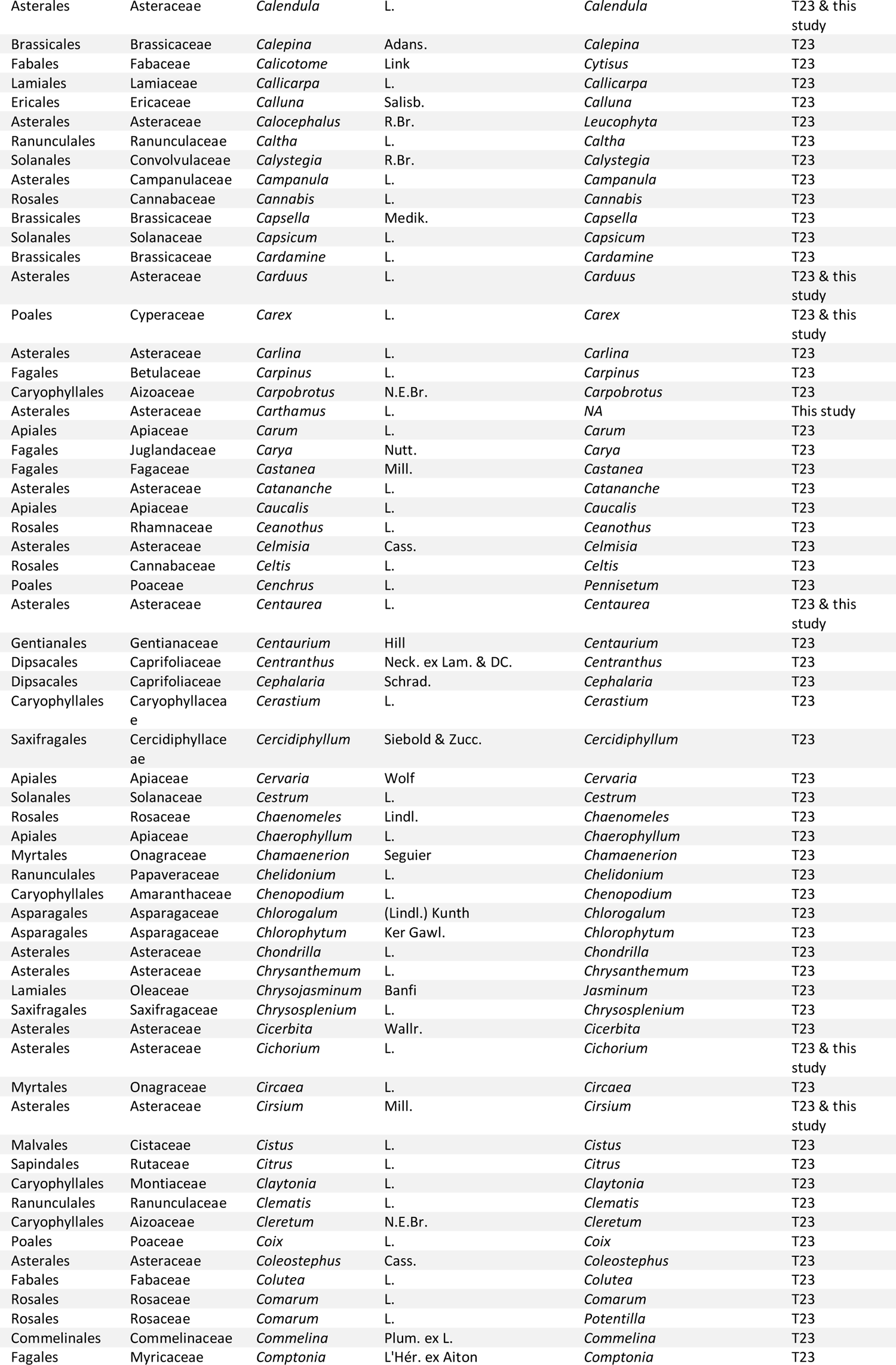
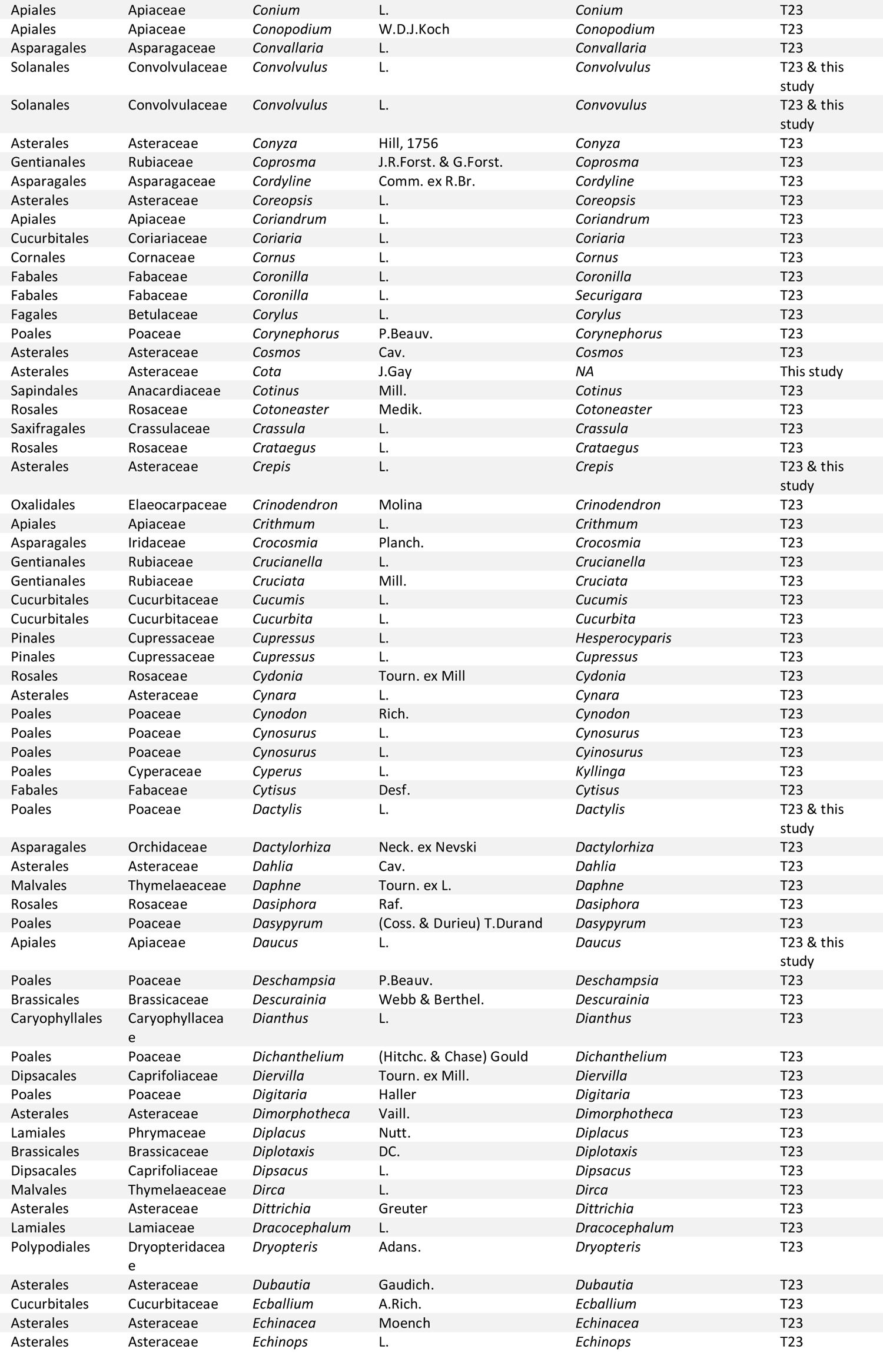
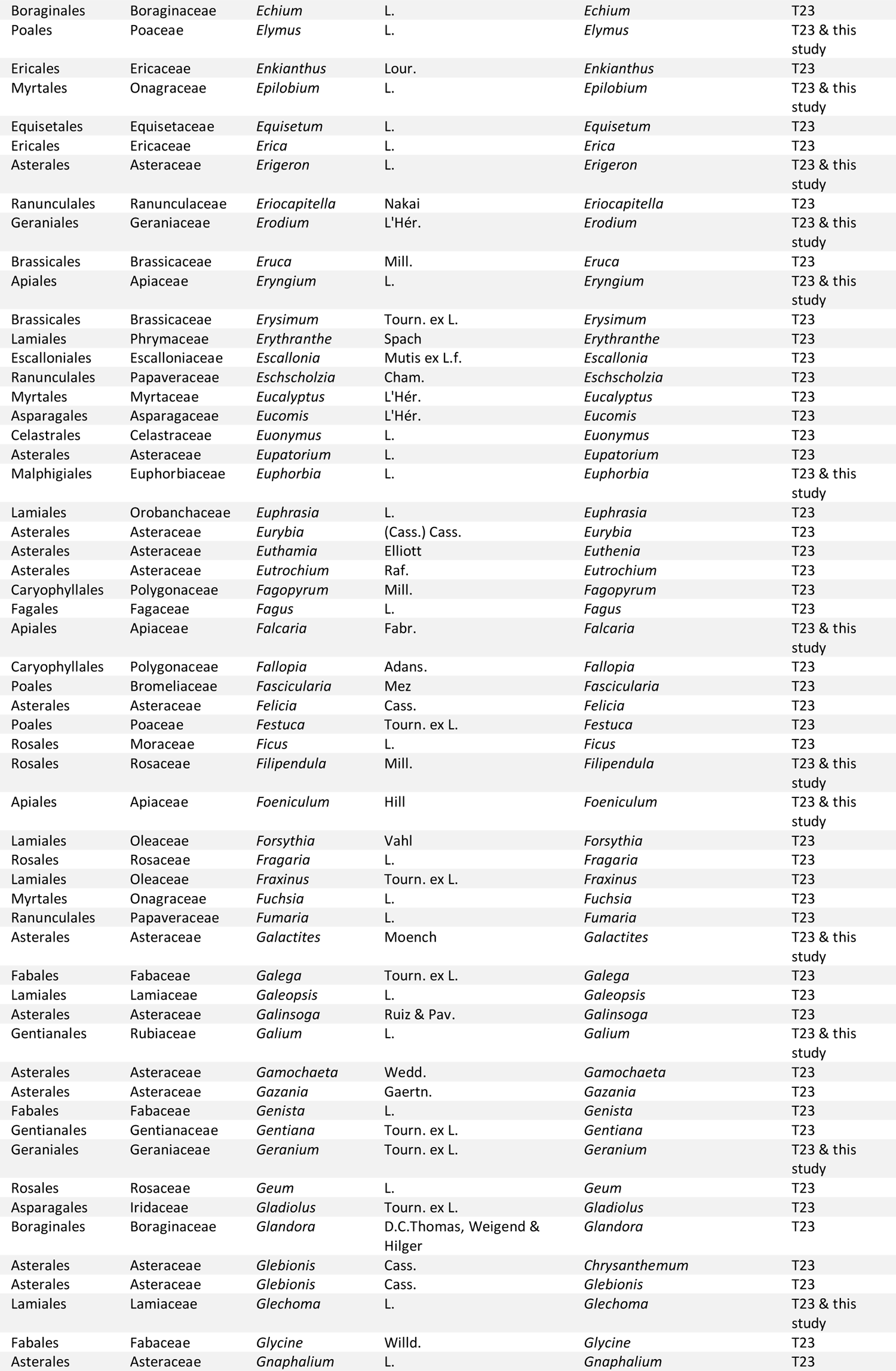
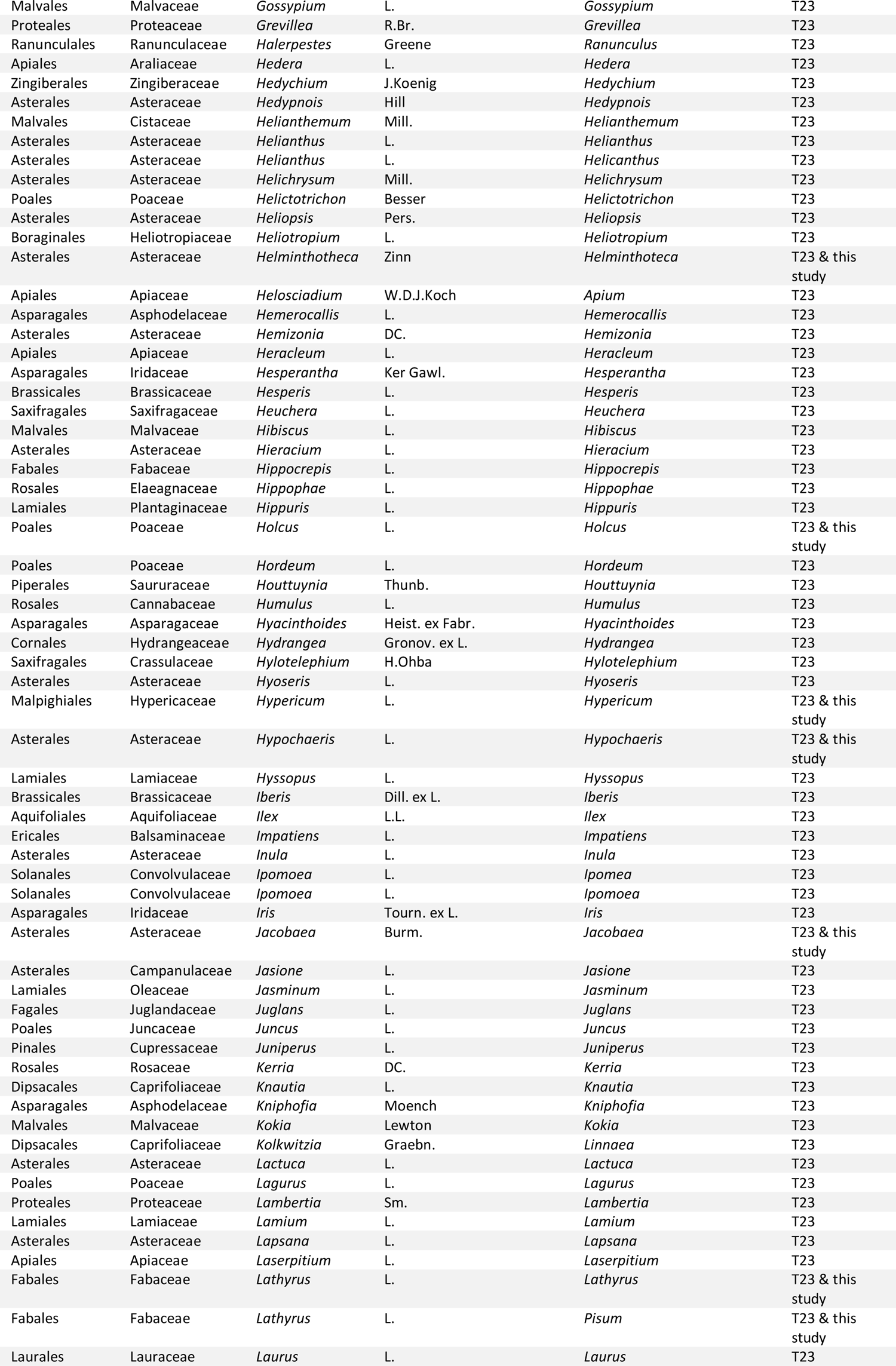
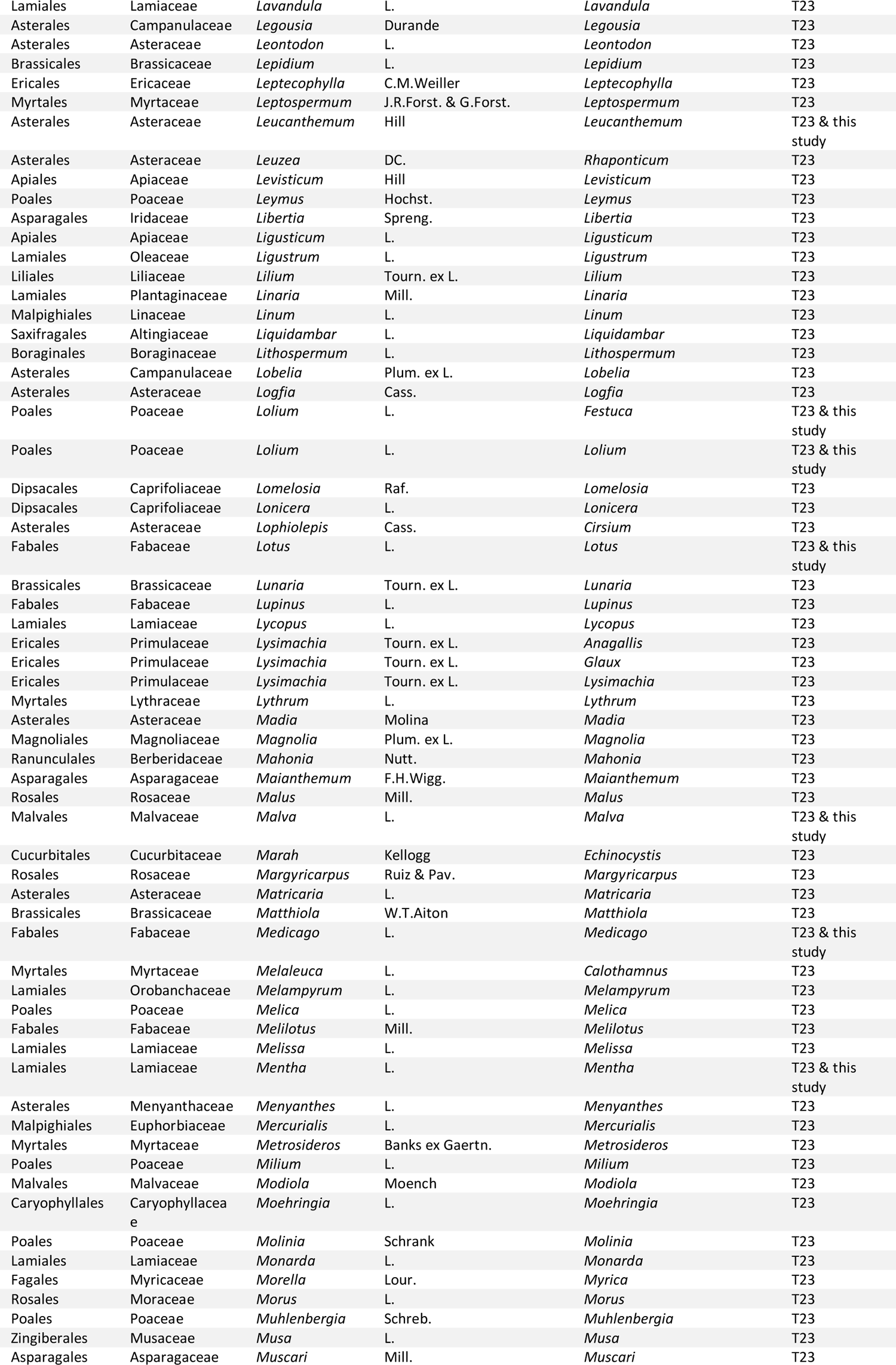
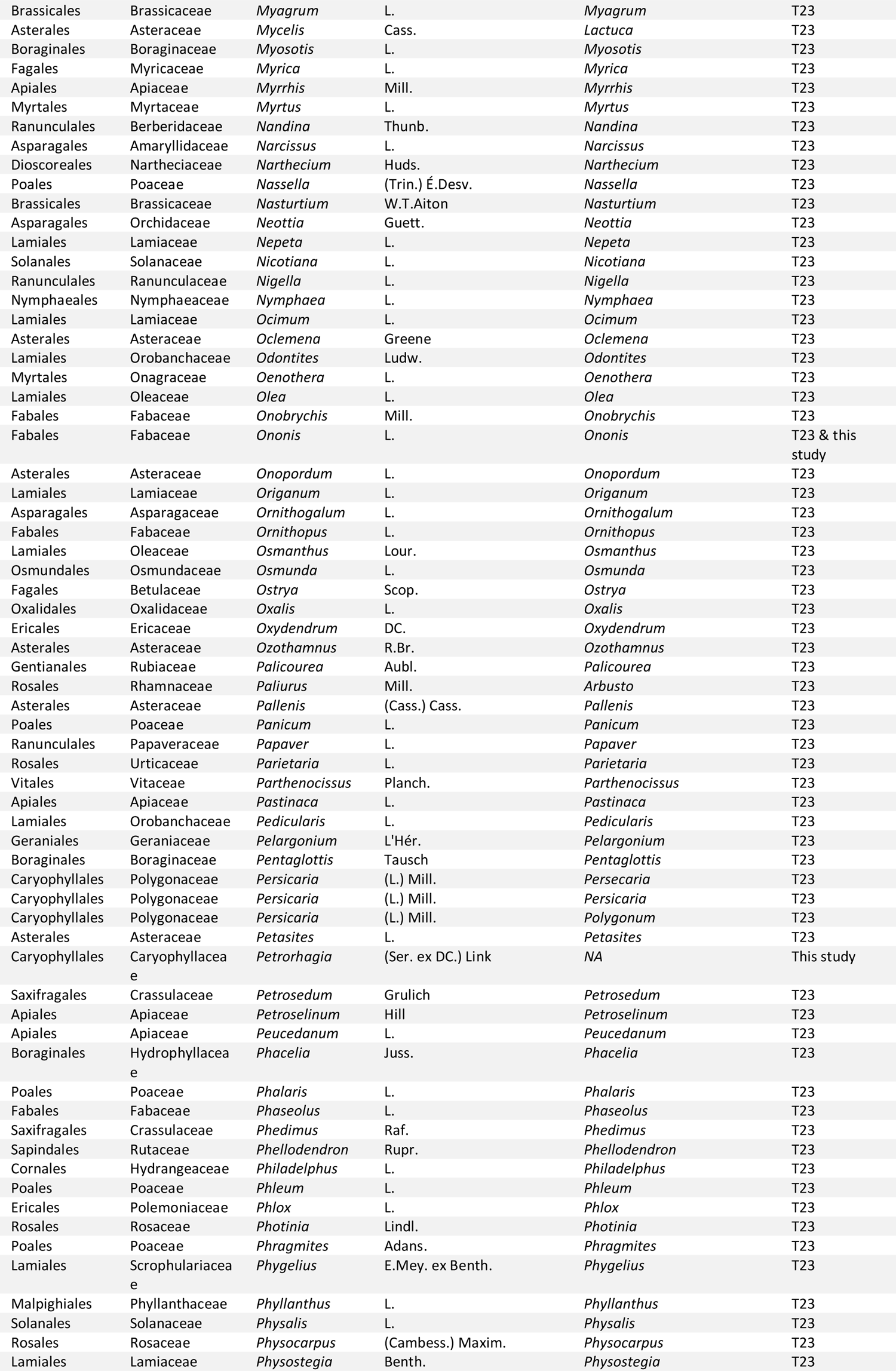
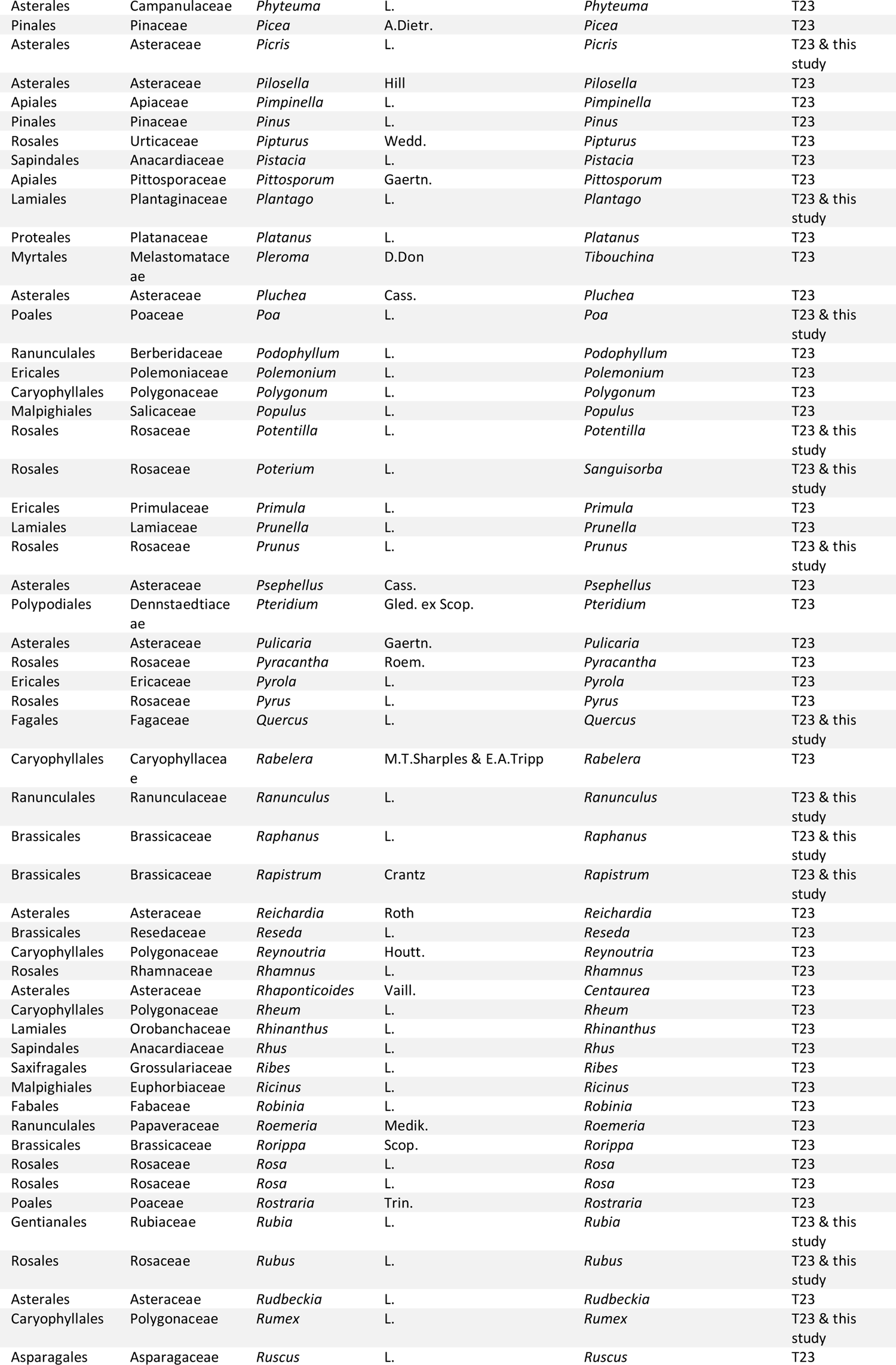
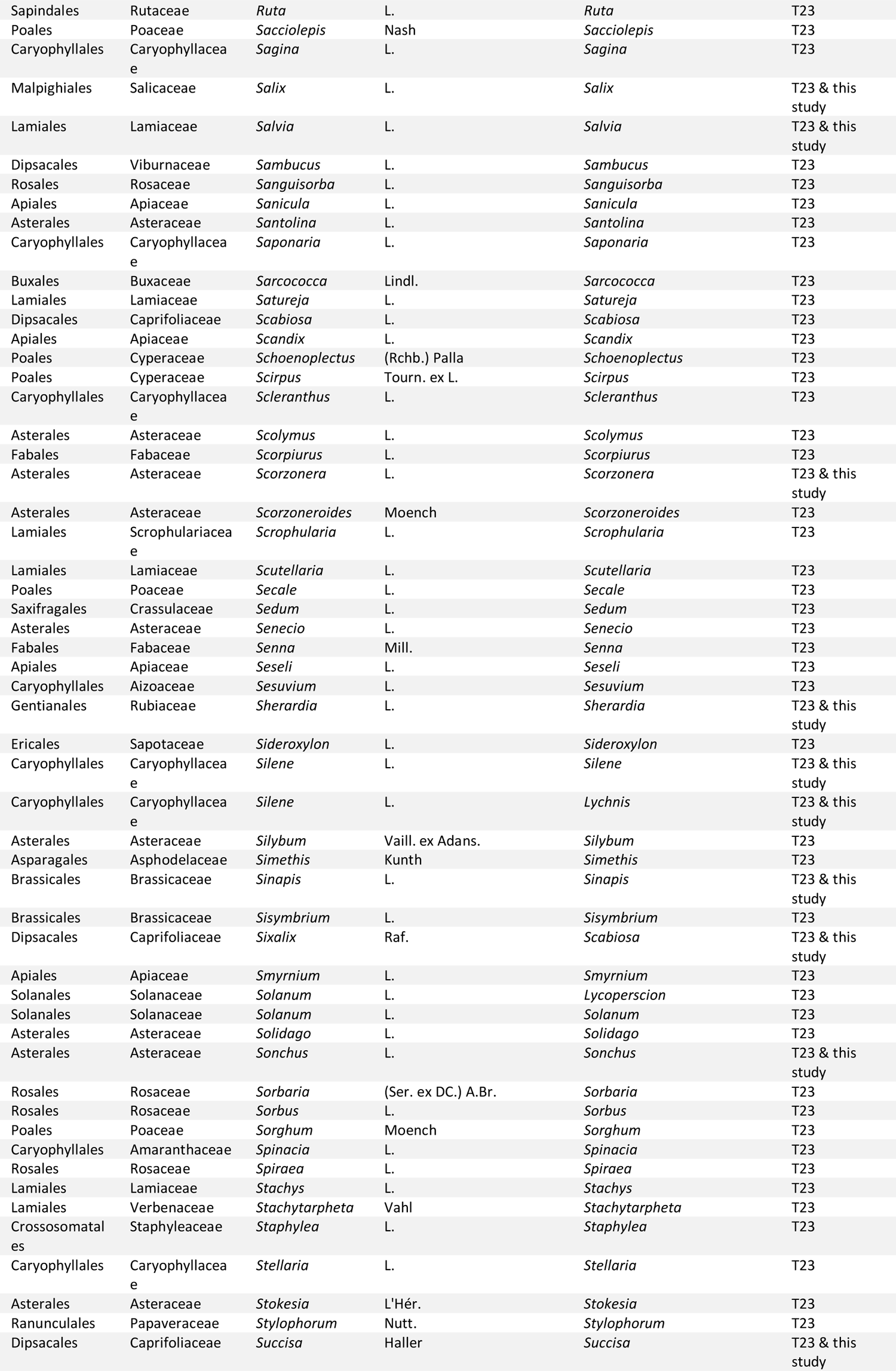
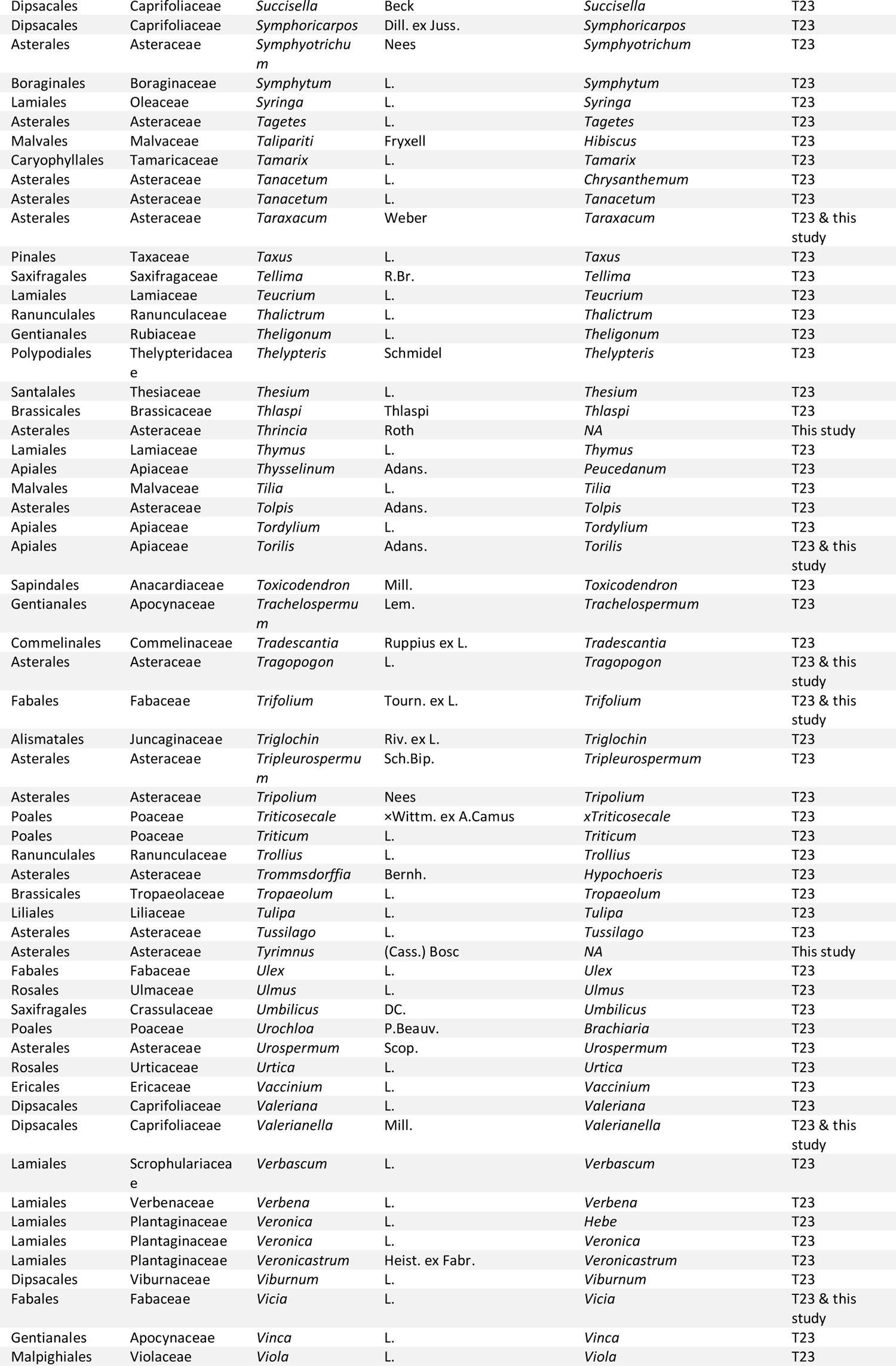
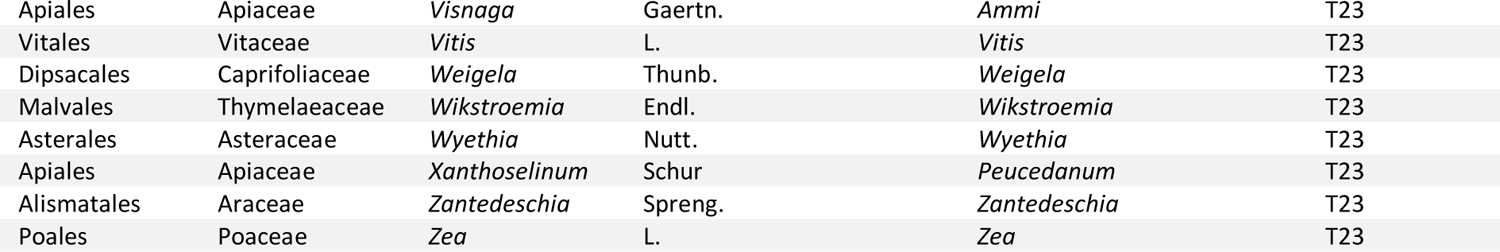
Plant genera listed in Thompson et al. (2023) and in the present study as hosts of *P. spumarius*. Taxonomy follows the currently accepted names in GBIF (GBIF.org, 2023) after check with ‘get_gbif_taxonomy » function in R (Schneider, 2018). As several species initially described in the same genus may be distributed in different genera, we retrieved the corrected species names in Table S8.2 to perform the taxonomic revision of genera. “T23“ in the “Source” column stands for “Already reported (Thompson et al., 2023)”.

**Table S10.2.**
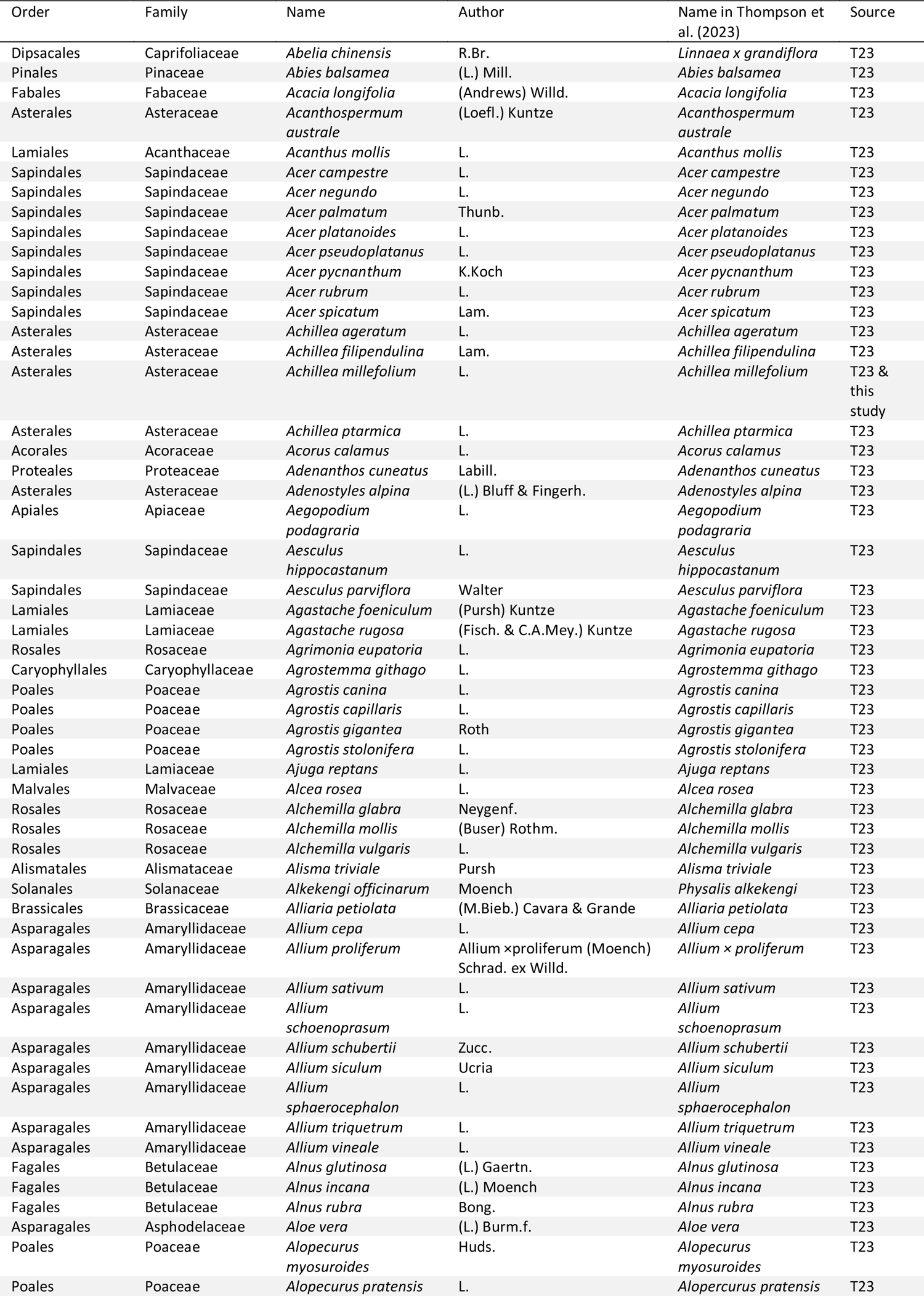
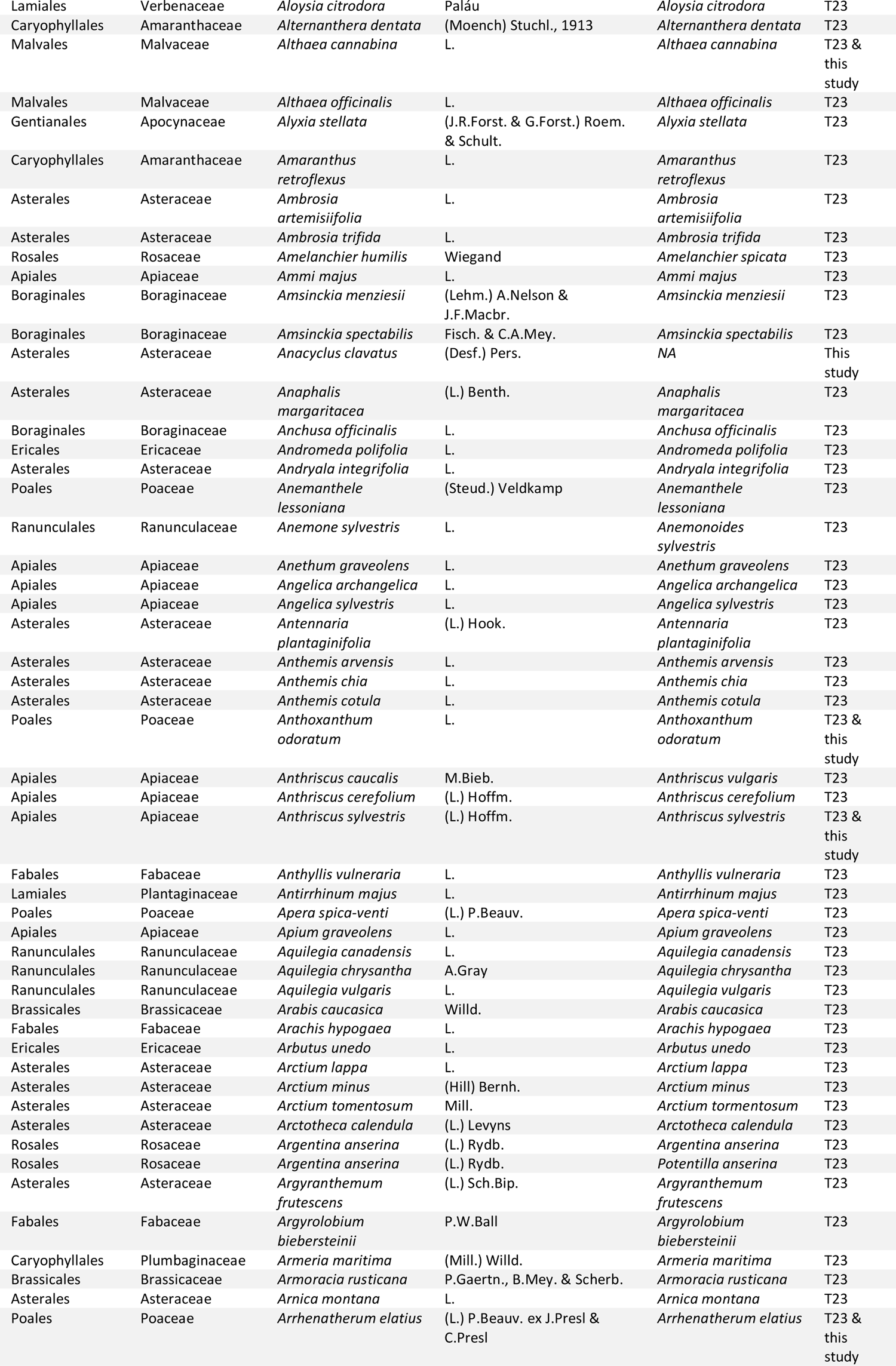
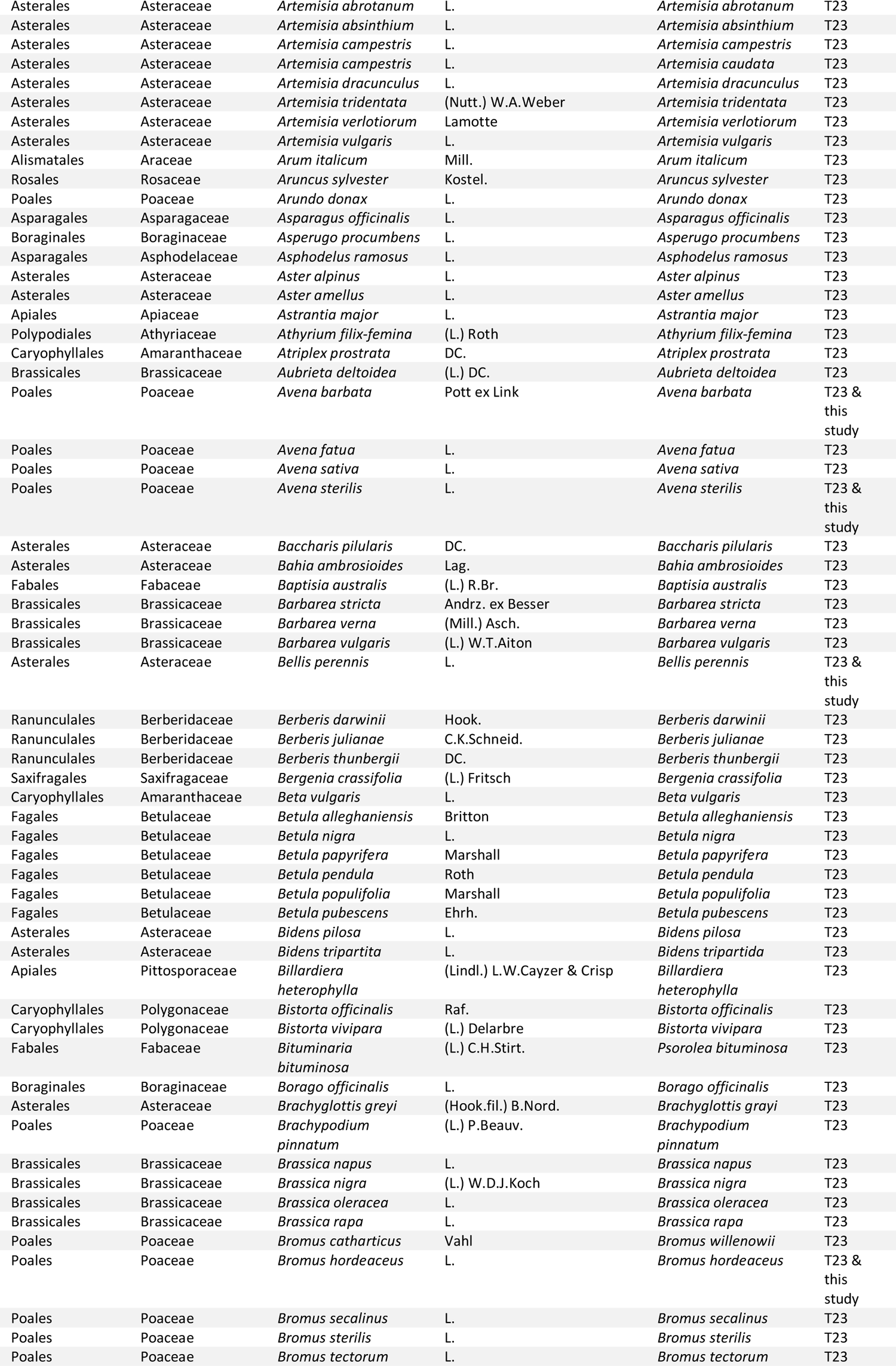
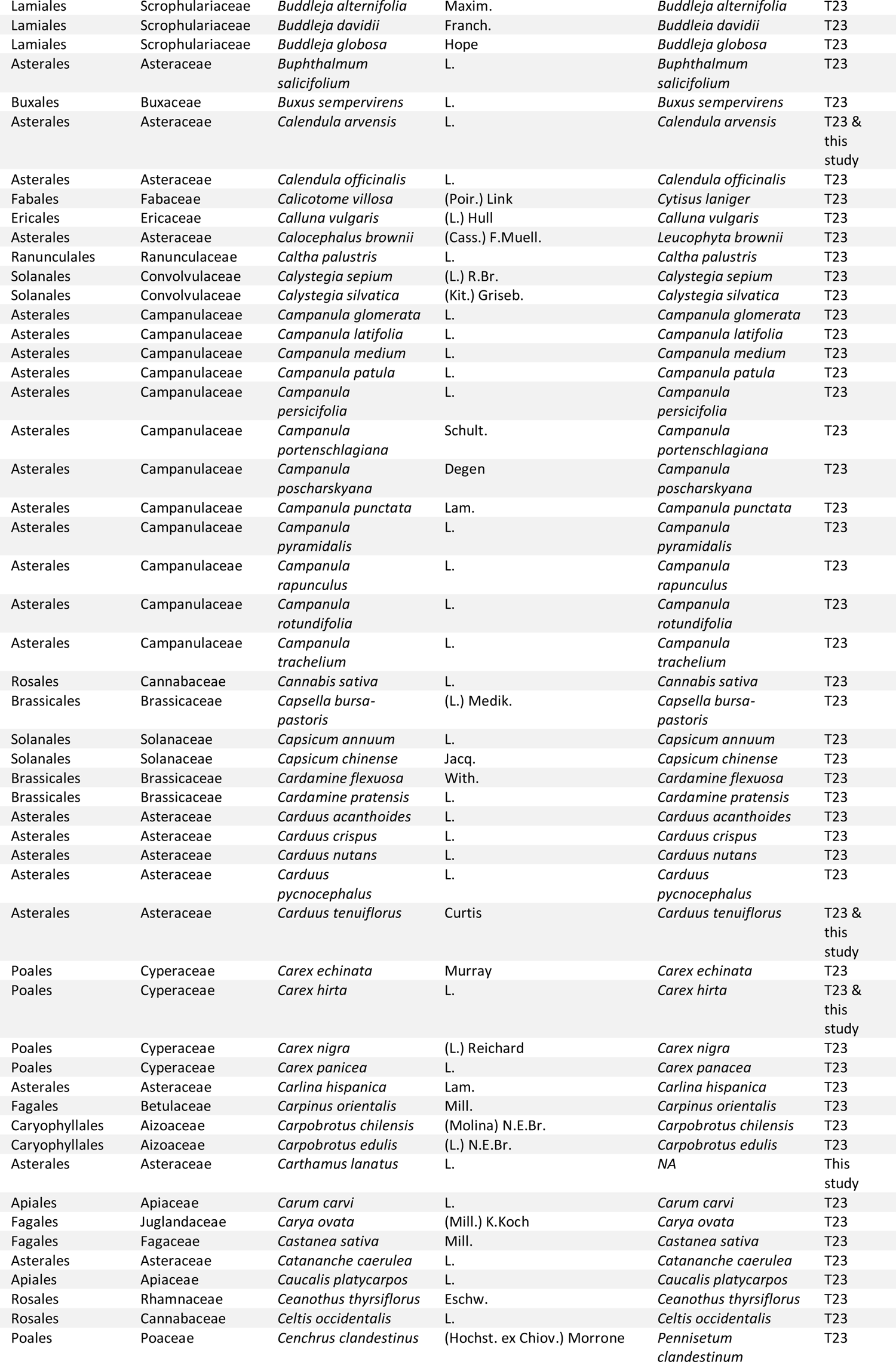
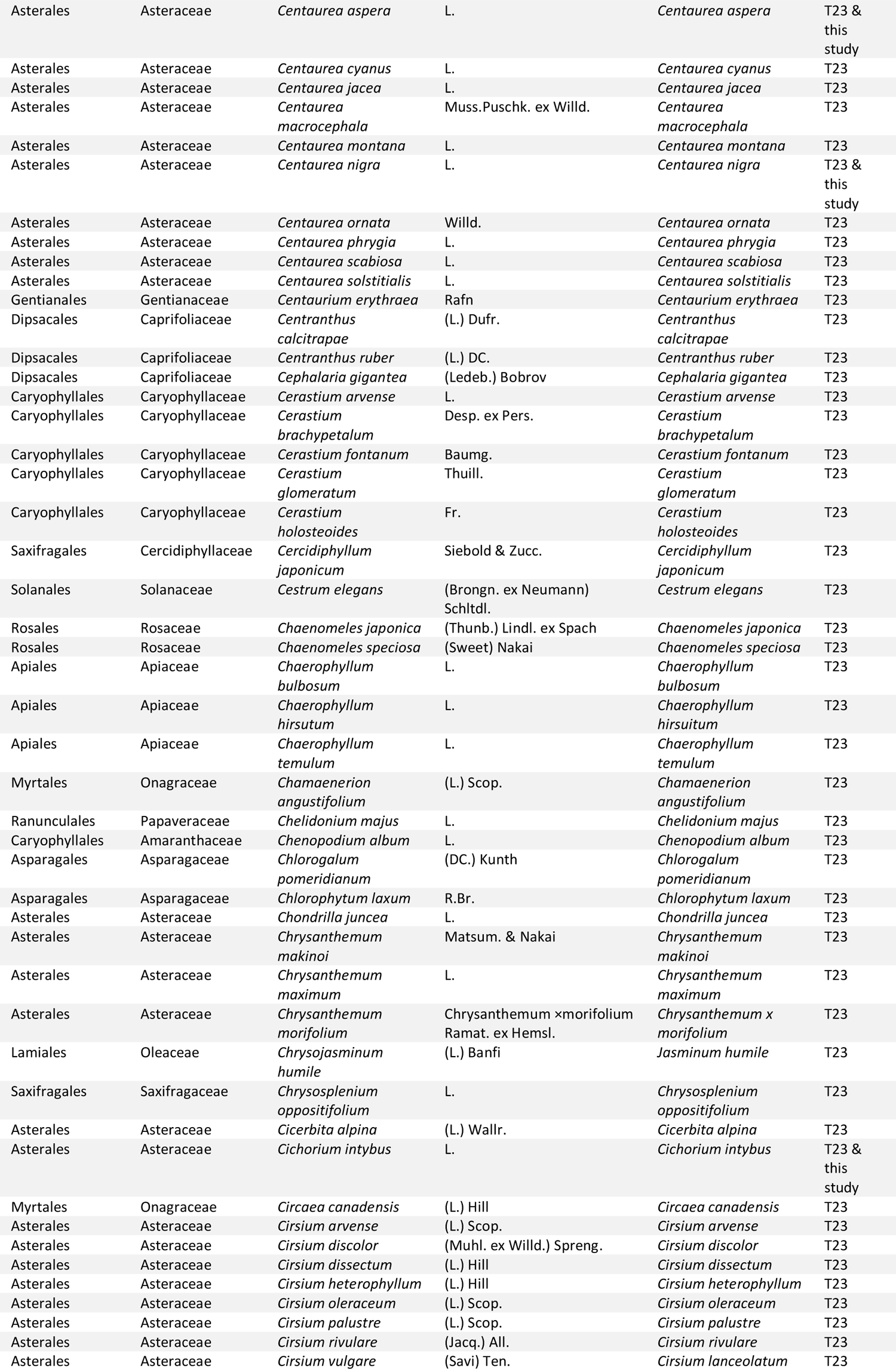
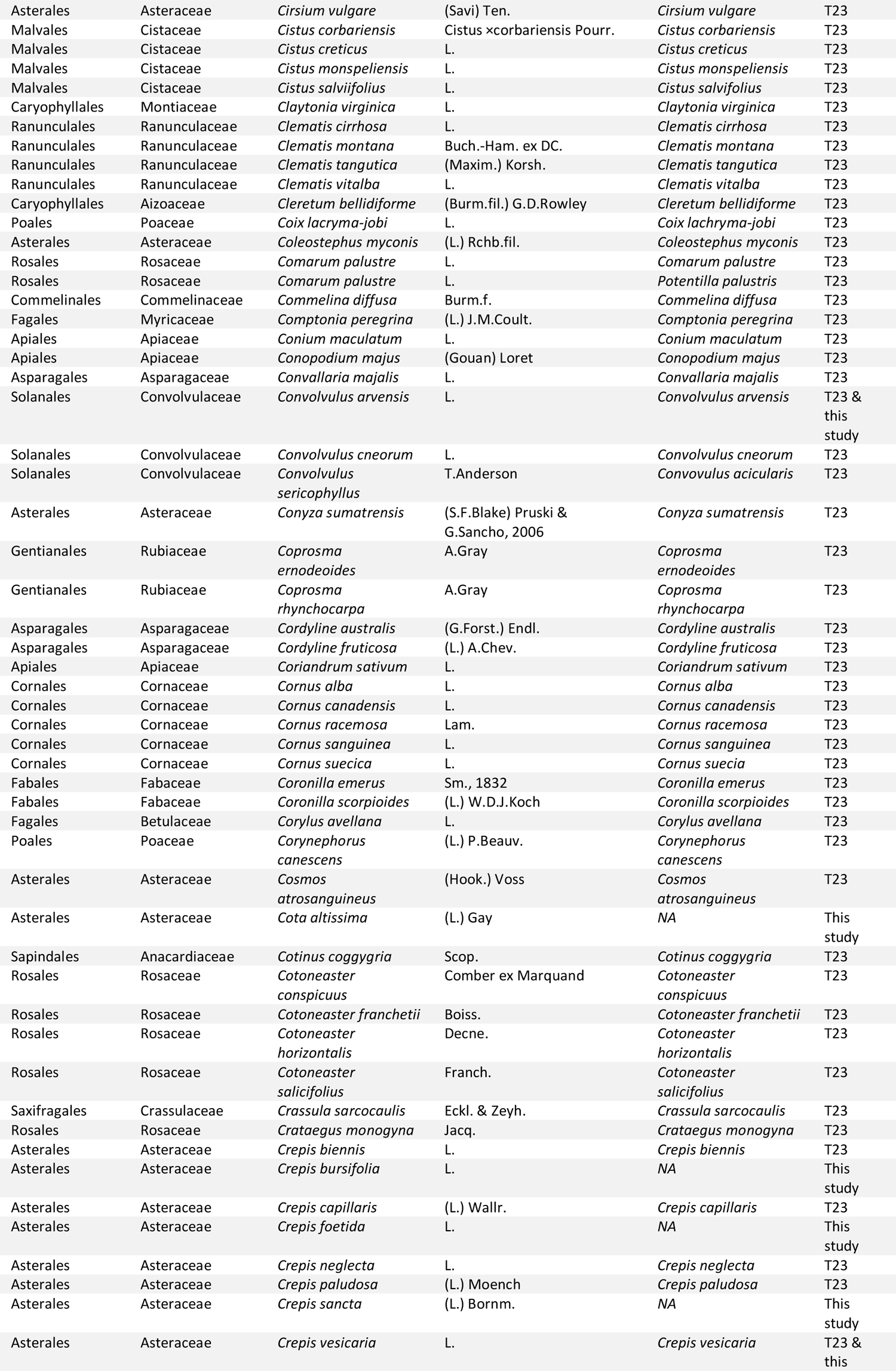
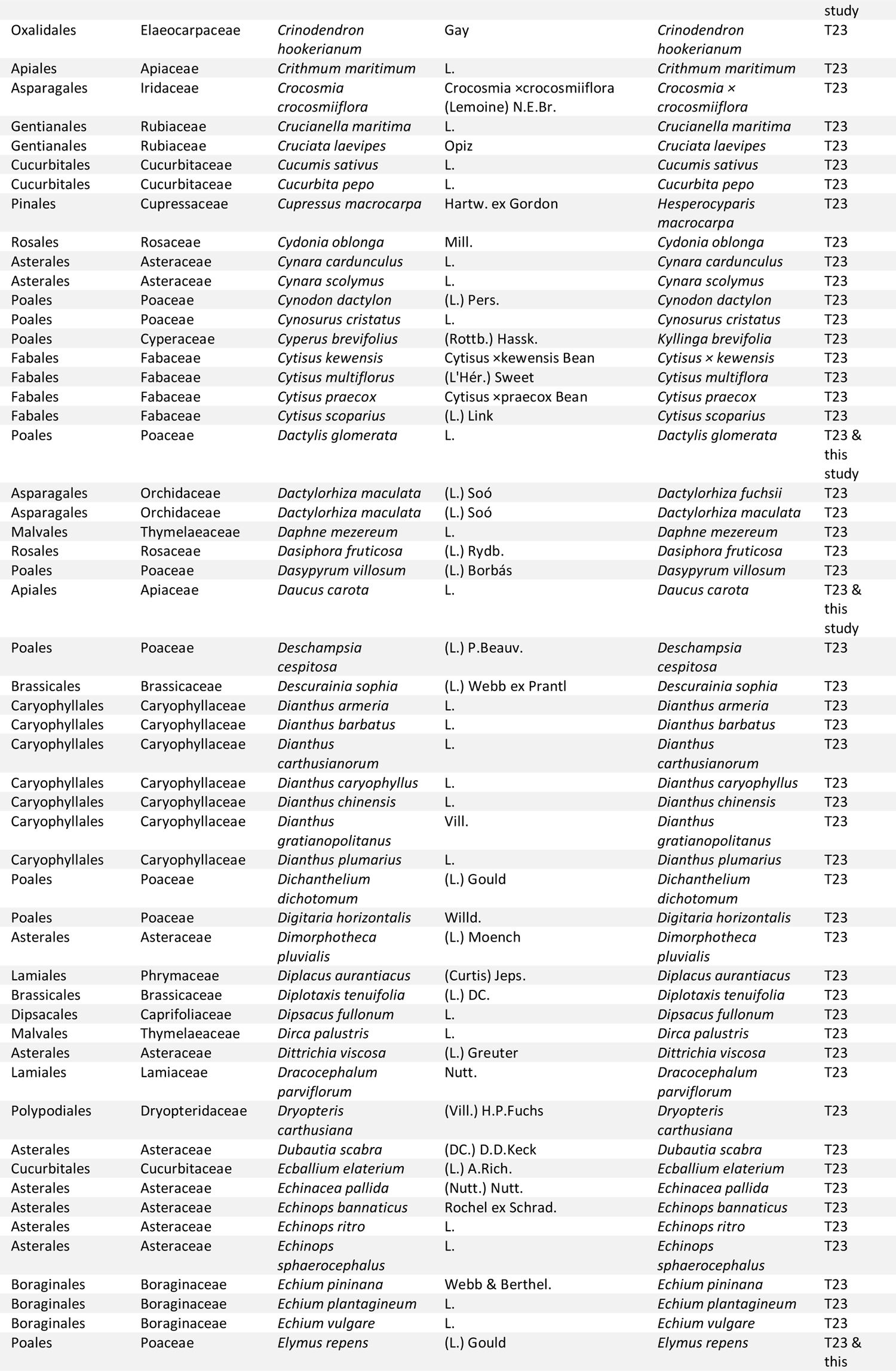
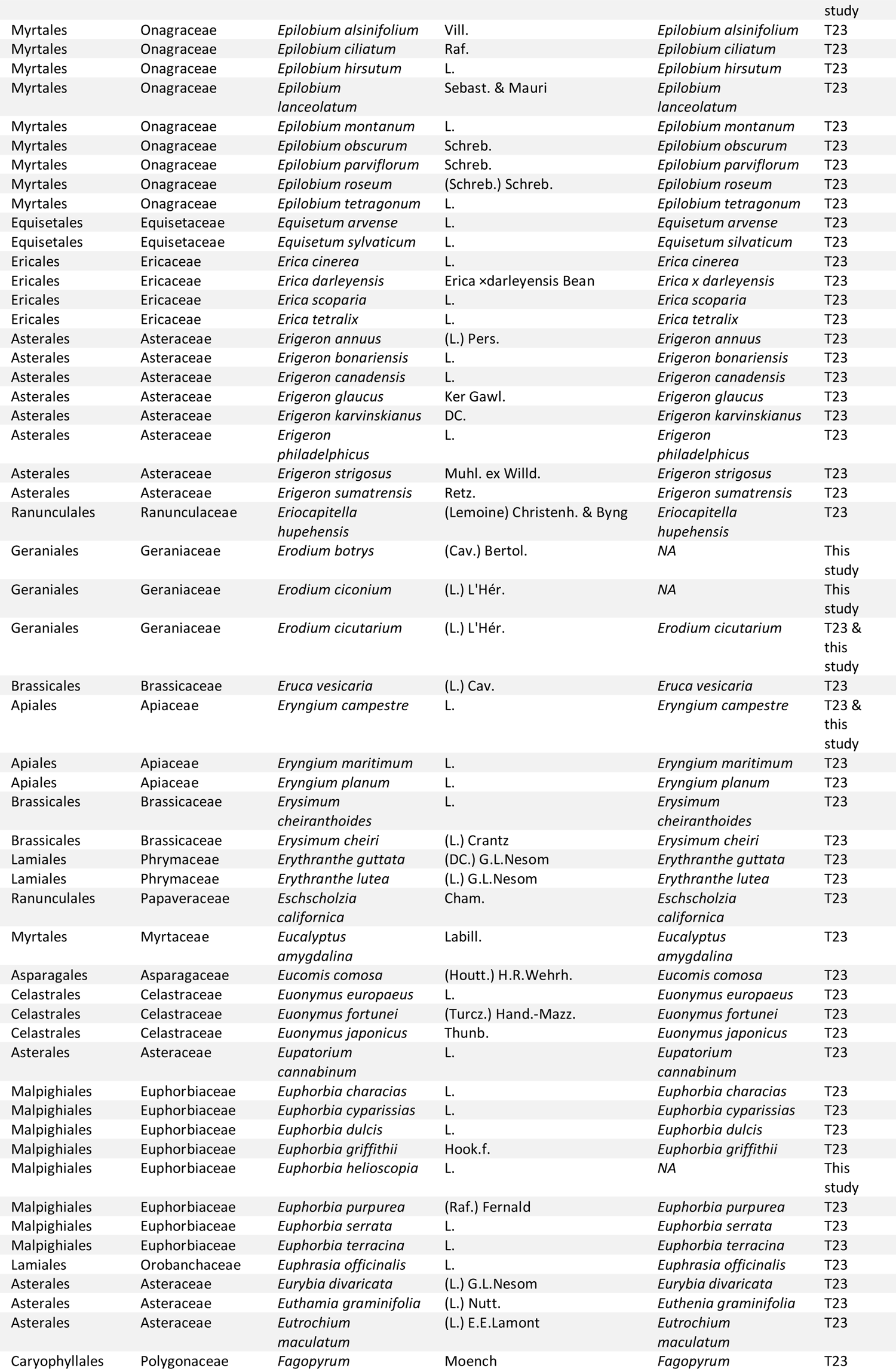
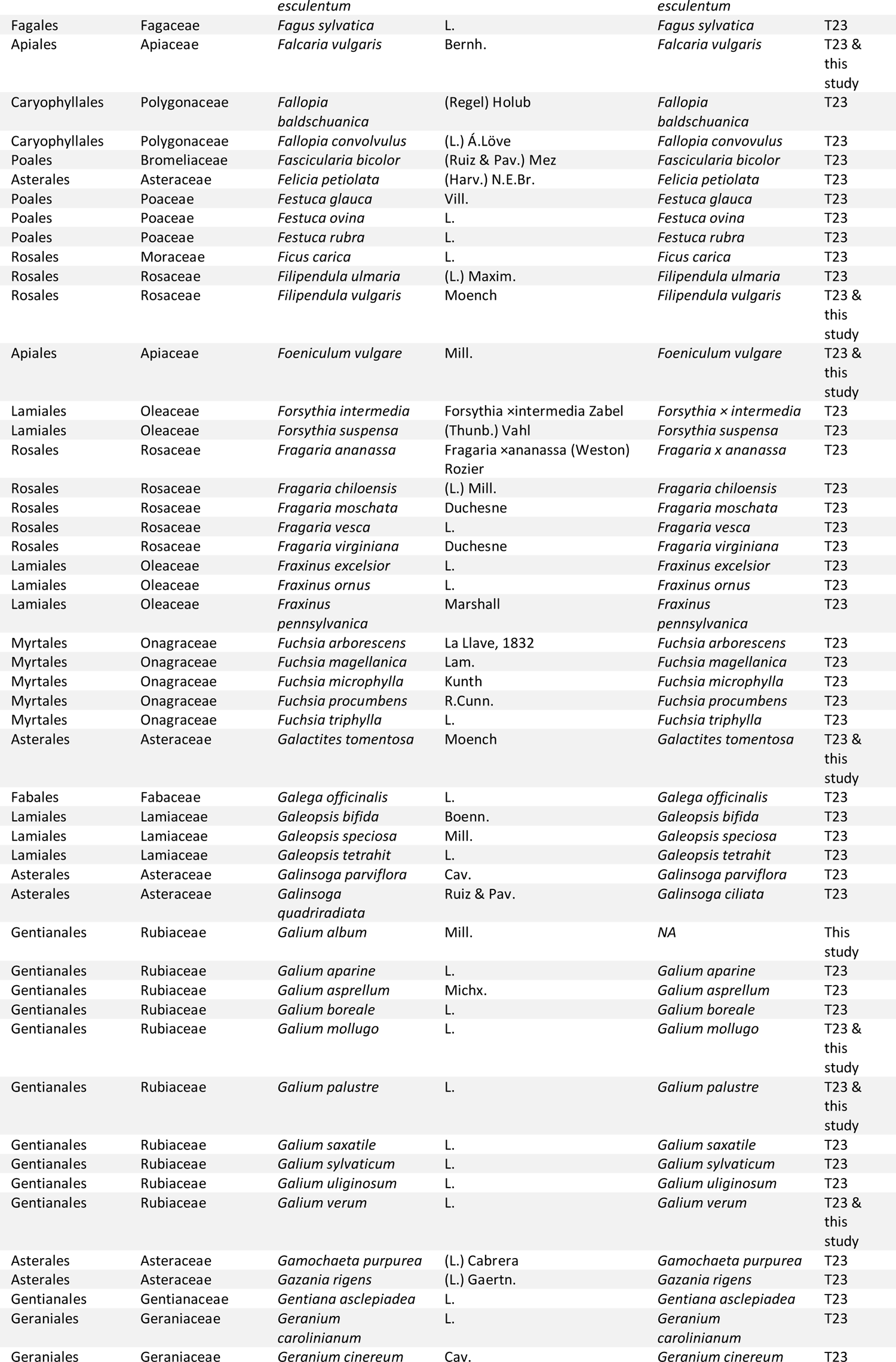
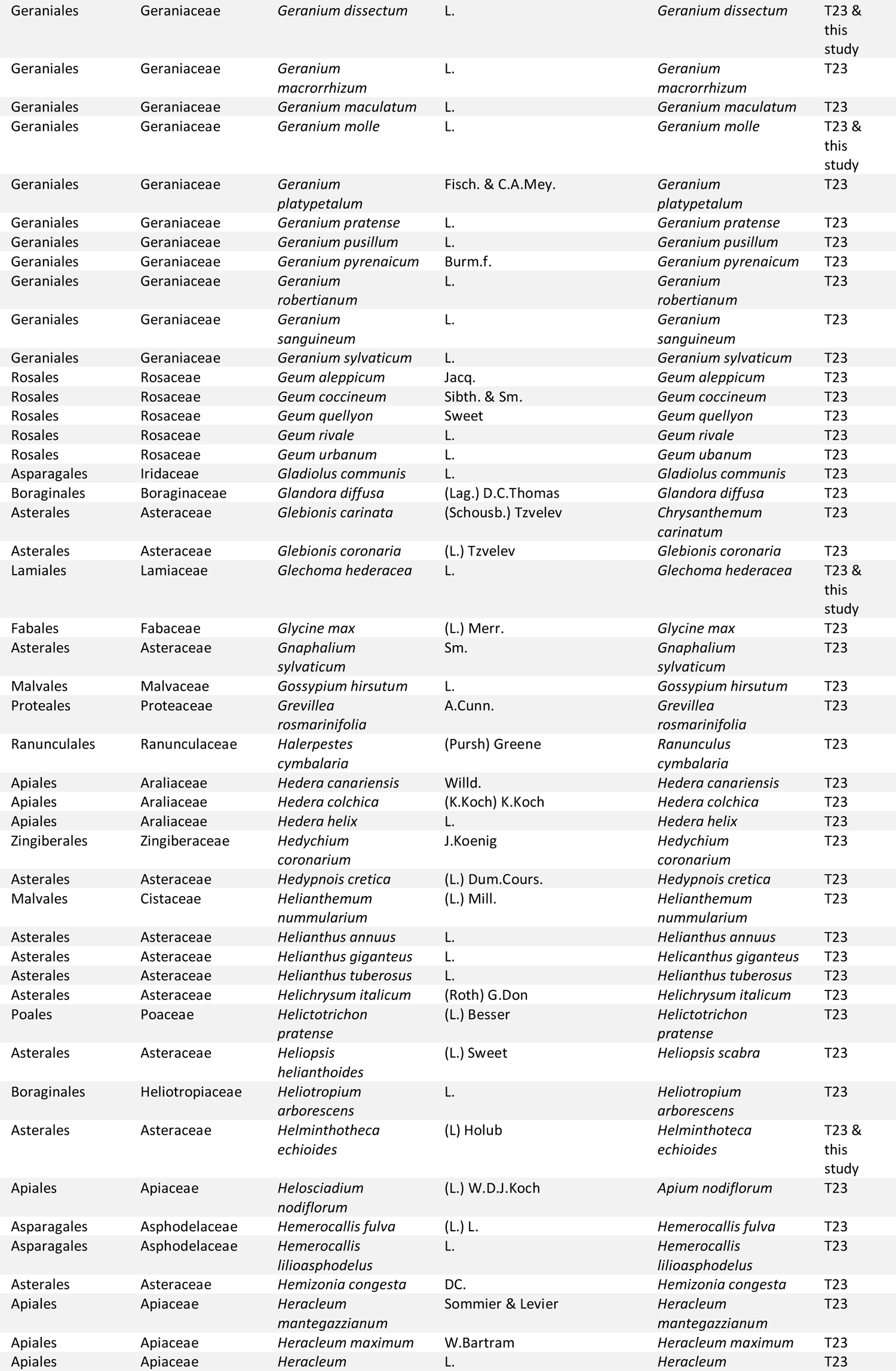
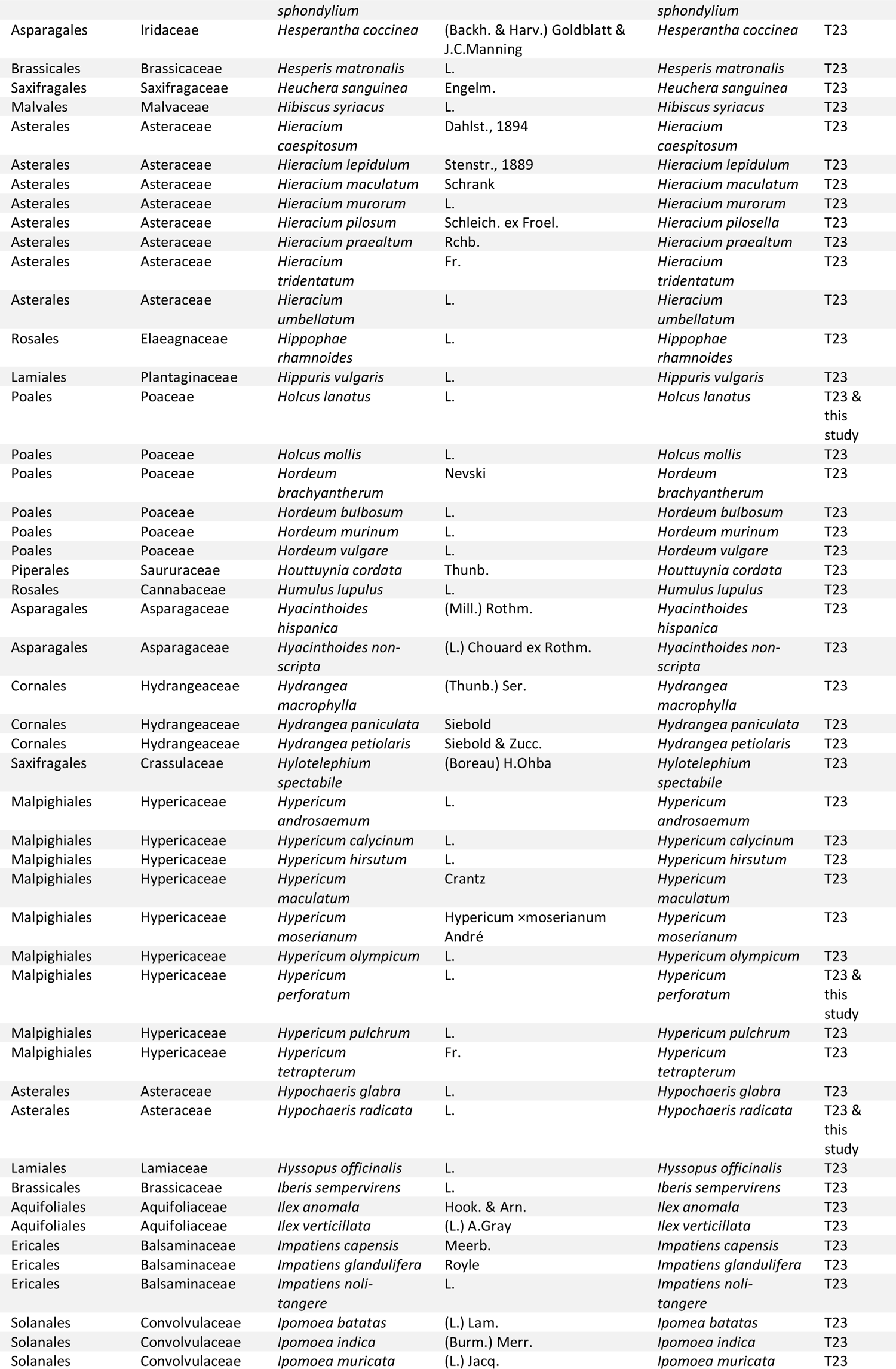
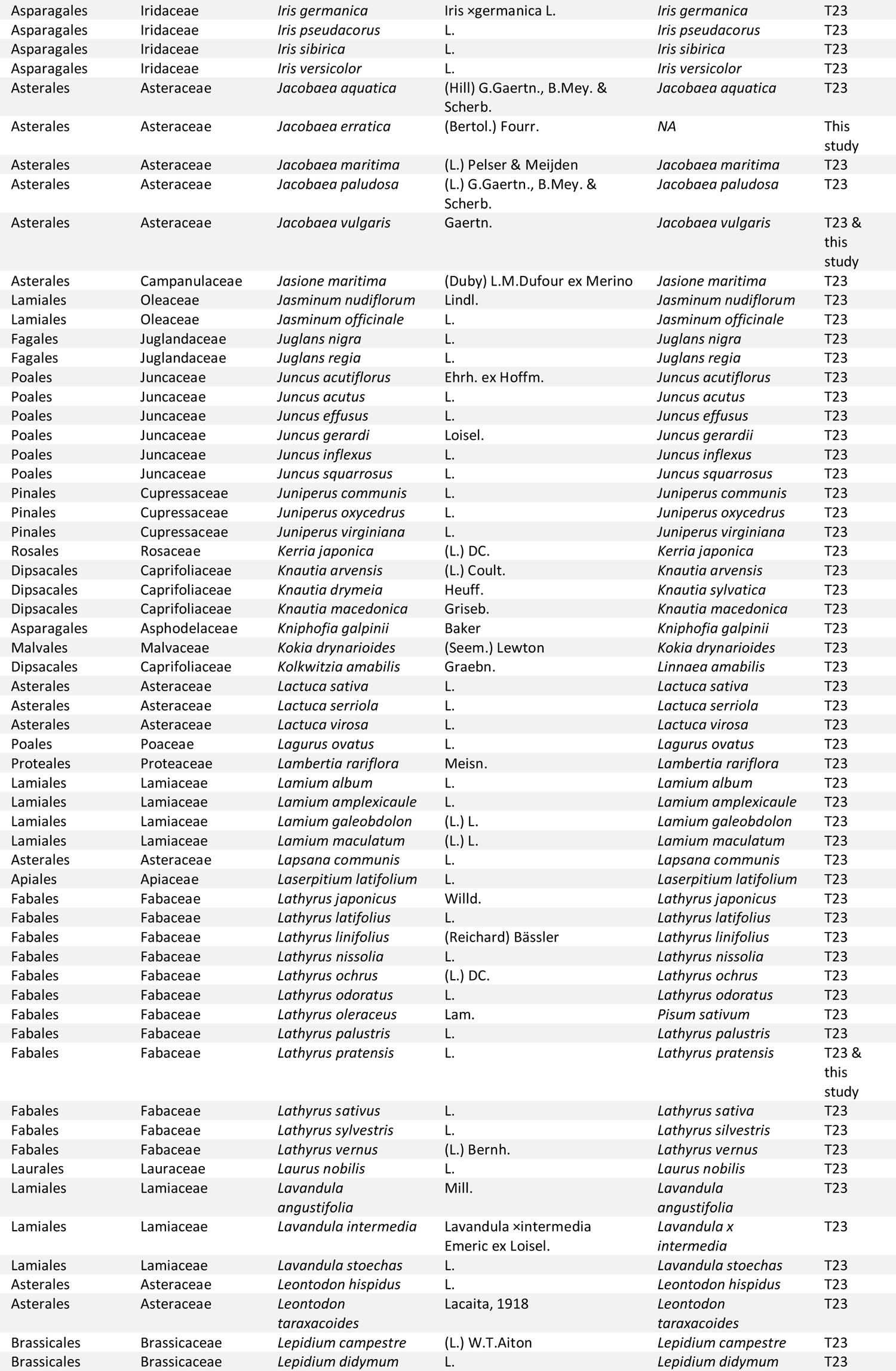
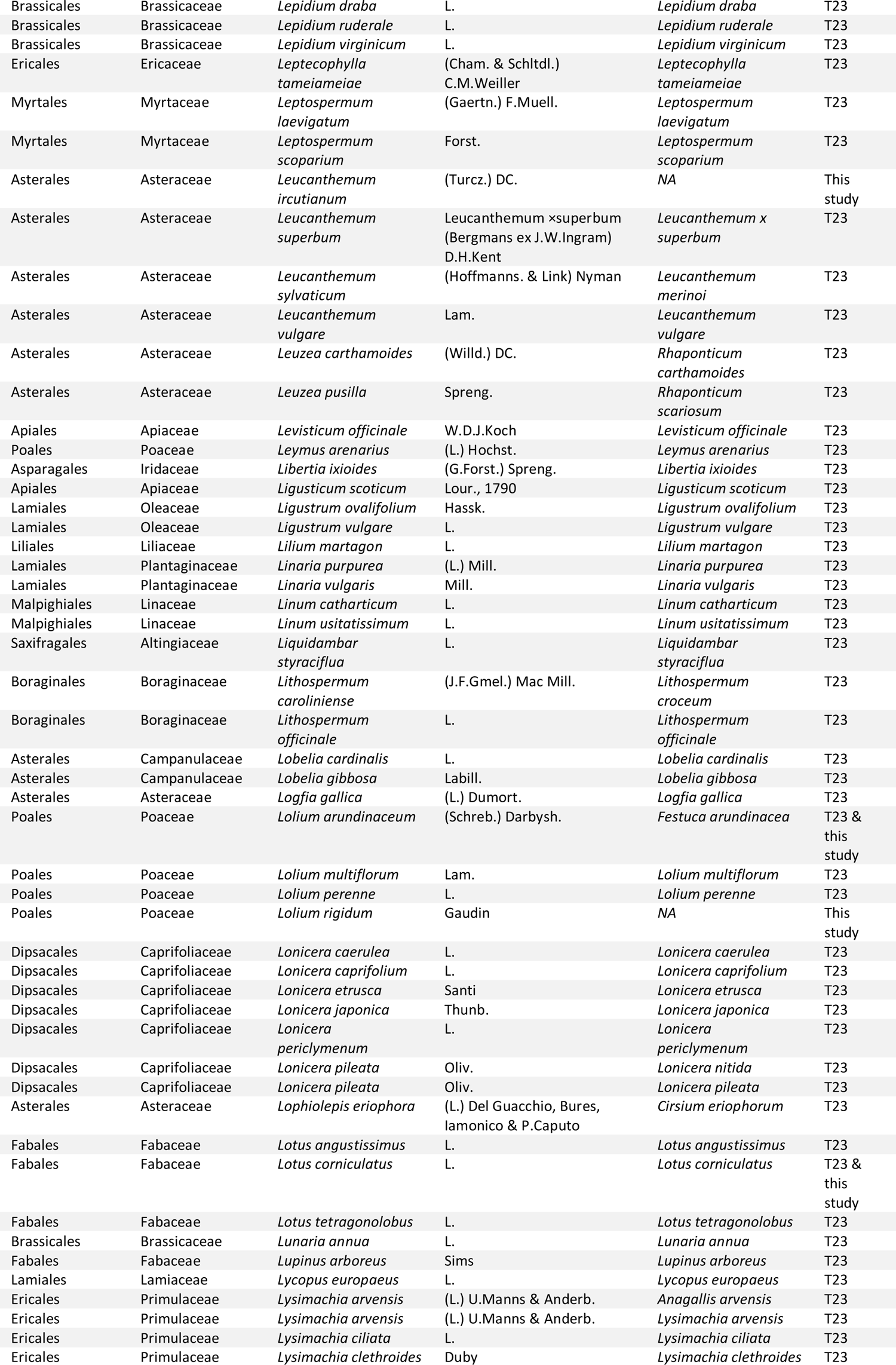
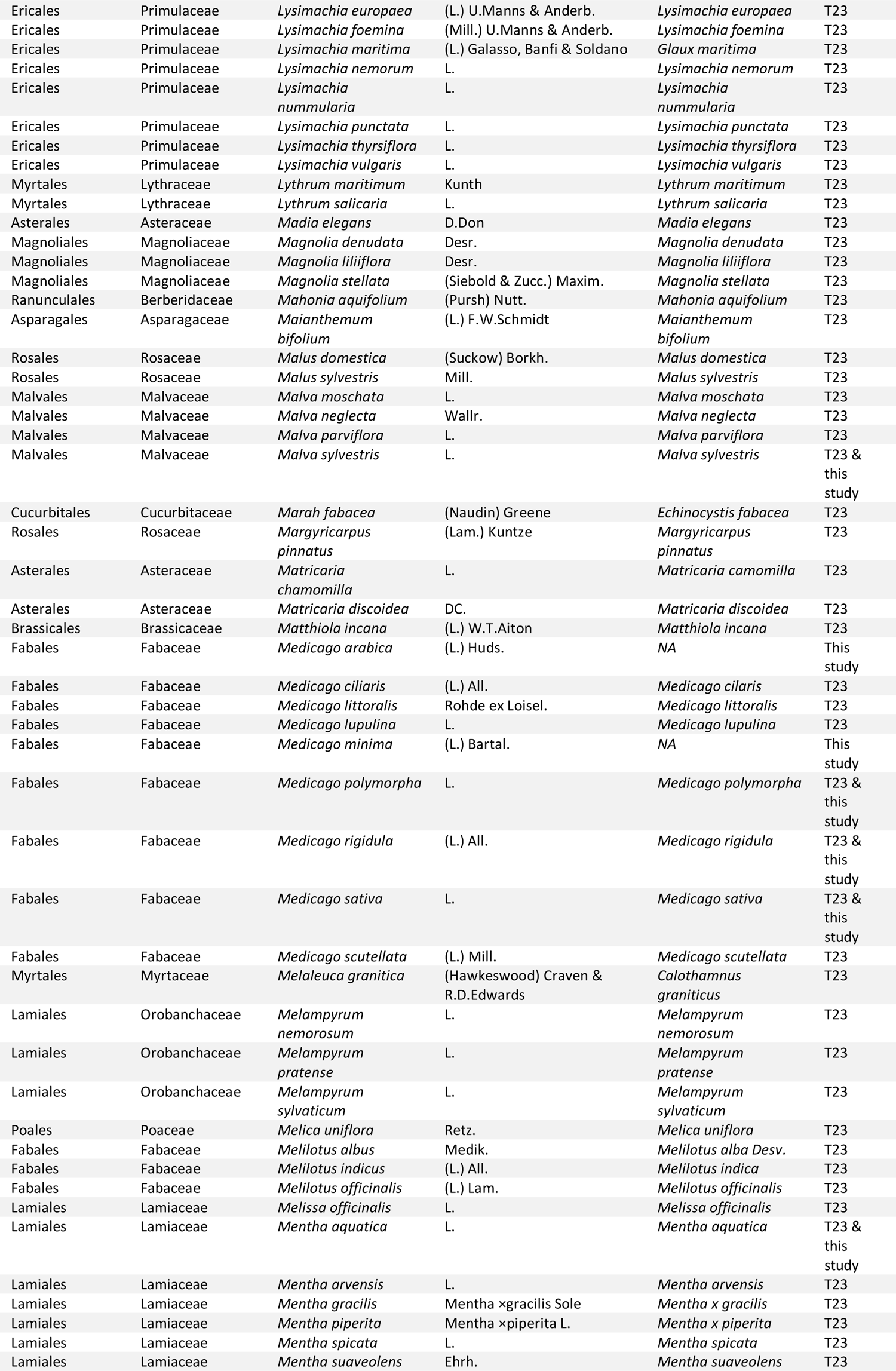
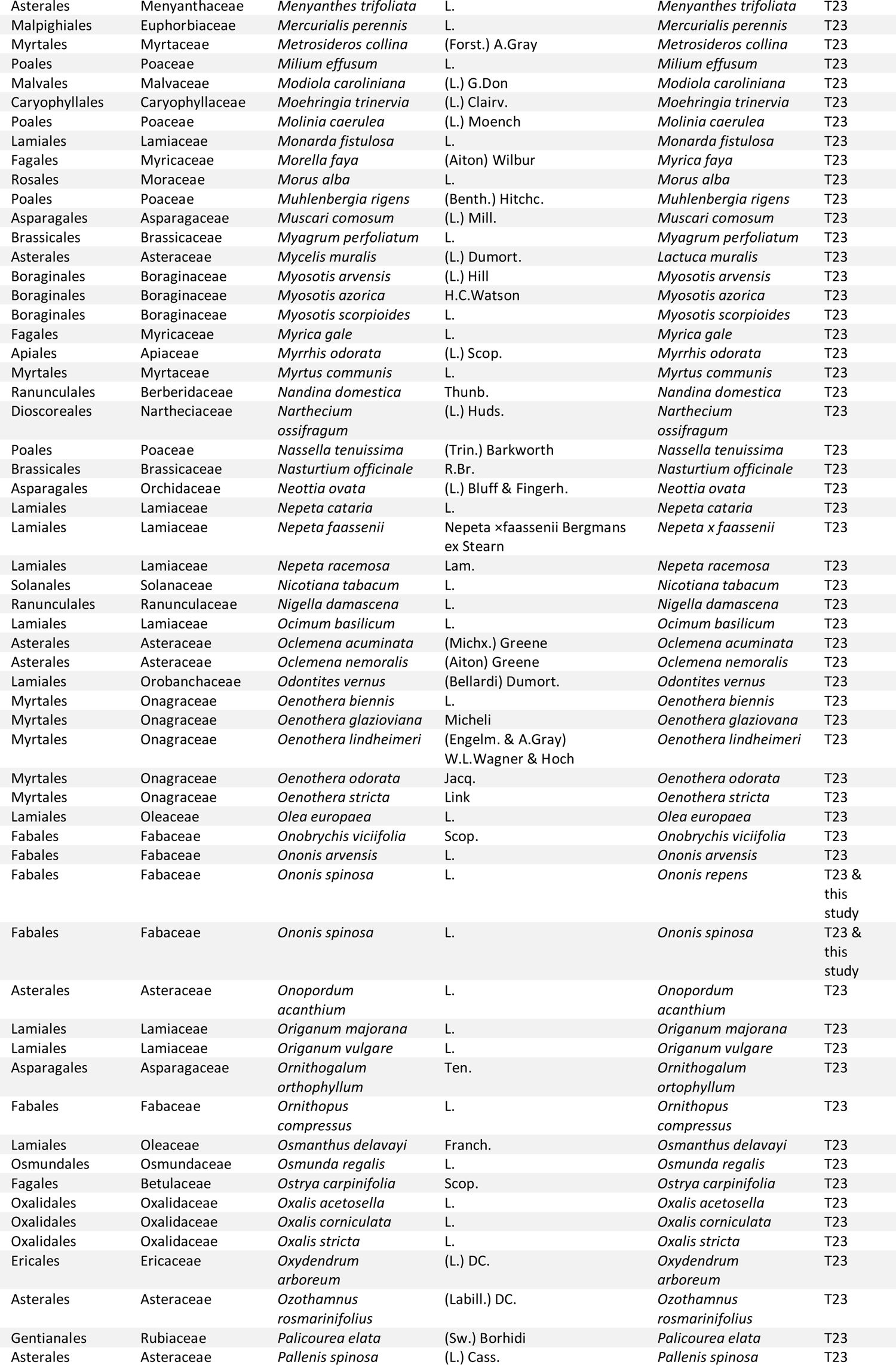
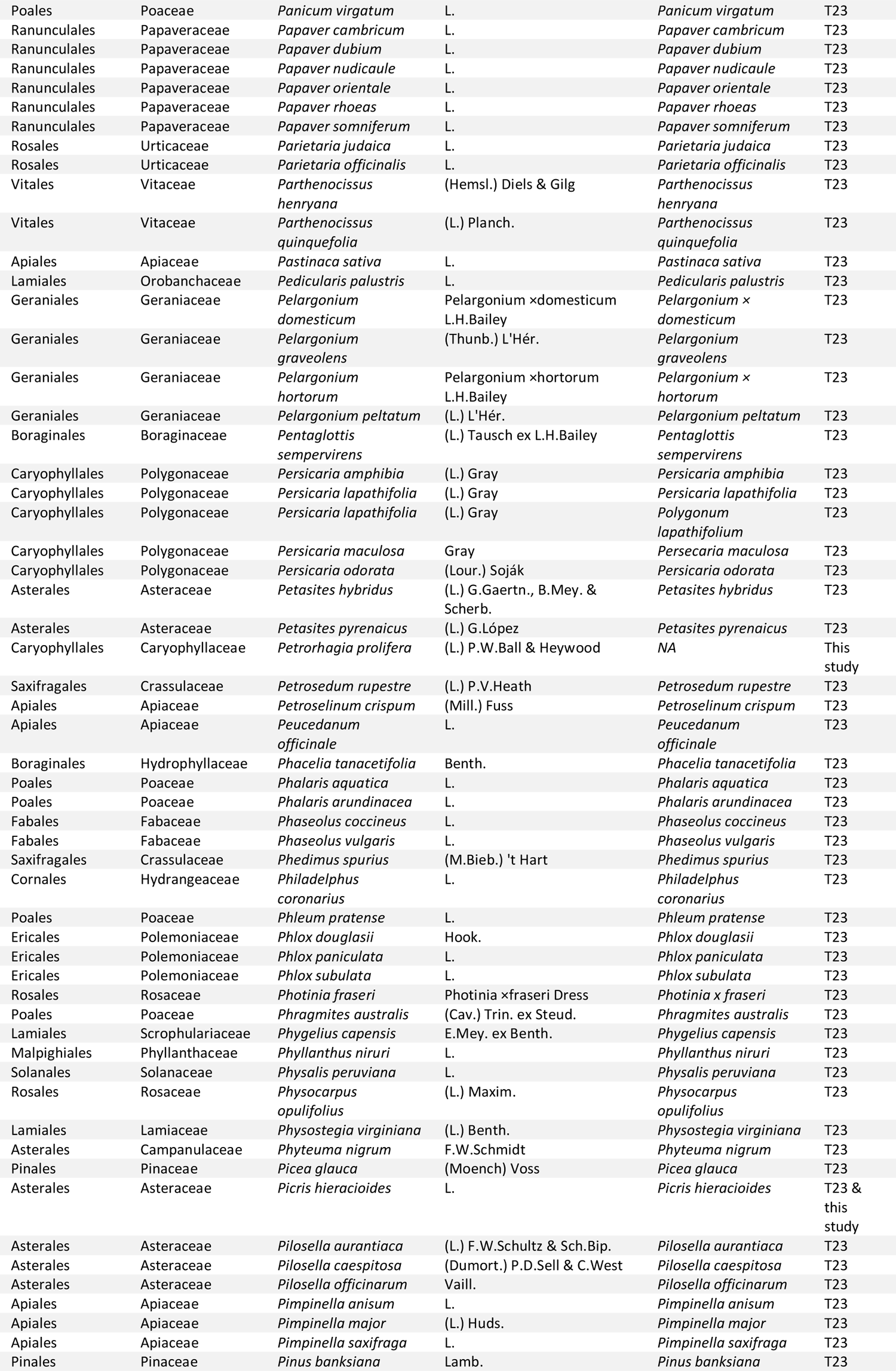
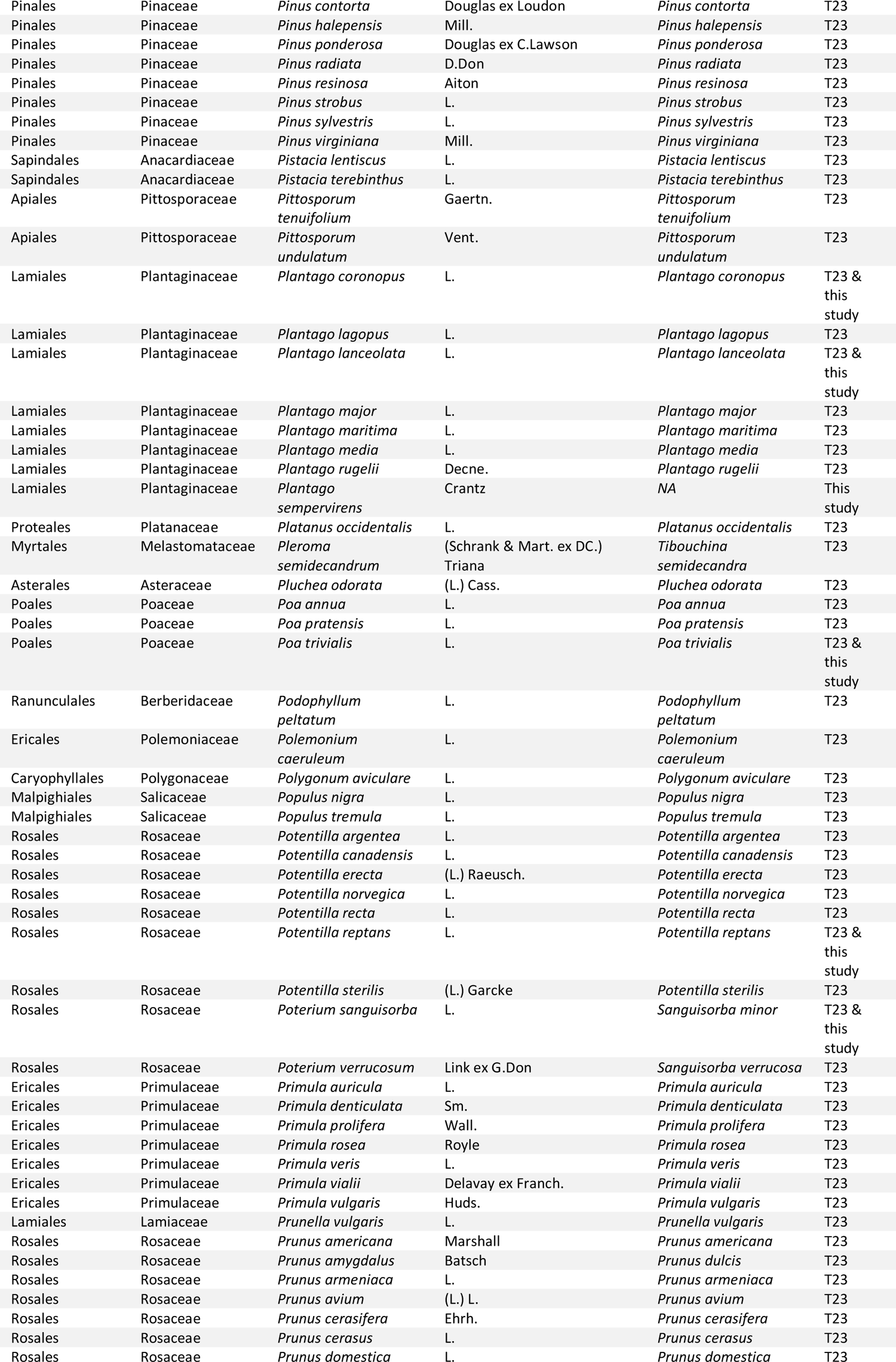
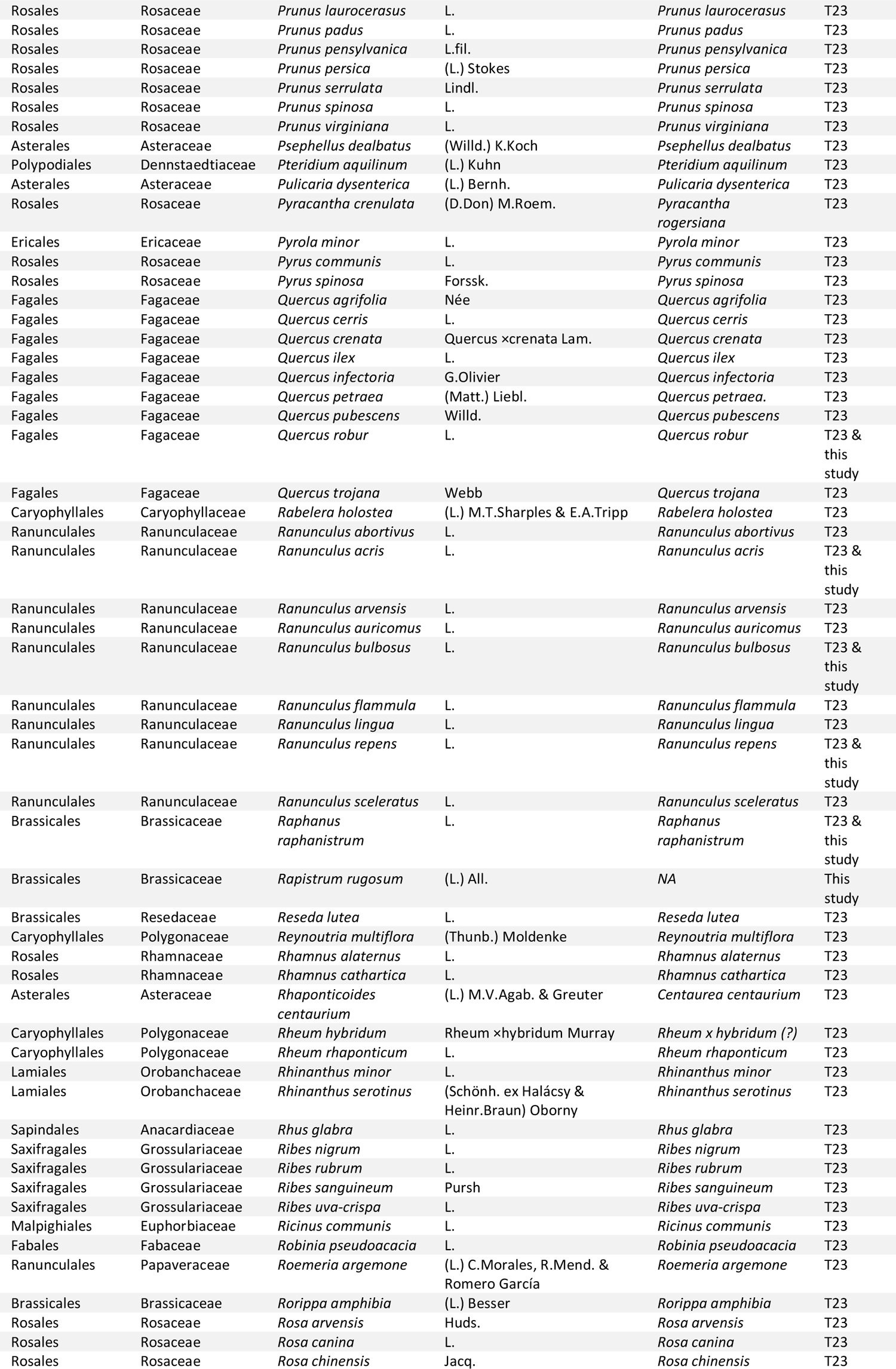
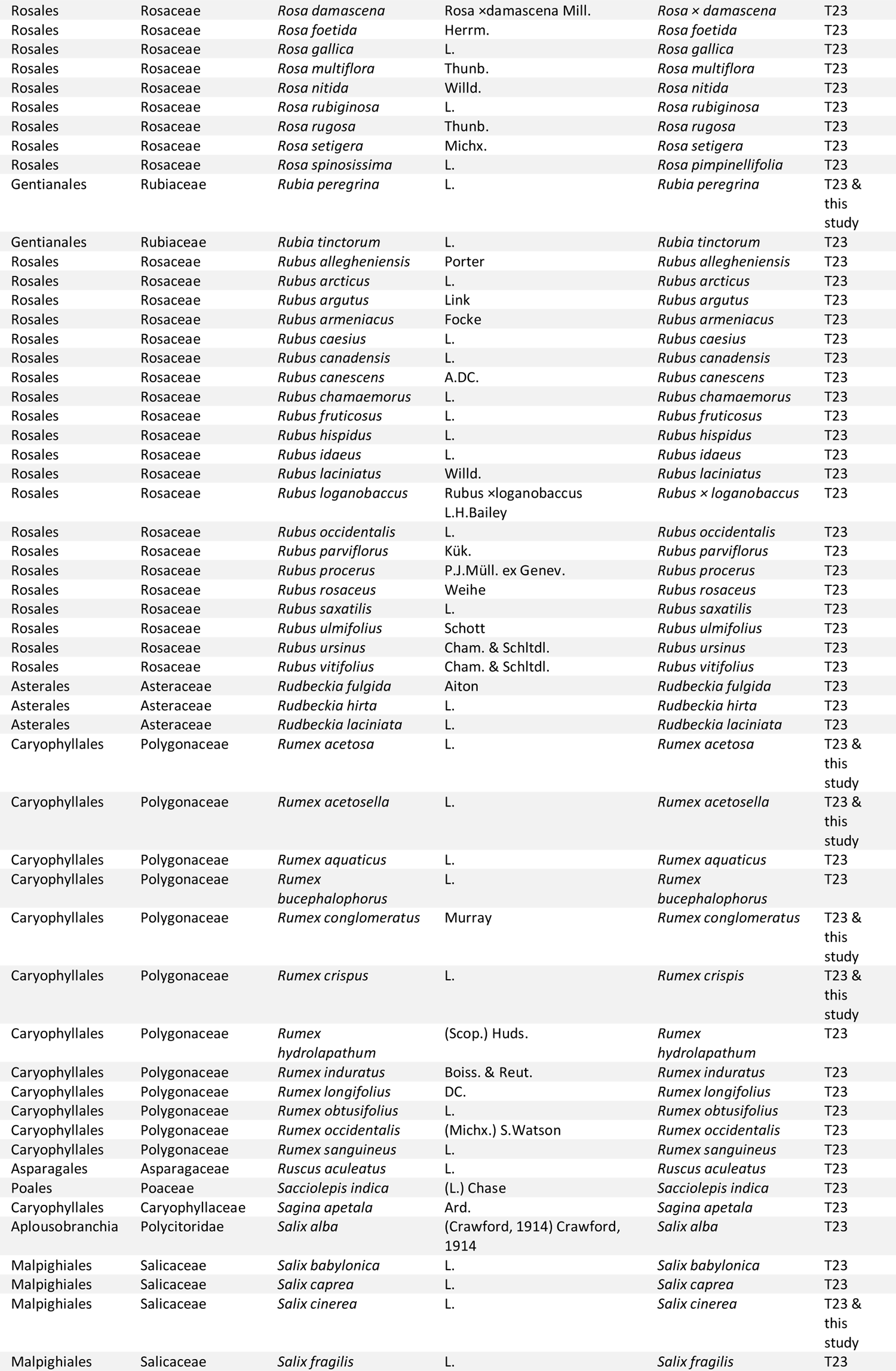
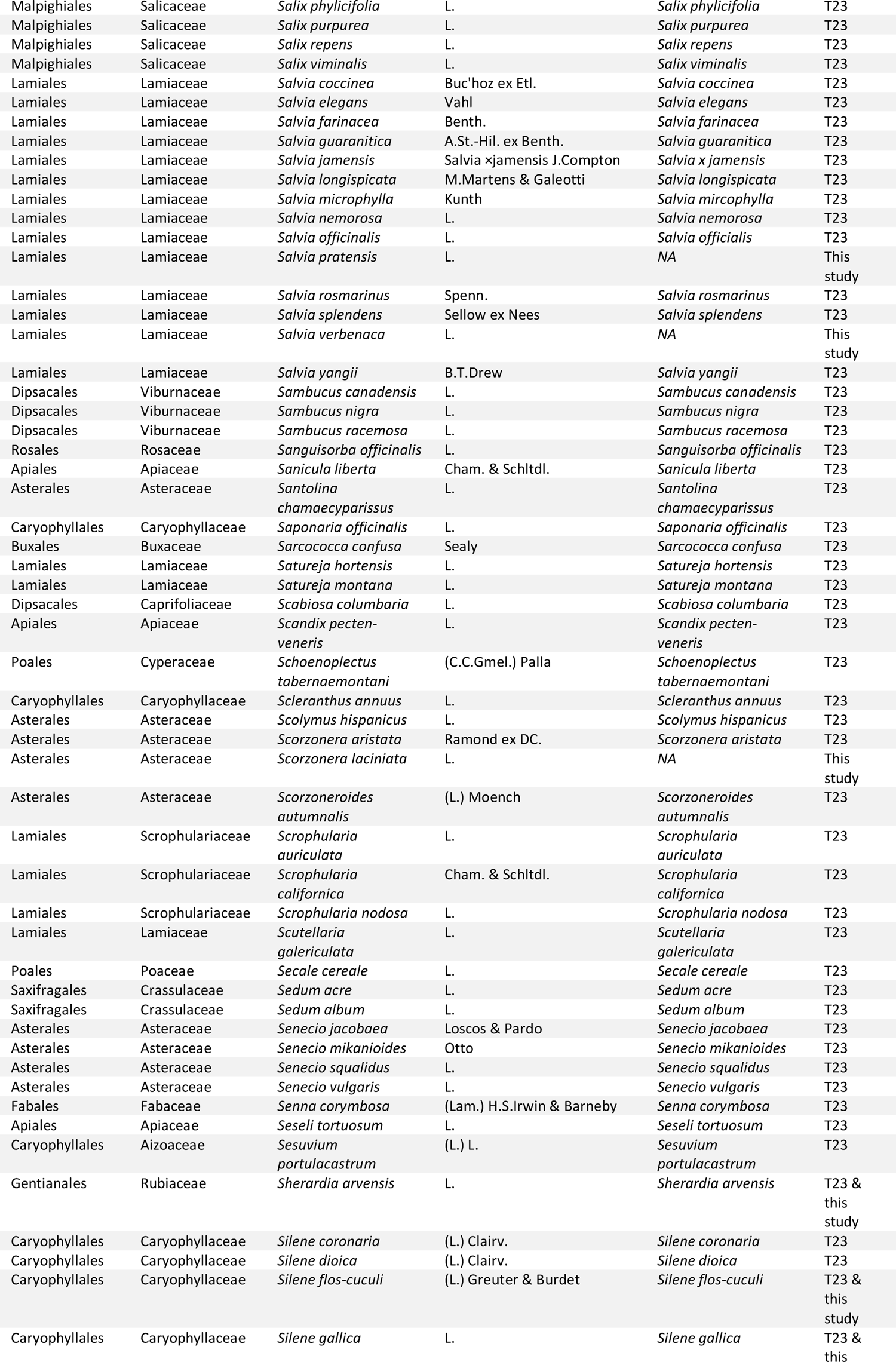
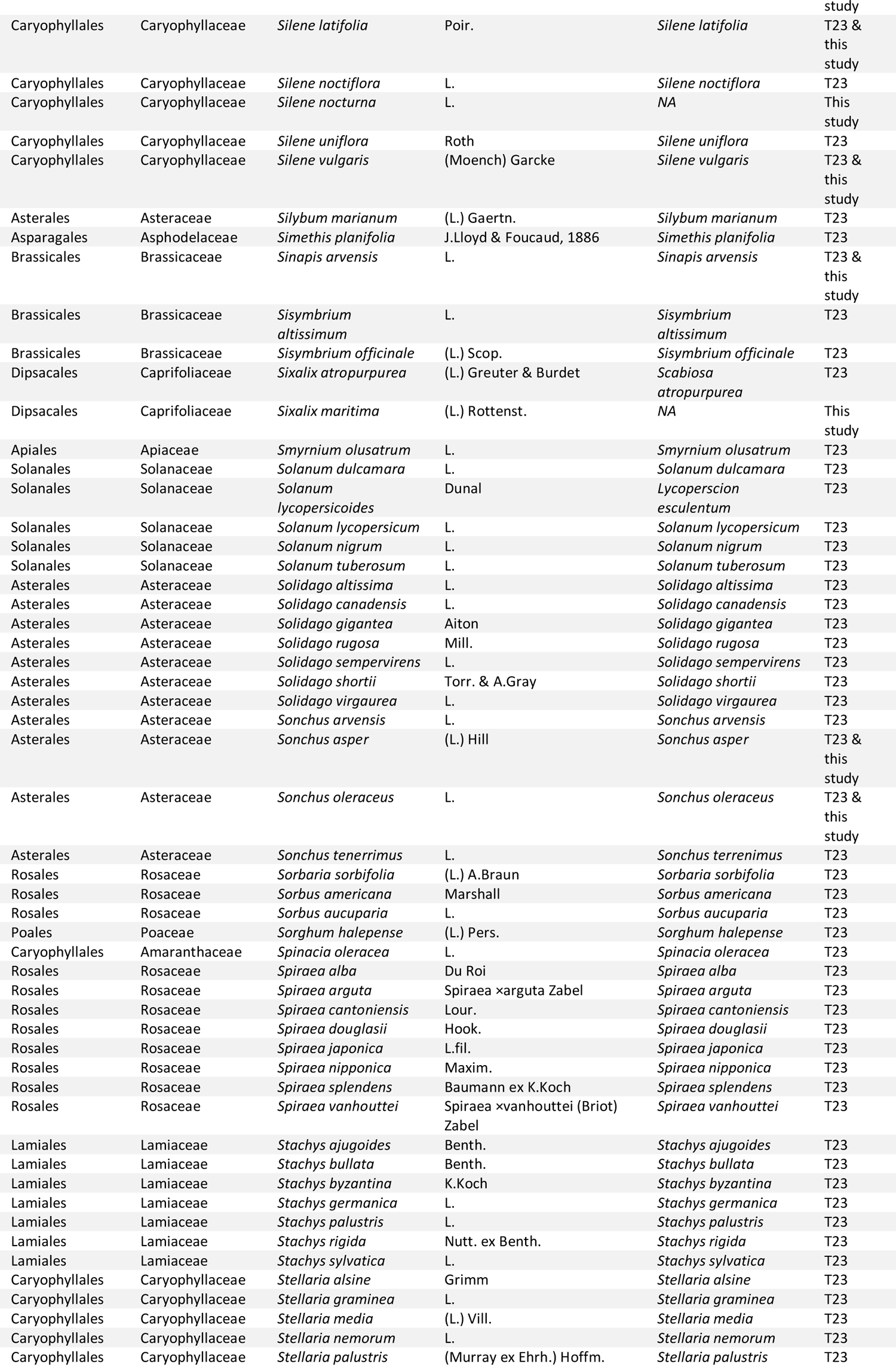
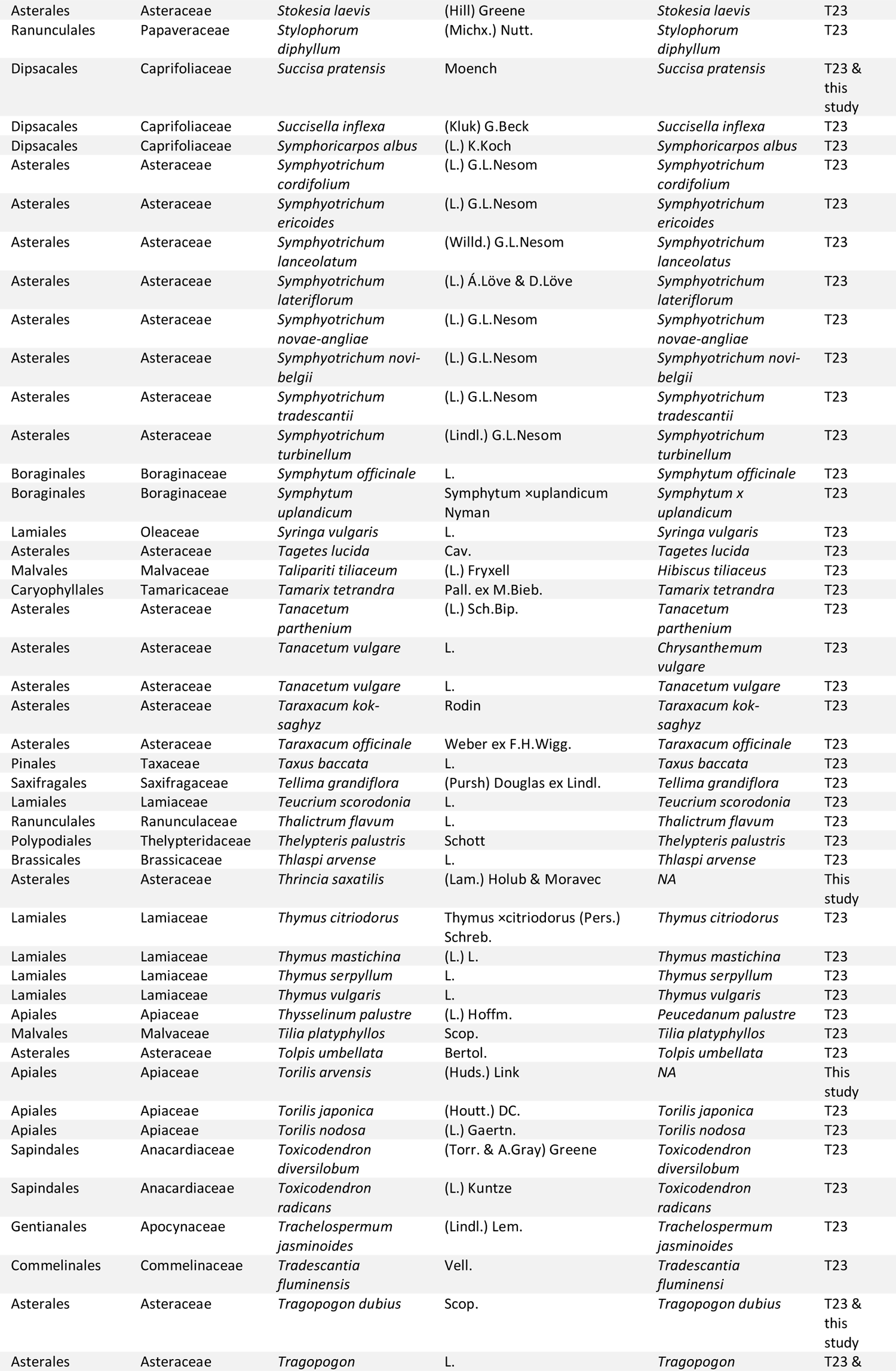
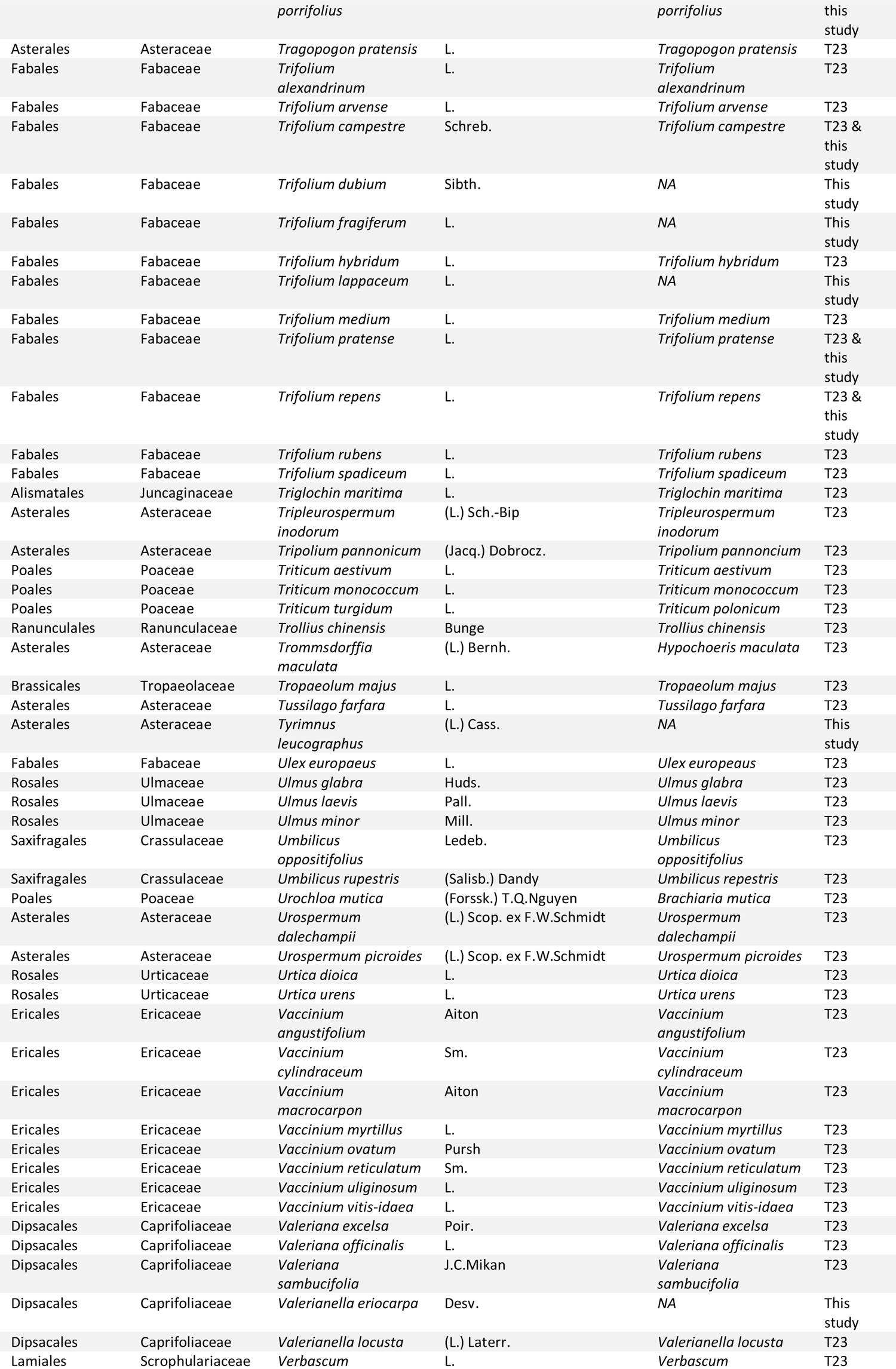
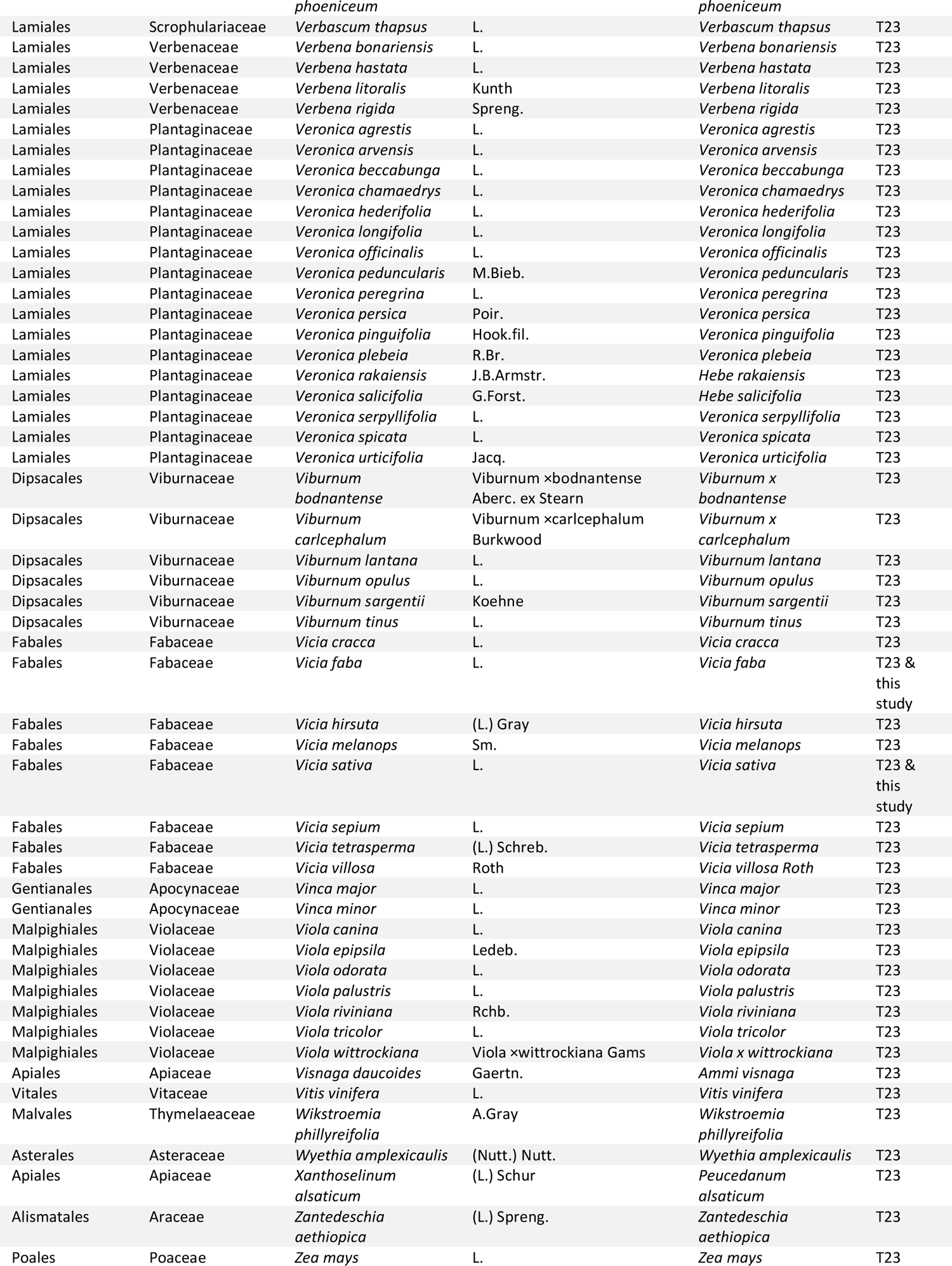
Plant species listed in Thompson et al. (2023) and in the present study as hosts of *P. spumarius*. Taxonomy follows the currently accepted names in GBIF (GBIF.org, 2023) after check with ‘get_gbif_taxonomy’ function in R (Schneider, 2018). “T23“ in the “Source” column stands for “Already reported (Thompson et al., 2023)“.

